# Autism and Williams syndrome: dissimilar socio-cognitive profiles with similar patterns of abnormal gene expression in the blood

**DOI:** 10.1101/2020.03.15.992479

**Authors:** Amy Niego, Antonio Benítez-Burraco

## Abstract

Autism Spectrum Disorders (ASD) and Williams Syndrome (WS) exhibit quite opposite features in the social domain, but also share some common underlying behavioral and cognitive deficits. It is not clear, however, which genes account for the attested differences (and similarities) in the socio-cognitive domain. In this paper we adopted a comparative-molecular approach and looked for genes that might be differentially (or similarly) regulated in the blood of people with these two conditions. We found a significant overlap between differentially-expressed genes compared to neurotypical controls, with most of them exhibiting a similar trend in both conditions, but with genes being more dysregulated in WS than in ASD. These genes are involved in aspects of brain development and function (particularly, dendritogenesis) and are expressed in brain areas (particularly, the cerebellum, the thalamus and the striatum) of relevance for the ASD and the WS etiopathogenesis.

## Introduction

Cognitive disorders usually exhibit complex phenotypical profiles. In cases with an unknown molecular etiology, as in autism spectrum disorders (ASD), many genes (and many variants of specific genes) have been found to contribute to the cognitive and behavioral problems exhibited by affected people, with each of them conferring low risk to the disease (Geschwind and State, 2015; Gyawali and Patra, 2019). In cases with a known etiology, like in Williams syndrome (WS), robust gene-to-phenotype correlations are also difficult to establish, particularly, for cognitive and behavioral deficits (see Korenberg et al., 2000; Tassabehji, 2003; Karmiloff-Smith et al., 2012; Ghaffari et al., 2018 among others for discussion), seemingly because these problems result in most cases from the dysregulation of several other genes outside the affected genomic regions (e.g. Lalli et al., 2016 or Kimura et al., 2019 for WS). Consequently, for these clinical conditions, it is of particular interest to examine the expression pattern of genes across the whole genome. This approach ultimately follows the ‘omnigenic’ theories of complex diseases, according to which such diseases result from the altered expression of most of the genes expressed in the affected tissues, with many of them located well outside the core pathways leading to disease (Boyle et al., 2017; Peedicayil and Grayson, 2018a,b). Still, even if hundreds of genes are found to be dysregulated in patients, specific pathways or specific biological processes are expected to be differentially affected in different conditions, enabling the identification of intermediate, disease-specific phenotypes (e.g. abnormal, disease-specific gene expression profiles). This should help clarify the genetics of these conditions and the clinical symptoms observed in affected people, as well as achieve earlier and more precise diagnoses. For instance, several co-expression modules of genes outside the region deleted in WS are found to be dysregulated in the blood of subjects with this condition and enriched in genes related to RNA processing and RNA transport (Kimura et al., 2019). Eventually, traits that appear to be omnigenic in lesser studies seem to have finite genetic determinants (Jakobson and Jarosz, 2019). Consequently, more studies of this sort are needed if we want to achieve robust conclusions about the etiology of complex cognitive disorders.

At the same time, it has been argued that a promising way of bridging the gap between the genome and the phenotype in these conditions is to adopt a comparative approach, instead of focusing on each disorder separately. For instance, ASD and schizophrenia (SZ) exhibit distinctly contrasting features, from neurodevelopmental pathways to brain structure and function to cognitive (dis)abilities, including language (see Crespi and Badcock 2008, Murphy and Benítez-Burraco, 2017 among others for discussion). As far as gene expression profiles are concerned, there is also some evidence of diametric gene-dosage in some cases (Byars et al. 2014). Thus it can be hypothesized that SZ and ASD might share the same genetic determinants, but with some key genes exhibiting opposite patterns of abnormal down- or upregulation compared to controls. The development of next-generation sequencing facilities and the analyses of thousands of cases by large consortia have exponentially increased the number of available genetic variants. These suggest that common biological mechanisms can be in fact implicated in both SZ and ASD in spite of their distinct clinical profiles and onset times, mostly converging on aberrant synaptic plasticity and remodeling, and ultimately, on altered connectivity between brain regions (Liu et al., 2017). Accordingly, dosage-sensitive gene expression emerges as a key etiological factor of complex diseases. This is reinforced by the finding that copy number variations (CNVs) in the human genome impacting on the same genes are a risk factor for different psychiatric disorders, especially SZ and ASD (Zarrei et al., 2019). In some cases, mechanistic insights can be provided. For instance, altered excitatory/inhibitory balance is implicated in both SZ and ASD, seemingly accounting for many of their distinctive cognitive features, including language deficits (see Murphy and Benítez-Burraco, 2017 for discussion). CNVs in the gene *CYFIP1* have been associated to both conditions, with *CYFIP1* upregulation increasing excitatory synapse and decreasing inhibitory synapses and with *CYFIP1* knockout resulting in synaptic inhibition (Davenport et al., 2019).

In this paper, we have adopted a comparative approach to cognitive disease with the aim of illuminating aspects of the problems that people with ASD and WS experience with social cognition and social behavior; two aspects which exhibit quite opposite features in many domains. Also in line with our discussion above, we have further adopted a molecular comparative approach. Accordingly, we have relied on the abnormal transcriptional profiles in the blood of affected people to identify genes that can be similarly or dissimilarly dysregulated between conditions and that can account for some of the similarities and the differences observed at the cognitive and behavioral levels. Certainly, as far as cognitive conditions are concerned, the most important changes are expected to occur in the brain. However, because blood and brain transcriptional profiles exhibit a notable overlap, ranging from 20% (Rollins et al., 2010) to 55% (Witt et al., 2013), blood expression profiles can be employed to infer changes of relevance for the etiopathogensis of these conditions (see Bjørklund et al., 2018 or Shen et al., 2019 for ASD). The paper is structured as follows. First, we provide a comparative characterization of the socio-cognitive and socio-behavioral profiles of people with ASD and WS. Afterwards, we discuss several neurobiological hypotheses aimed to account for the observed deficits and strengths in the social domain. Then, because much less is known about the genes accounting for these problems, we have determined the genes that are abnormally expressed in the blood of people with ASD or WS compared to the neurotypical controls. We discuss the biological roles played by the genes that exhibit similar abnormal expression patterns in both conditions, as well as the role of those showing opposite expression trends. We conclude with some reflections about our findings and more generally, about the utility of our approach for achieving a better understanding of the etiology of these two conditions.

## Materials and Methods

### Comparative characterization of the socio-cognitive profile of patients

For the comparisions of the socio-cognitive deficits exhibited by subjects with ASD and WS, as well as for the discussion around their neurobiological basis, we relied on available repositories of technical papers, particularly, PubMed (https://pubmed.ncbi.nlm.nih.gov/). Whenever possible, we made use of metaanalyses and review papers.

### Gene expression profiles in the blood of patients

For determining the genes that are differentially expressed in the blood of subjects with ASD or WS, we analyzed publicly available Gene Expression Omnibus (GEO) dataset (accession number: GSE 89594). The GSE 89594 dataset was consisted of 32 patients with ASD (mean age 24.0 years, male/female ratio 50:50), 32 patients with WS (mean age 21.6 years, male/female ratio 50:50), and 30 controls (mean age 23.9 years, male/female ratio 50:50) (Kimura et al., 2019). These data were obtained with Agilent SurePrint G3 Human GE v2 8×60K microarray (Agilent Technologies) from peripheral blood samples (all samples had RNA integrity number (RIN) values over 8). Differentially expressed genes (DEGs) were calculated based on diagnosis, age, gender, and RIN using the Limma R package (Smyth, 2005). Genes were considered to be differentially expressed when the false discovery rate (FDR) <0.1 and the |fold change (FC)| > 1.2. The Benjamini-Hochberg procedure was used for controlling the FDR in multiple testing (Benjamini and Hochberg, 1995). All the human protein-coding genes were considered and 17446 genes were regarded as background. The list of DEGs in subjects with ASD compared to controls encompasses 242 genes (Supplemental file 1; column A). The list of DEGs in subjects with WS compared to controls encompasses 882 genes (Supplemental file 1; column B). A hypergeometric test was used to determine the significance of the overlapping DEGs between the two clinical conditions.

### Functional characterization of genes of interest

For providing a detailed characterization of the functions performed by DEGs, we compiled information about their association with ASD and/or WS, their involvement in comorbid conditions, and/or their role in physiological aspects of relevance (mostly at the brain level) for the etiopathogenesis of the socio-cognitive dysfunctions observed in ASD and/or WS. For achieving this, we checked the available literature via PubMed (ncbi.nlm.nih.gov/pubmed), but also relied on other common gene databases, particularly GeneCards (https://www.genecards.org/).

### Gene ontology analsysis

Gene ontology (GO) analyses of the sets of DEGs were performed via Enrichr (amp.pharm.mssm.edu/Enrichr; Chen et al., 2013; Kuleshov et al., 2016). We considered biological processes, molecular functions, cellular components, or human pathological phenotypes as enriched if their p<0.05.

## Results

### 1. Contrasting the ASD and the WS socio-cognitive phenotypes

#### General overview

At first sight, ASD and WS can be viewed as opposite conditions, at least as far as social behavior is concerned. Individuals with ASD are normally characterized as withdrawn, difficult to engage in social interaction, ignorant of social norms, and generally uninterested in social relationships with others (for a general review, see Newschaffer et al., 2007). This hyposocial phenotype starkly contrasts with the hypersocial phenotype exhibited by people with WS, who are usually characterized as overly friendly, gregarious, and eager to interact with others, sometimes to an excessive degree (for a general review, see Bellugi et al., 2000; Jones et al. 2000, Doyle et al. 2004; Martens, et al. 2008; Järvinen et al. 2013). However, this is just a rough picture that deserves a closer examination, not just because such a close look reveals an intricate profile of similarities and differences, as we show in this section, but because these two conditions have been claimed to share genetic determinants (Newschaffer et al 2007, Jawaid et al 2012). Ultimately, social aspects and the management of the social context can be crucial for understanding the different clinical presentation of both conditions, considering that in terms of general cognition, ASD and WS are much more similar when social motivation is removed. Hence, Vivanti and colleagues (2016) found that when the objective of a task is learning and not social interaction, both ASD and WS subjects are equally able to imitate and learn in social situations (see Ingersoll et al., 2013; Berger and Ingersoll, 2015 for similar findings). Eventually, exploring these overlaps and divergences is expected to reveal more about the etiology of atypical social behavior in humans and about the evolution of human social cognition. Our findings are summarized in Figure 1 and are discussed in more depth below.

**Figure 1.**
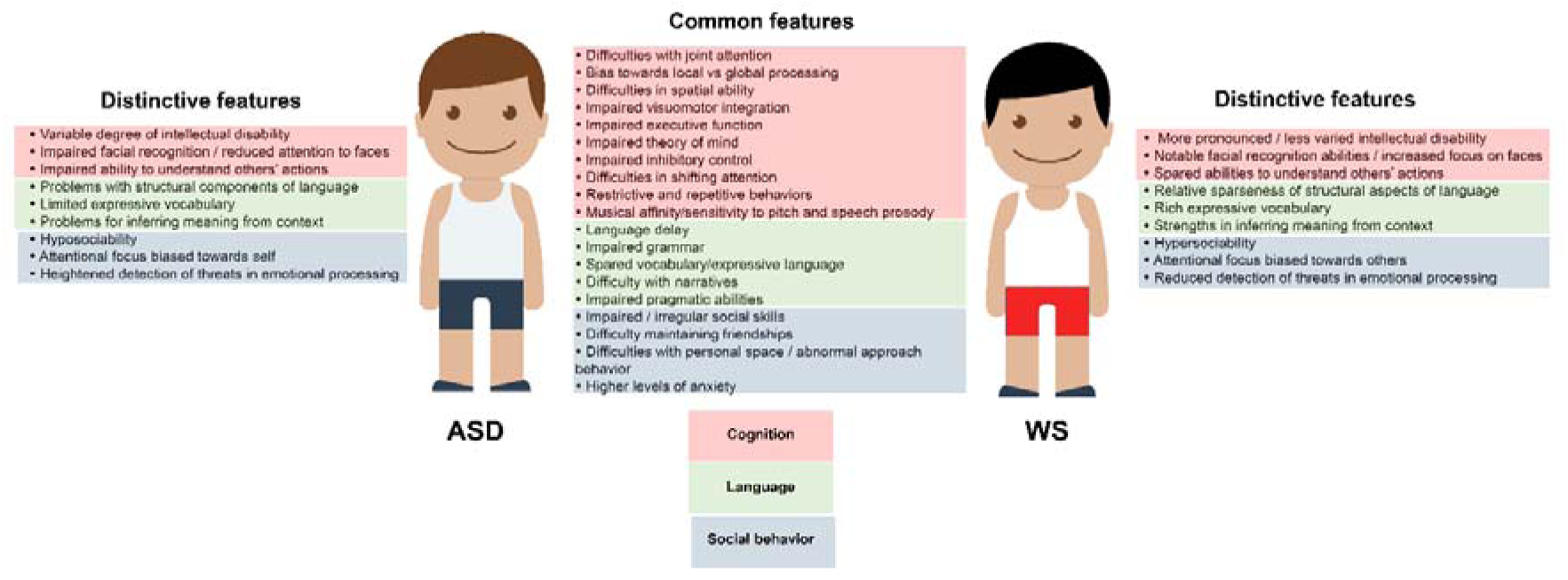
An overview of the socio-cognitive differences between ASD and WS.

#### Socio-cognitive similarities between ASD and WS

In spite of people with WS being labeled as ‘overly friendly’ and ‘hypersocial’, they exhibit many difficulties in the social arena that overlap in part with ASD. Accordingly, parents of children with WS often report that they have poor social skills, difficulties with understanding important social cues or information, and difficulty maintaining friendships (Mervis et al., 2001; Sullivan et al., 2003; Stojanovik, 2006; Klein-Tasman et al., 2009; Jarvinen et al., 2015). Included in these similarities are “…social isolation, and other types of social impairment, distractibility, inflexibility, ritualism, obsessiveness, and pragmatic deficits” (Gillberg and Rasmussen, 1994). Both groups also show difficulties with the typical boundaries of personal space (Lough et al., 2015). These shared social difficulties make them more vulnerable socially, prone to disadvantages when it comes to forming productive relationships and social connections. Social vulnerability is thought to be the source of some of the anxiety documented in both groups, which report higher levels compared to neurotypical people (Dykens, 2003; Graham et al., 2005; Jawaid et al., 2012). This elevated anxiety is coupled in many cases with restricted and repetitive behaviors (Rodgers et al., 2012).

To an important extent, these problems in the social sphere are expected to arise from deeper cognitive and behavioral dysfunction. Young children with ASD or WS show delays in pointing behaviors and joint attention, and underperform in theory of mind (ToM) tasks (i.e. tasks aimed to evaluate the subject’s ability to attribute mental states to others), like in the false belief test (Baron Cohen et al., 1985; Charman et al., 1997; Klein-Tasman et al., 2009; Vivanti et al., 2016). Also documented are difficulties in inhibitory control and shifting attention (Rhodes et al. 2010; Riby et al., 2011). Sparaci and colleagues (2014) found that both participants with ASD and WS had difficulty specifying the ‘why’ of an experimenter’s actions. That is, in a task where they were asked to observe a hand-object action (e.g. grasping a mug to drink tea or touching the handle to put it in a cupboard), both groups had difficulty specifying why they were doing so (i.e. they couldn’t say they were grasping the object to put it away). In contrast to their similarities in ‘why understanding’, participants with ASD showed superior ability in specifying the ‘what’ of an experimenter’s actions. That is, they were better at specifying that the individual was grasping a mug vs. simply touching it, while those with WS showed more difficulty with this task. As Sparaci and colleagues (2015) point out, this difference in ‘what’ ability coupled with a similarity in ‘why’ ability is indicative of the fact that certain low level abilities may shape higher level processes in unexpected ways (Sparaci et al., 2015). In a similar vein, the difficulty ASD and WS individuals have with predicting others’ intentions and actions have been hypothesized to arise from deficits in predicting physical actions. Indeed, motor impairments have been widely documented in both ASD (e.g. Teitelbaum et al., 1998; Jansiewicz et al., 2006; Dewey et al., 2007) and WS (see Trauner et al. 1989; Elliott et al. 2006; Gagliardi et al. 2007).

Finally, the socio-cognitive problems exhibited by people with ASD or WS are expected to have an impact on (and result in part from) their language deficits. Language development is generally delayed in both conditions, although it tends to follow the typical progression (Asada and Itakura, 2012). In both ASD and WS, grammar was originally thought to be relatively spared, although more recent research has highlighted the fact that it is more impaired than in appears at the surface (see Perovic et al. 2013 and Lacroix et al., 2016 for direct comparisons). In both WS and ASD, grammatical impairment has been shown to happen with aspects of grammar defined later in typical development, like raising and passives (Tager Flusberg 1981; Perovic et al., 2007; Perovic and Wexler, 2007), grammatical morphology (Kjelgaard and Tager –Flusberg, 2001; Roberts et al., 2004), relative clauses (Riches et al., 2009), and subject-control structures (Perovic and Janke, 2013). Importantly for our concerns here, pragmatics (that is, the ability to use language for communicating in a social context) is impaired in both conditions (see Tager-Flusberg, 2000; Doyle et al., 2004: Laws and Bishop, 2004; Stojanovik, 2006; Philofsky et al., 2007; Jarvinen-Pasley et al., 2008; Asada and Itakura, 2012; Lacroix et al. 2016 among others), although, as we will show in the next section, it is more so in ASD. Specifically, both ASD and WS individuals have been rated as impaired in the quality of their conversational initiations with others, with WS subjects underperforming people with ASD (Philofsky et al., 2007). Likewise, narration, a basic pragmatic skill, has also been shown to be impaired in both conditions. Accordingly, Diez-Itza and Martinez (2016) reported in their study that WS individuals often give the impression of having in-tact narrative skills because of their overuse of discourse markers and exclamations, but in-depth analyses further reveal deficiencies in sequencing narratives. Similar deficits in sequencing narratives have been reported in ASD (see Freeman and Dake, 1996; Happé and Frith, 1996).

Studies have shown a direct correlation between socio-cognitive skills and deficits and pragmatic language (dis)abilities in both WS and ASD (Happé, 1993; Surian et al., 1996; Hale and Tager-Flusberg, 2005; John et al., 2009), suggesting that deficits in the social domain impact negatively on language use in both conditions. Difficulties with pragmatics can be hypothesized to derive from specific deficits that both groups have, particularly, problems with inferring the mental states of others (Asada and Itakura, 2012). Likewise, problems with narration, and specifically, the lack of ability to sequence events, might stem from difficulties in spatial cognition, at least in the case of WS (Phillips et al., 2004). At the same time, socio-cognitive deficits impacting language use certainly contribute to language delay in both conditions, as many aspects of language acquisition require inference of the speaker’s intentions (Preissler and Carey, 2005). Other probable factors contributing to language delay are deficits in joint attention, which especially impact the acquisition of vocabulary and semantic meaning of words (Baron Cohen et al., 1985; Charman et al., 1997; Klein Tasman et al., 2007; Mervis and Becerra, 2007; Mervis and John, 2012). At the same time, as noted, there are aspects of language that appear similarly preserved in both conditions in spite of socio-cognitive impairment. For instance, in their study on grammatical binding, Perovic and colleagues (2013) came to the conclusion that the impairment of grammatical knowledge on binding in both conditions is independent of difficulties in social interactions and pragmatics (Perovic et al., 2013).

#### Socio-cognitive differences between ASD and WS

Although, as discussed above, similarities between ASD and WS can indeed be found and while the social profiles of both conditions are in no way homogeneous, research consistently shows that those with WS have less socio communicative deficits than those with ASD do (e.g. Bellugi et al., 2000; Lincoln et al., 2007; Klein-Tasman et al., 2009; Lacroix et al., 2016). In general, people with WS and ASD have been shown to have opposite preferences in terms of their orientation to social vs. nonsocial information, as reported by eye tracking studies such as Riby and Hancock (2008, 2009). More specifically, in social situations, individuals with ASD tend to direct their attention inward towards themselves and prioritize information related to them as individuals, while those with WS direct their attention to others. As pointed out by Kuang (2016) in a social setting, neurotypical people rely on both systems of attention (attention to self and attention to others) in order to interact properly. This increased attention towards others may in turn help individuals with WS to preserve more emotional empathy than those with ASD can, in spite of the fact that, as discussed previously, both exhibit difficulties with imagining mental states of others (Tager-Flusberg and Sullivan, 2000). As a consequence, individuals with WS are eager to engage socially and are highly motivated to approach familiar and unfamiliar people (Bellugi et al., 2000). In contrast, people with ASD attend much less to socially salient features and are generally reluctant to engage with others, regardless of whether they are familiar or not (Sigman et al., 2006; Riby and Hancock, 2009). Specifically, ASD and WS show distinct differences in the area of face recognition skills; people with WS are hyper-atentive to faces and reportedly perform better than mental age matched controls on standardized tests of face recognition skills, while those with ASD attend much less to faces and perform distinctly worse (Bellugi et al 1994; Kiln et al., 1999; Schultz, 2005; Tager Flusberg et al., 2006; Rose et al., 2007). The results vary, however, when emotional recognition is involved, as people with WS have more difficulty recognizing emotions from facial expressions than those with ASD do (Lacroix et al., 2009). Interestingly, the results are the opposite when subjects are asked to analyze emotion in music, with individuals with WS outperforming those with ASD (Bhatara et al., 2010), in spite of both conditions exhibiting similar affinity and interest towards music in general (Heaton et al., 1998; Bonnel et al., 2003; Heaton, 2003; Bhatara et al., 2010). In truth, enhanced musical abilities of people with WS do not concern the structural aspects of music, but instead are related to musicality and expressivity, commonly expressed through a heightened emotional responsiveness to music (Thakur et al., 2018).

Both conditions contrast as well in the management of anxiety in social settings. In both WS and ASD levels of anxiety correlate with their degree of social impairment, although those with ASD are thought to have higher levels of anxiety in general (Rodgers et al., 2012). Higher levels of restricted and repetitive behaviors correlated to higher levels of anxiety in ASD but not WS (Rodgers et al., 2012), which suggests that these behaviors may serve different functions in both conditions. Barak and Feng (2016) posited that these differences in anxiety levels correlate to social cognition, that is, it may be that the social drive of WS acts as a buffer against social anxiety, while in ASD social impairments might make them more vulnerable to anxiety (see also Frigeria et al., 2006). White et al., 2010).

Regarding language, and particularly, language use in social settings, differences between conditions can be observed as well. As noted above, generally individuals with ASD show a more marked overall deficit in communicative use of language, while those with WS show certain strengths including ample, descriptive vocabularies, often with elaborate attention to detail and expressive phrases that are full of emotion and affect (Udwin and Yule, 1990; Bellugi et al., 1994; Bellugi et al., 2000; Reilly et al 2004; Brock et al., 2007; Gothelf et al., 2008; Fishman et al., 2011;). Still, there are indications that the processes involved in acquiring and using these characteristics are irregular, as can be seen from research exploring atypical activation of semantic networks after pointing (Lukács et al., 2004). By contrast, in ASD, vocabulary progresses steadily with age, but it contains a disproportionately high number of nouns and a much lower number of mental state terms when compared to typically developing children (Fein et al., 1996; Gastgeb et al., 2006; Kelley et al., 2006; Swensen et al., 2007; Tek et al., 2008). Regarding the access to implicatures (i.e. non-explicit meaning of utterances), Fishman and colleagues (2011) found that individuals with WS show larger N400 effects (N400 is an ERP component which is inversely correlated with the semantic fit of a word or phrase), suggesting that they do rely on context to infer meaning. This contrasts with the significantly smaller N400 effect found in individuals with ASD, which suggests that they make less use of contextual information when it comes to language use (Fishman et al., 2011). Other studies (e.g. Firth and Snowling, 1983; Happé 1997; Tager-Flusberg, 2003 and 2004; Harris et al., 2006; Walenski et al., 2006,) also document difficulties with comprehension of meaning from context in ASD; even individuals with high-functioning ASD show irregularities in an otherwise typical IQ profile, with significantly lower scores in comprehension tasks, such as comprehending idioms (Siegel et al., 1996; Goldstein et al., 2002). Fishman et al. (2011) highlight as well the fact that ASD individuals have been shown to rely on visual imagery instead of linguistic cues to comprehend sentences (Kana et al., 2006), while people with WS tend to rely on sentence-level context cues. Pragmatic problems, and differences between conditions, have been documented in other studies. For instance, in their comparative study of the pragmatic language profiles of children with ASD and WS, Philofsky and colleagues (2007) found that in certain areas (coherence, stereotyped language, nonverbal communication, and social relations scales) individuals with WS performed better than subjects with ASD, although in other areas (inappropriate initiation, use of context, and interests scales) the impairment was similar. Also, WS children were rated by their parents as being slightly better than ASD individuals at communicative tasks like appropriately sequencing and referencing events for a listener, but they still had difficulties (Philofsky et al., 2007). As expected, these differences in language use between ASD and WS can be attributed to their divergent socio-cognitive profiles: after all, language is learned and used in social situations, and attention to the facial area and emotional state of the speaker, as well as interest in the interlocutor play a crucial role at these levels. Hence, Fishman and colleagues (2011) suggest that the different patterns of attention exhibited by people with ASD and subjects with WS may lead to different perceptual inputs, which in turn lead to different communicative behaviors. Likewise, the decreased sensitivity to speech prosody documented in ASD (e.g. Korpilahti et al., 2007) could explain their impaired ability to infer meaning from context, whereas the auditory hypersensitivity documented in WS (e.g. Klein et al., 1990; Blomberg et al., 2006) could account for their enhanced ability at this level (see Fishman et al. 2011 for discussion).

#### 2. Contrasting the (neuro)biological causes of the ASD and the WS abnormal socio-cognitive phenotypes

In studies of both ASD and WS, a wealth of research has focused on specific brain networks thought to be implicated in behavioral and cognitive abnormalities of both conditions. Here we provide a brief overlook of research focused on the social realm. Such networks provide a more biologically-grounded account of the similarities and differences in the socio-cognitive phenotypes of these conditions and are expected to facilitate the formulation of bridging theories linking these phenotypes to the underlying genetic factors (our focus of interest in the next section of the paper). An important outcome of the neurobiological characterization of ASD and WS is that similar irregularities in brain structures and their functional connectivity do not always translate to similar behaviors. As research repeatedly shows, what may look like a similar impairment at the neurobiological level can translate to a completely different trait at the phenotypic level. Overall, studies of WS and ASD consistently provide additional evidence of the brain’s flexibility and its ability to adapt its connections and functions when irregularities and deficits arise.

#### Socio-cognitive similarities in ASD and WS: explanatory hypotheses

Regarding the associations between irregularities in certain brain networks and some core social deficits found in both ASD and WS, four networks are worth considering. The first is the default mode network, associated with various elements of social cognition, which includes the medial prefrontal cortex (mPFC), the posterior cingulate cortex (PCC), the precuneus, the inferior parietal lobes (IPL), and medial temporal regions; its functional connectivity has been found to be irregular in ASD (Assaf et al., 2010; Lynch et al., 2013) as well as WS (Sampaio et al., 2016). Even more relevant is the second network, namely, the social brain network, which includes the superior temporal sulcus (STS), the anterior cingulate cortex (ACC), the medial prefrontal cortex (mPFC), the inferior frontal gyrus (IFG), and the anterior insula and the amygdala. Research has uncovered atypical connections and irregularities in this circuitry in both ASD (Gotts et al., 2012; Kennedy and Adolphs, 2012) and WS (Barak and Feng, 2016). Also of interest is the circuitry involved in self-representation, which includes the mPFC, the PCC/precuneus, the temporo-parietal junction (TPJ), the anterior insula, the middle cingulate cortex (mCC), the ventral premotor cortex (PMv), and the somatosensory cortex, which have also been deemed atypical in ASD (Lombardo et al., 2010) and WS (Haas et al., 2013). Specifically, in their voxel-level study of resting state functional connectivity in ASD, Cheng and colleagues (2015) found that the medial temporal gyrus exhibits reduced cortical connectivity and increased connectivity to the medial thalamus in ASD participants, and posited that this may be related to face processing deficits and ToM impairments (Cheng et al., 2015). Cheng and colleagues (2015) also found in people with ASD a key system in the precuneus/superior parietal lobe with reduced functional connectivity, which is implicated in spatial functions, including those related to self and the environment. These elements are substrates of ToM, so it stands to reason that reduced connectivity in these regions may help to explain key elements in the social phenotype of ASD. In studies of WS, Sampaio and colleagues. (2016) also found decreased functional connectivity in the precuneus, as well as the posterior cingulate of the left hemisphere, which is also implicated in the default mode network. Finally, also of interest is the reward circuitry, including the ventral tegmental area (VTA), the striatum, the orbitofrontal cortex (OFC), the ventromedial prefrontal cortex (vmPFC) and the ACC, which have been shown to be irregular in ASD and WS (Dichter et al., 2012).

Special attention regarding the similarities between both conditions has been paid to two particular brain structures mentioned above: the amygdala and the frontal lobes. Although differences are attenuated as subjects get older (Martens et al., 2009), the amygdala is disproportionately large in both ASD and WS (Reiss et al., 2004; Schumann et al., 2004; Martens et al., 2009; Mosconi et al., 2009; Haas et al., 2012; Murphy et al., 2012; Jarvinen et al., 2013; Gibbard et al., 2018). The amygdala shows atypical functional connections to several brain regions, specifically with the ACC, the PFC, and the OFC (all of which are implicated in cognitive processing, attention, and inhibition) (Martens et al., 2008; Dedovic et al., 2009; Haas et al., 2014; Gibbard et al., 2018), and importantly, with various components of the social brain, particularly, the frontal lobes (Meyer-Lindenberg, 2005; Paul et al., 2010; Jawaid et al., 2012). A key component of the limbic system, the amygdala is a set of brain structures that support emotion and motivation, among other functions (see Rolls, 2015 for review), and is forefront in much of the research about the social (dys)function in both conditions (Stefanacci and Amaral, 2000; Meyer-Lindenberg et al., 2005; Haas et al., 2010; Paul et al., 2010; Jawaid et al., 2012; Zalla and Sperduti, 2013; Barak and Feng, 2016). The amygdala is also instrumental in the anxiety reported in both ASD and WS; a common source of this anxiety has been found to be irregular hyper- and hypo-activation of the amygdala in response to both social and non-social stimuli in both conditions (see Reiss et al., 2004; Martens et al., 2009; Jarvinen et al., 2013 for WS; see Baron-Cohen and Wheelwright, 1999; Critchley et al., 2000; Dalton et al., 2005; Corbett et al., 2009; Kliemann et al., 2012; for ASD). Specifically, Barak and Feng (2016) highlight that the amygdala is the source of the non-social anxiety and phobias which are typical of WS, pointing to deficits in the prefrontal-amygdala white matter pathways as the cause (see also Avery et al., 2012). Research on a salience network including the amygdala, the ventral striatum, the dorsomedial thalamus, the hypothalamus, and the substantia nigra (SN)/VTA has shown these functional connections to be atypical in ASD (Uddin et al., 2013) as well as in WS (Haas and Reiss, 2013). This network has been linked to attention switching, as well as detection and attention to sensory and emotional stimuli. Regarding the frontal areas, it should be noted that individuals with WS exhibit similar approach behavior to people with frontal lobe damage, suggesting that this abnormal approach behavior could be due to a lack of inhibitory control in the frontal lobe (Porter et al., 2007). Specifically, abnormal functional connectivity between the OFC and the amygdala has been linked to the uninhibited social nature in WS, since the frontal lobes have been shown to regulate and inhibit inappropriate social behavior (Meyer-Lindberg et al., 2005; Mobbs et al., 2007; Porter et al., 2007; Little et al., 2013; Barak and Feng, 2016). Similar lack of inhibitory control has also been shown in people with ASD, but in terms of abnormal personal space boundaries (Christ et al., 2007). Moreover, language deficits and language delay have been associated to a frontal lobe dysfunction and irregular functional connectivity to the amygdala, seemingly impacting on aspects like inferencing or joint attention (Lincoln et al., 2002; Cornish et al., 2007; Martens et al., 2008; Barak and Feng, 2016).

Research has also implicated the mirror neuron system (MNS) in social dysfunction in ASD and WS (e.g. Jarvinen et al., 2013). This network includes the frontal gyrus, the STS, and the IPL (Van Overwalle and Baetens, 2009). Apart from its role in imitation, decoding, and implementation of actions (see e.g. Rizzolatti and Craighero, 2004), the MNS is also related to the social realm in terms of empathy (e.g. Gallese, 2001; Iacoboni, 2009). Reduced cortical surface area, but preserved cortical thickness, in structures implicated in the MNS have been found in WS (Ng et al., 2016). Studies of the MNS in ASD report cortical thinning in selected areas, which positively correlate to degree of social dysfunction (Hadjikhani et al., 2006; Wallace et al., 2012). These findings indicate that the common social deficits found in both WS and ASD may stem from an atypical MNS, while the distinct social drive in WS is most likely derived from systems independent of this network (Ng et al., 2016).

Finally, the HPA axis (a major neuroendocrine system resulting from the interaction between the hypothalamus, the pituitary gland, and the adrenal glands) comes into play here, because of its involvement in stress response---particularly, the response to cortisol in the amygdala, the PFC, and the hippocampus--all of them areas implicated in situations of fear and social stress (see Martens et al, 2008; Dedovic et al., 2009; Lense and Dykens, 2013; Bitsika et al., 2015). Both ASD and WS individuals have been shown to exhibit interrupted HPA axis function (Spratt et al., 2012; Jacobson, 2014; Benítez-Burraco et al., 2016; Niego and Benítez-Burraco, 2019, among many others). This might explain in part the prevalence of anxiety in the two disorders, although as mentioned above, the anxiety generally seems to happen in separate arenas for each group: social in ASD and non-social in WS (see also Dykens, 2003; Graham et al. 2005; Rodgers et al., 2012; Lense and Dykens, 2013).

#### Socio-cognitive differences in ASD and WS: explanatory hypotheses

Just as the observed similarities in their respective social phenotypes happen to follow from similar dysfunctions in similar brain areas and circuits, the differences between ASD and WS in the realm of social behavior and social cognition stem from the impairment of different devices. One focus of attention has been the amygdala. As noted above, ASD and WS share similar irregularities in the amygdala. Nonetheless, some of the differences between these two conditions in the social domain can be hypothesized to lie in certain sub circuits of the amygdala, its differential response to stimuli, and/or its different connections to brain regions upstream or downstream from it. For instance, during eye gazing, a positive activation of the amygdala has been reported in participants with WS while aversive activation was reported in people with ASD (Barak and Feng, 2016). An interesting point about this activation is that, according to Barak and Feng (2016), in both cases the amygdala is hyperactivated. However, this hyperactivation takes on different forms in the two conditions. In ASD, it seems that this hyper activation is negatively valenced, creating an aversive response in those with ASD. In contrast, those with WS experience an appetitive response from this hyperactivation, making them more apt to continue the eye gaze (Riby et al, Barak and Feng, 2016). In terms of processing faces, there seems to be a difference at the level of responsiveness of the amygdala; it is hyper responsive to unfamiliar faces in ASD and hypo responsive to the same stimulus in WS (Lough et al., 2015). This might explain their distinctive response to faces and eye gaze. Accordingly, as mentioned above, the the aversive response triggered by hyperactivation of the amygdala would cause an aversive response to faces, resulting in a reduced sustained attention to the facial region, and hence, problems for facial recognition (Dalton, 2005; Kliemann et al., 2012; Strauss et al., 2009). The opposite may happen in WS: the appetitive response triggered by amygdala hyperactivation may be a motivating factor to spend more time assessing facial features (Riby et al., 2009; Barak and Feng, 2016). It has also been shown that individuals with WS have a heightened amygdala reaction to images depicting non-social fear, but a somewhat muted amygdala response to fearful social images and faces. In contrast, subjects with ASD seem to present a negative over arousal of the amygdala when looking at faces (not necessarily fearful ones) which seems to follow the pattern of their social behavior (Meyer-Lindenberg et al., 2005; Haas et al., 2009; Mimura et al., 2010; Munoz et al., 2010; Barak and Feng, 2016). Eventually, one possible (complementary) explanation of all these differences is that the neurons within the amygdala responding to social stimuli come from different classes in each condition, e.g. glutamatergic or GABAergic (Barak and Feng, 2016). Still, it should be noted here that the results are far from straightforward and contrasting results have also been reported (e.g. Thornton-Wells et al., 2011).

Other brain areas have been implicated in the differences between ASD and WS in the socio-cognitive domain. These include the fusiform gyrus (Haxby et al., 2000), the pSTS (Allison et al., 2000; Nummenmaa and Calder, 2009), the amygdala (Adolphs and Spezio, 2006), and parietal-frontal areas such as the TPJ and the mPFC (Gallese and Goldman, 1998; Decety and Jackson, 2004; Amodio and Frith, 2006; Lieberman, 2007; Sui et al., 2013). Specifically, recent research indicates that the mPFC and the pSTS both have crucial roles in both sides of the attention spectrum, i.e., attention to self vs. attention to others, responding to each function with an activation response or an inhibition response: whereas the pSTS area is key in ‘attention to others’ functions, the mPFC seems to support the ‘attention to self’ function (Sui et al., 2013; Kuang, 2016). Not surprisingly, atypical connectivity and/or structures have been found in both the pSTS and the mPFC in both ASD and WS (Pelphrey et al., 2004; Amaral et al., 2008; Järvinen et al., 2013). Significant differences have also been found in the fusiform face area, an area in the fusiform gyrus involved in face processing. Accordingly, this area is twice the volume in individuals with WS than it is in typically developing individuals (Golarai et al., 2010, O’Hearn et al 2011; Haas and Reiss, 2012). In subjects with ASD, similar areas within the left fusiform gyrus have been shown to be hypo-activated during face processing (Nickl-Jockschat et al., 2015). Finally, differences in the structure and the connection patterns of the frontal lobes (particularly, with the amygdala) may contribute to distinctive features of the WS social phenotype, like sustained gaze towards faces, increased approachability perception for unfamiliar faces, difficulty in disengaging attention from faces, and difficulties with perceiving emotion from facial expressions (Bellugi et al., 1999; Porter et al., 2007). In contrast, it seems that altered connectivity with the frontal cortex in ASD is instrumental in decreased habituation of the amygdala response to emotional facial expressions, which correlates to their gaze aversion (Zalla and Sperduti, 2013; Swartz et al., 2013).

Finally, it is worth mentioning the role of oxytocin, a hormone known to regulate social behavior and interaction (Wojciak et al., 2012). Specifically, oxytocin inhibits the Hypothalamic–Pituitary–Adrenal (HPA) axis’ stress triggered activity (Neumann, 2002). Individuals with WS show an increased basal level of oxytocin, and levels of the hormone have been positively correlated with social engagement behaviors which are typical of this condition, such as tendency to approach strangers and emotionality (Dai et al., 2012). In contrast, lower levels of oxytocin have been reported in people with ASD (Modahl et al., 1998), with higher plasma concentrations of oxytocin correlating with enhanced verbal abilities (Zhang et al., 2016) and the retention of social information, like affective speech (Hollander et al., 2007). Administering oxytocin to individuals with ASD has been shown to help with attention to and retention of social cues, as well as promote eye contact (Hollander et al., 2007; Andari et al., 2010; Domes et al., 2013).

#### 3. Contrasting gene transcriptional profiles in the blood of subjects with ASD and WS

In contrast to what is known about the neurobiological causes of socio-cognitive dysfunctions in ASD and WS, less is known about the genetic factors that might contribute to the observed similarities and differences in the socio-cognitive domain, in spite of being conditions with a clear genetic basis. This is a consequence of the indirect link between genes, at the bottom, and behavior, at the surface, with many genes affecting a single phenotype (and vice versa), but more specifically, of the pervasive problems for mapping genes to complex phenotypes, which current approaches based on genome-wide association studies (GWAS) can only partially alleviate (Goddard et al., 2016, Guo et al., 2018). As noted in the Introduction, ASD and WS are no exception; the causes of ASD are not entirely known and no robust gene-to-phenotype correlations have been established for the cognitive features of WS. Still, as also noted, it is interesting that some genetic elements have been claimed to be shared between both conditions (Newschaffer et al 2007, Jawaid et al 2012). This circumstance raises the possibility that differences in gene dosage can account for some of the opposite features exhibited by affected people in the social cognition domain. An interesting instance is the gene *OXTR*, which encodes the oxytocin receptor. As noted in the previous section, individuals with WS exhibit increased basal levels of oxytocin correlating with social engagement behaviors. Higher levels of oxytocin in WS have been hypothesized to result from the hypomethylation (and thus, the overexpression) of *OXTR* (Haas and Reiss, 2015), seemingly as a result of the deletion of *WBSCR22*, which encodes a methyltransferase (Doll and Grzechik, 2001; Merla et al., 2002), and/or some effect of *GTF2I*, a gene also deleted in WS, which has proven to affect the reactivity to oxytocin and ultimately, sociability (Prosyshyn et al., 2017). By contrast, *OXTR* has been found to be hypermethylated (and thus, downregulated) in subjects with ASD, with this hypermethylation correlating with abnormal interconnection patterns between brain areas involved in the ASD pathogenesis, as well as with the severity of symptoms, including social cognitive deficits (Andari et al., 2020; see Maud et al., 2018 for review), particularly, with a decrease in the extent to which social information automatically captures attention (Puglia et al., 2018).

For all these reasons we conducted a comparative in vivo analysis aimed to uncover similar and dissimilar patterns of abnormal gene expression in the blood of subjects with ASD and WS. We found a very significant overlap (p = 7.93E-40) between DEGs in the blood of subjects with WS and DEGs in the blood of people with ASD. Interestingly, in spite of their opposite profile in many domains, particularly, in their socio-cognitive abilities, most DEGs exhibit a similar expression profile in the blood of ASD and WS patients, being either up-regulated or down-regulated in both conditions. Still, selected genes appear to be more strongly down- or upregulated in WS compared to ASD. Figure 2 shows the overlapping genes that are significantly upregulated in both conditions (n=18; Figure 2A), downregulated in both conditions (n=53; Figure 2B), upregulated in ASD but downregulated in WS (n=3; Figure 2C), and downregulated in ASD but upregulated in WS (n=1; Figure 2D) compared to controls (FDR <0.1, |FC| > 1.2).

**Figure 2.**
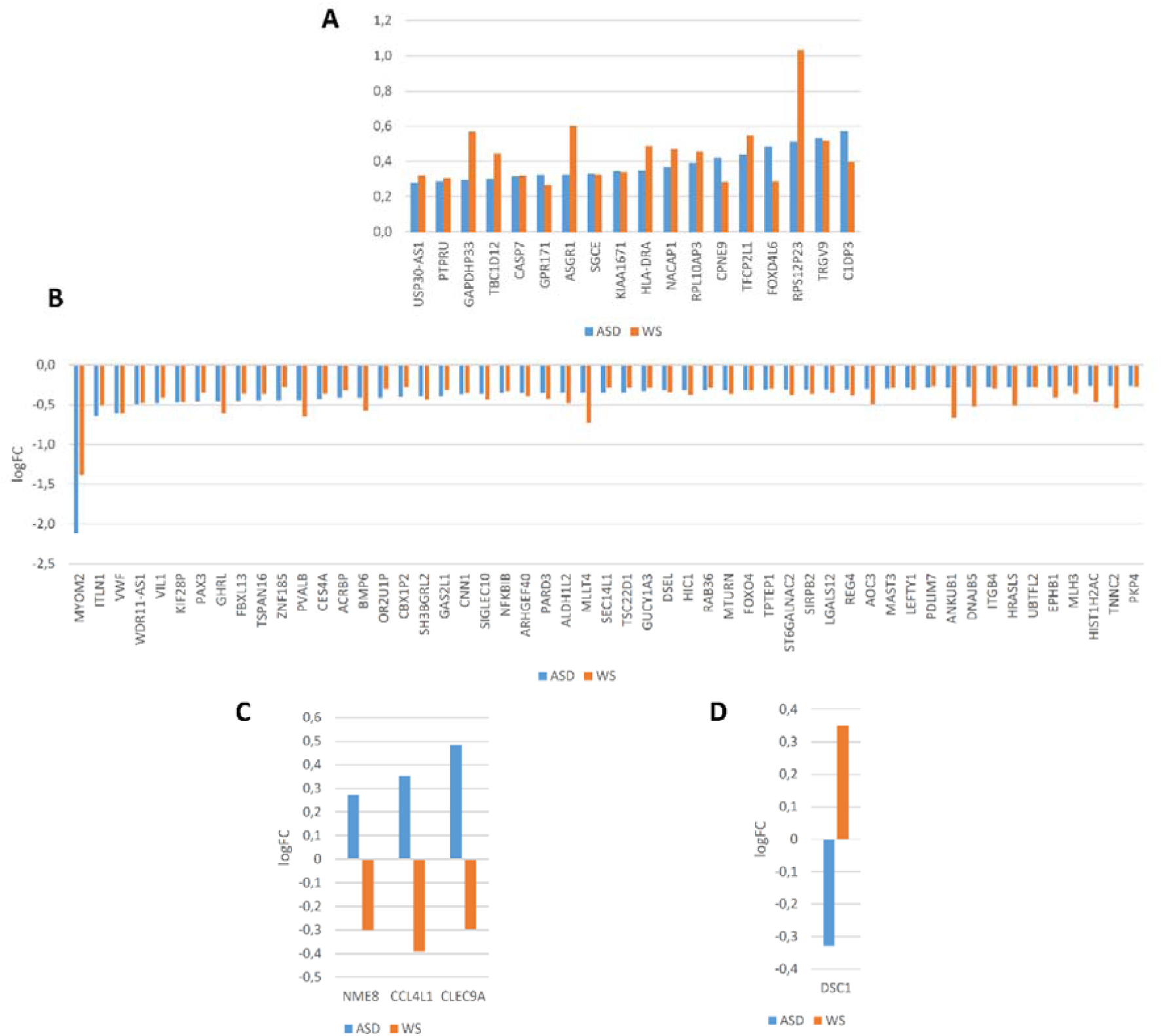
Abnormal expression patterns in the blood of subjects with ASD or WS. A. Genes found upregulated in both conditions. B. Genes found downregulated in both conditions. C. Genes found upregulated in ASD, but downregulated in WS. D. Genes found downregulated in ASD, but upregulated in WS. Histograms show changes in expression level of genes compared to neurotypical controls as fold changes (FC) if |FC| >1.2 and with false discovery rate (FDR<0.1). Values for ASD are colored in blue, whereas values for WS are colored in orange. Gene names are ordered according to the FC values for the ASD group.

We then interrogated whether these DEGs contribute to the etiopathogenesis of the socio-cognitive deficits of ASD and WS. Whereas genes found similarly dysregulated in the blood of people with ASD and WS can be expected to account for aspects of their similarities in the social cognition phenotype, the genes that exhibit opposite expression patterns might explain aspects of their differences in the socio-cognitive domain. Differences between these two conditions could result as well from fold change differences in the expression of genes that exhibit the same abnormal trends in ASD and WS.

## Functional characterization of genes of interest

As shown in Table 1, most of the DEGs (50 out of 75) have been associated to ASD and/or WS, are candidates for comorbid conditions, and/or might be involved in physiological aspects of relevance, mostly at the brain level, for the etiopathogenesis of the socio-cognitive dysfunctions observed in ASD and/or WS. A more detailed characterization of these genes is provided in Supplemental file 2.

**Table 1.**
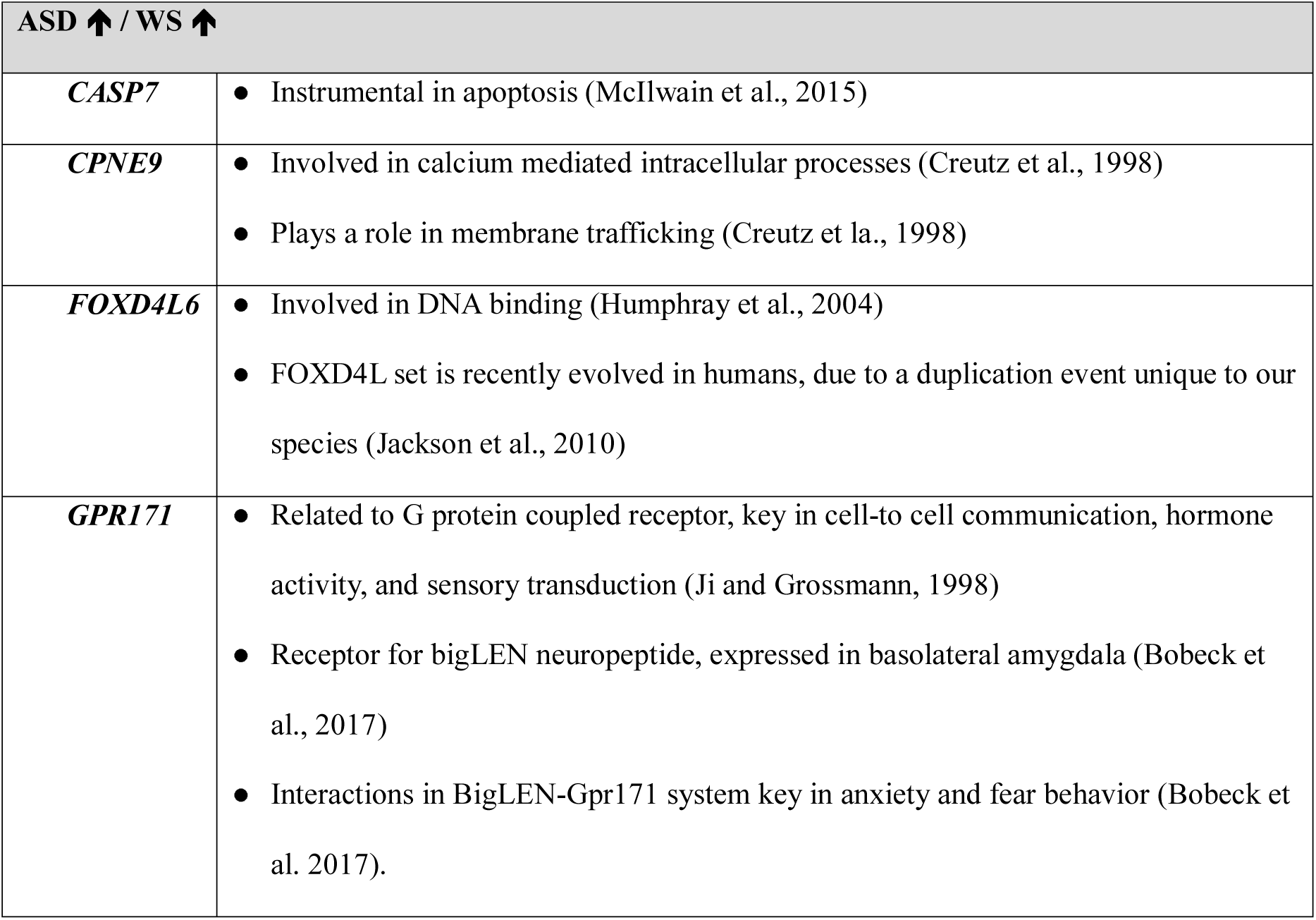
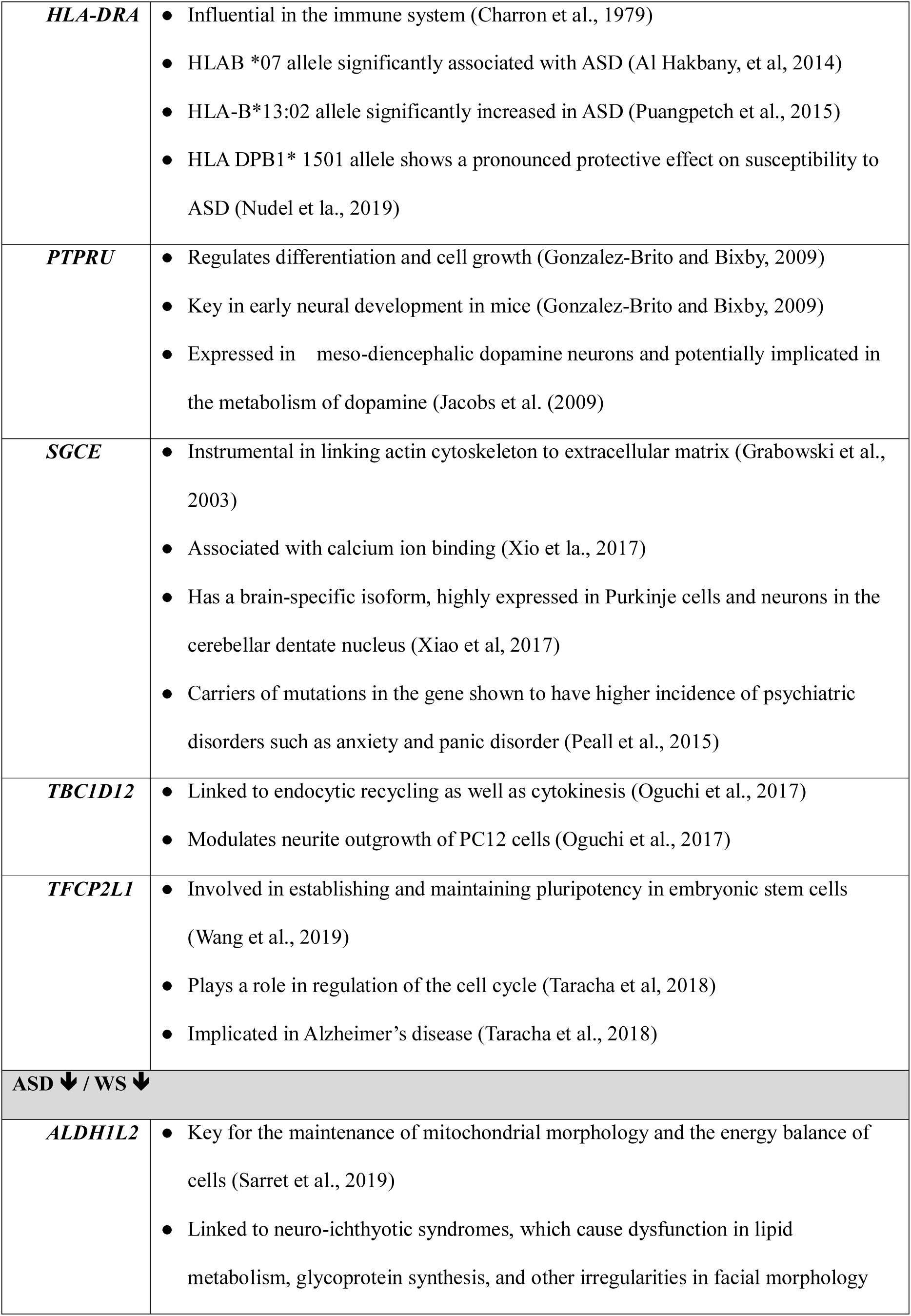
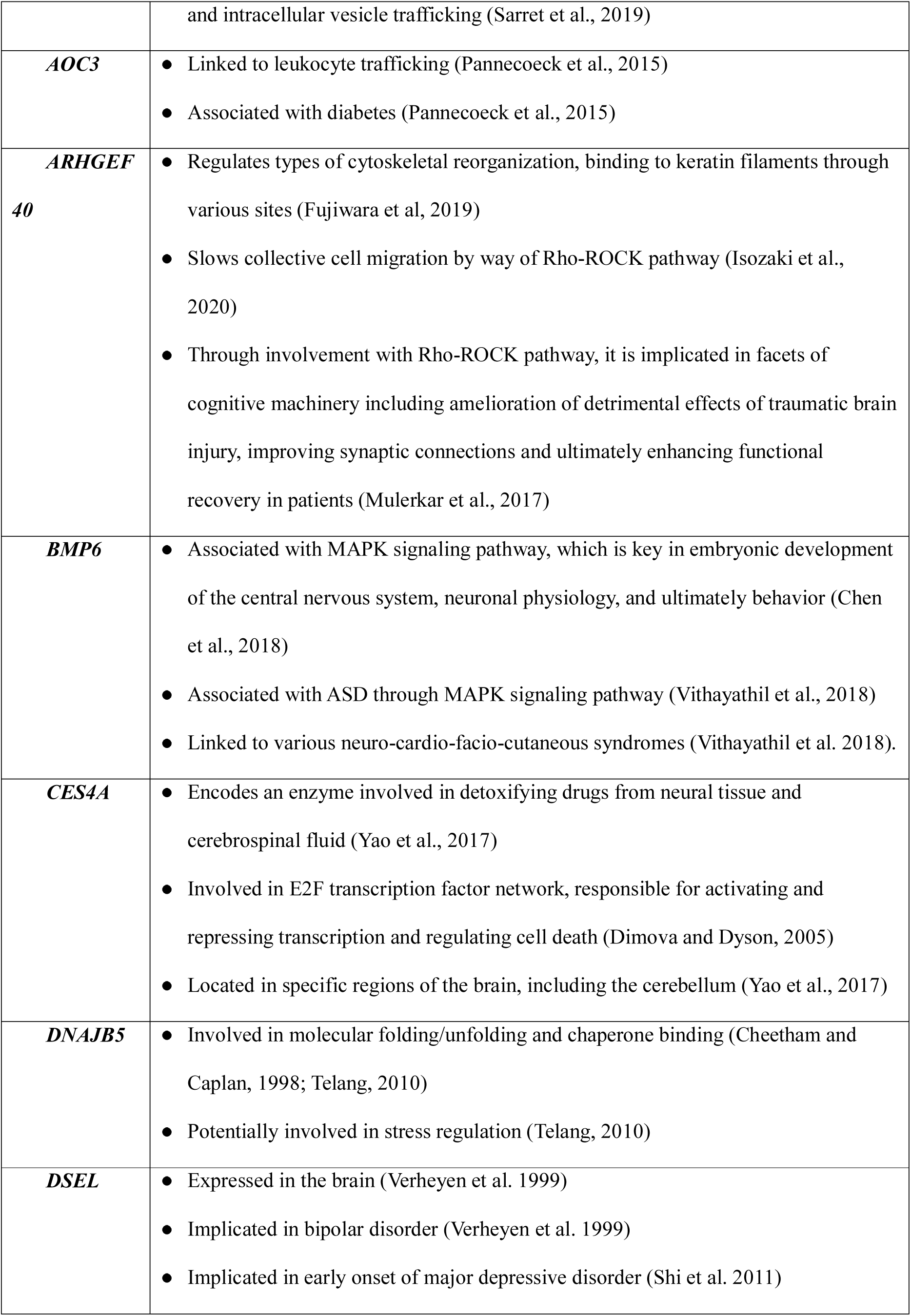
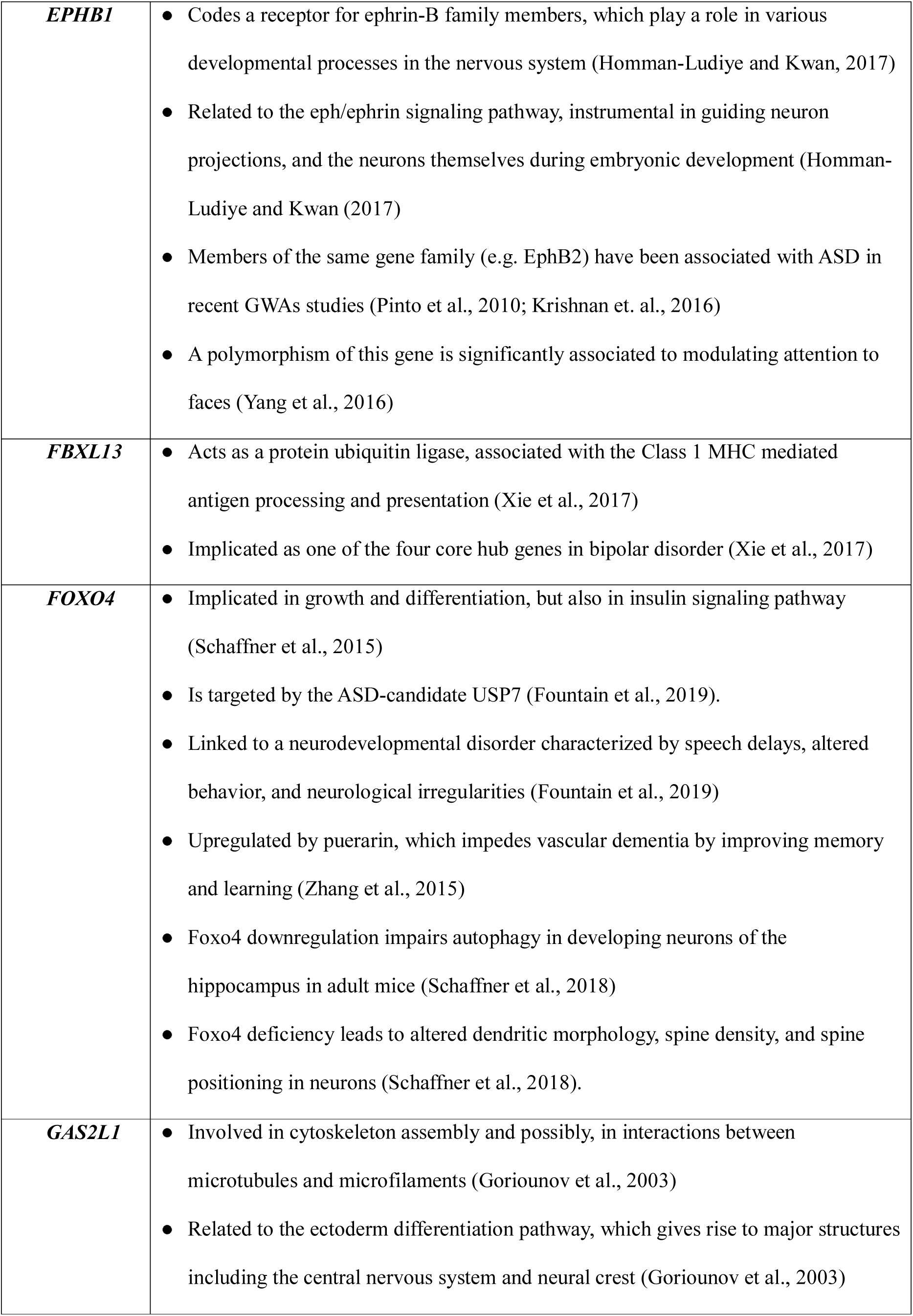
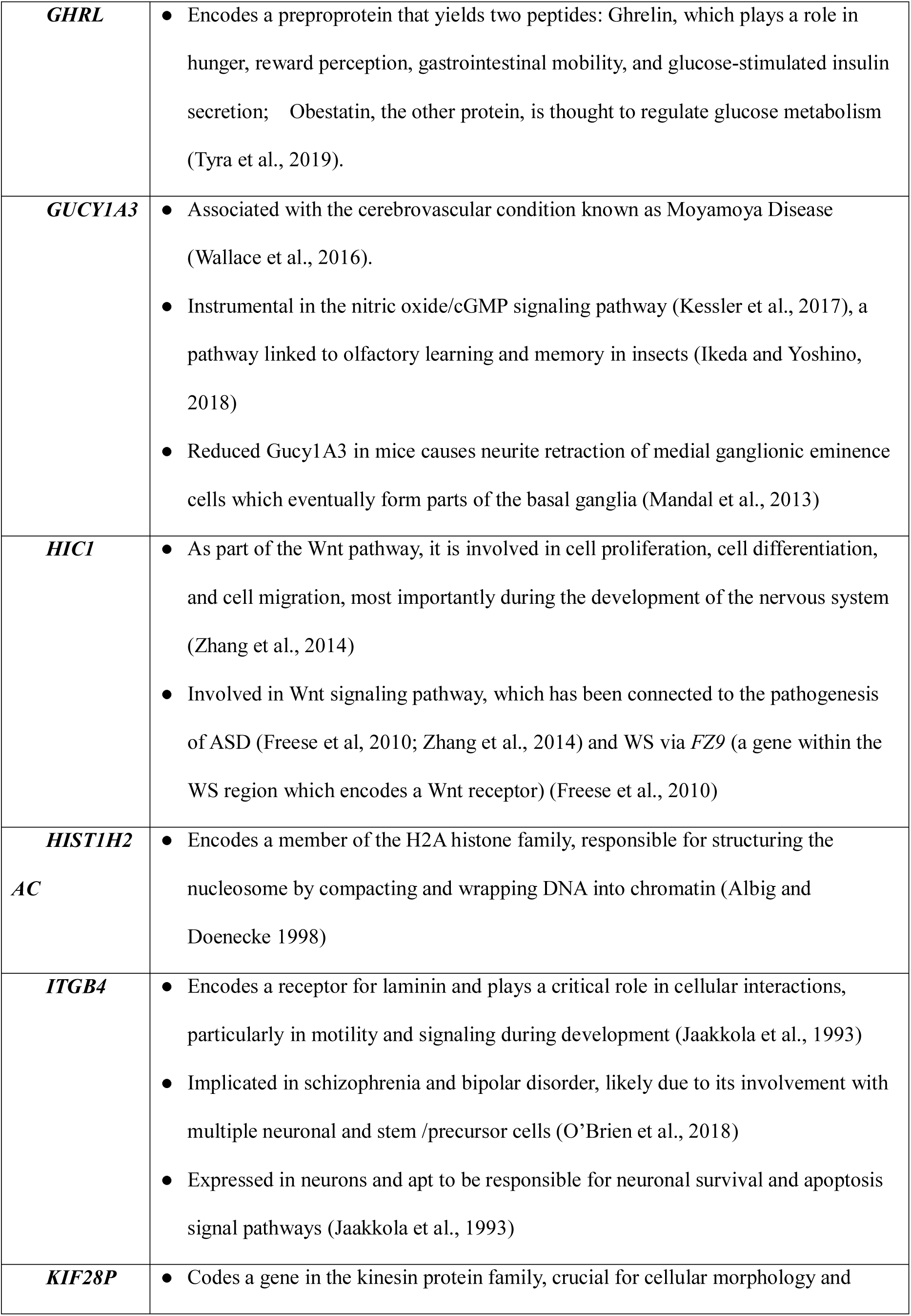
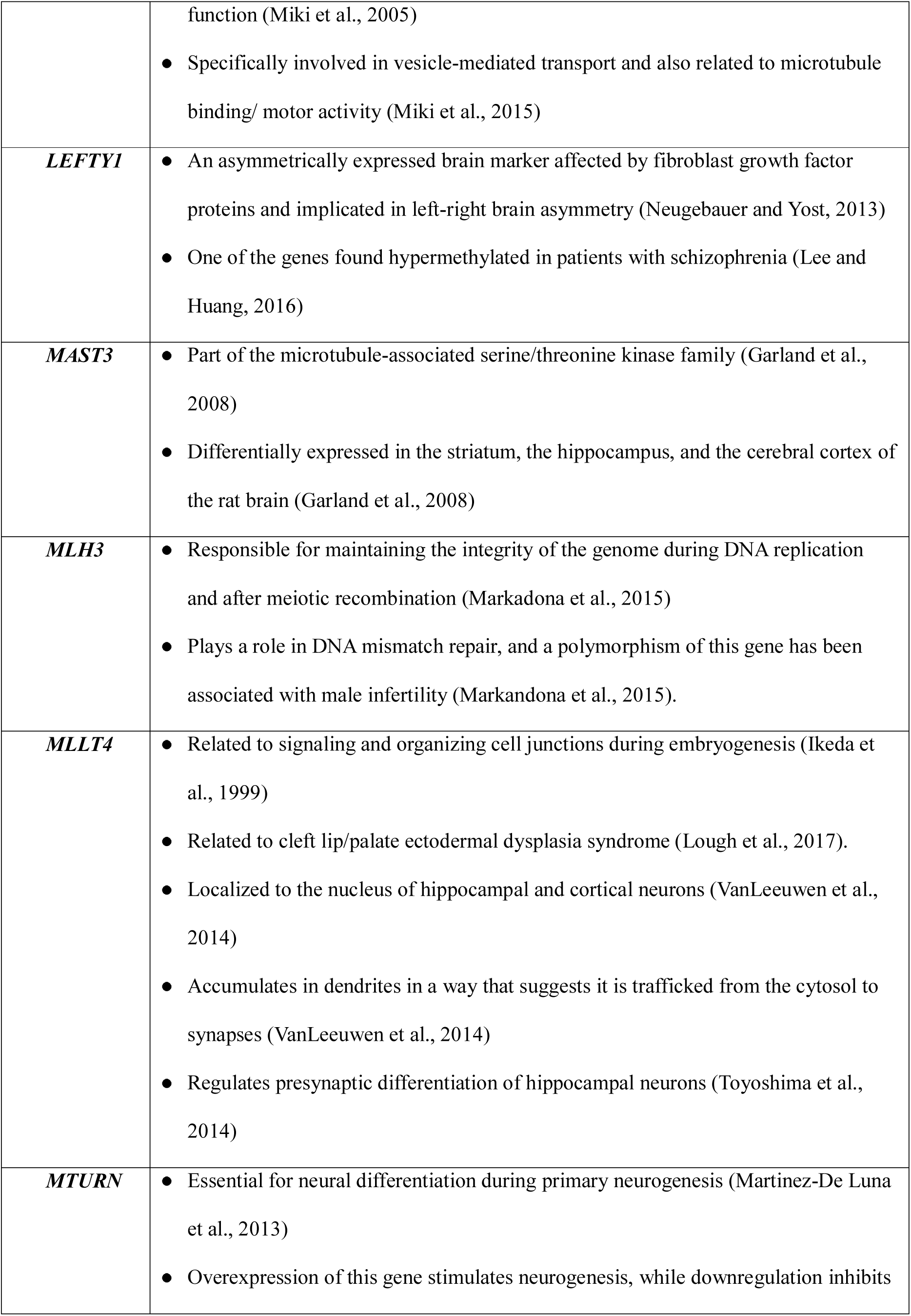
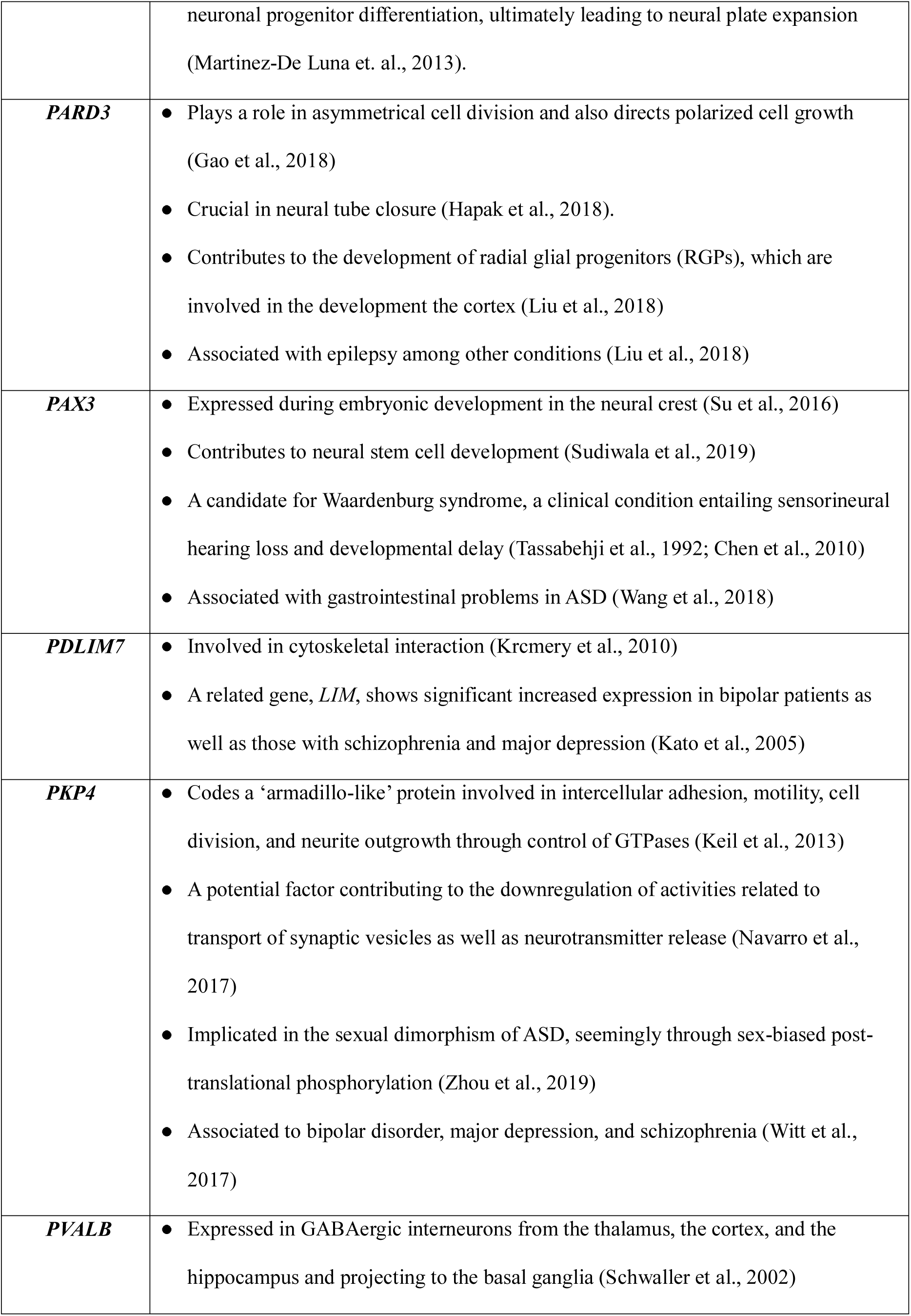
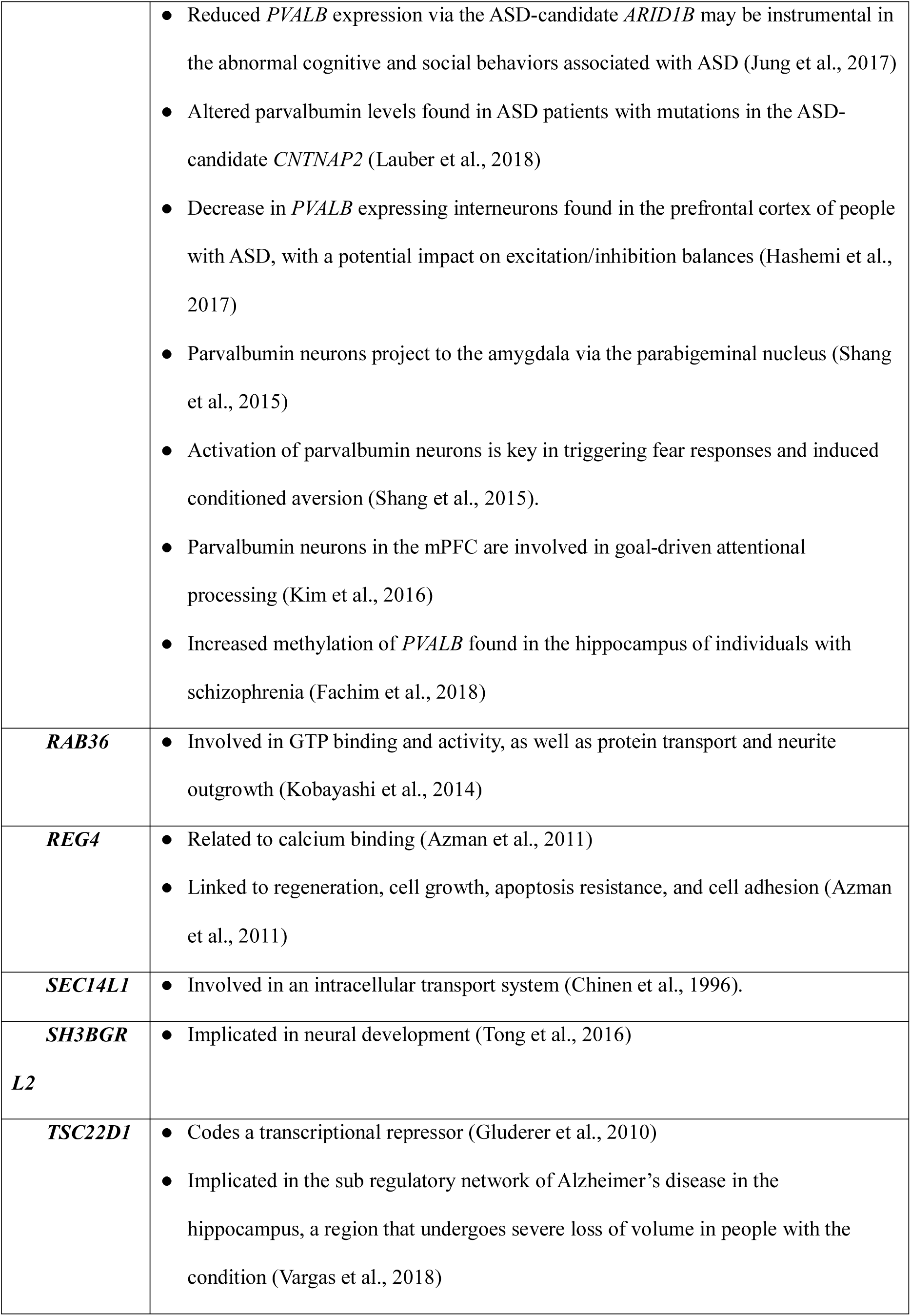
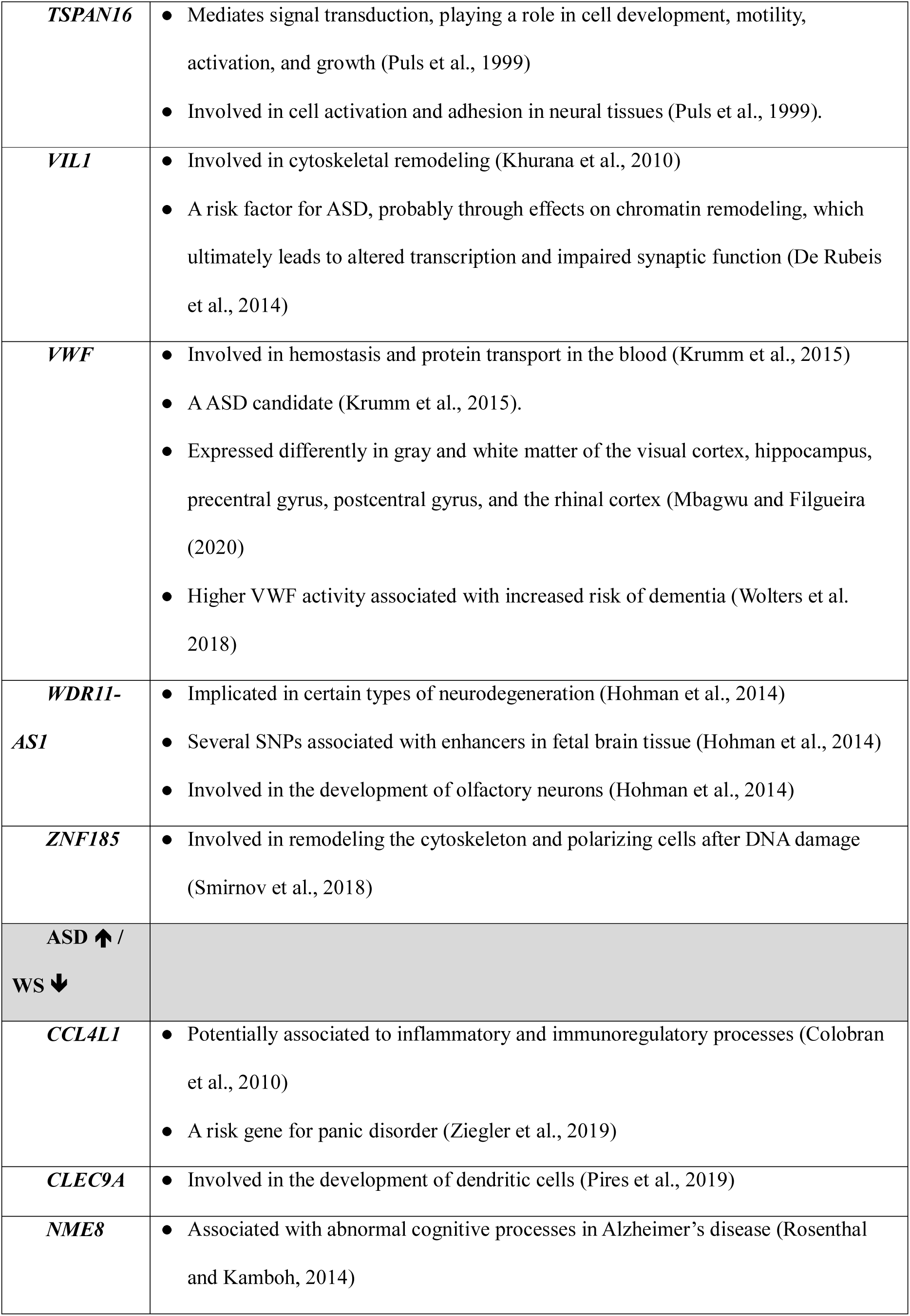
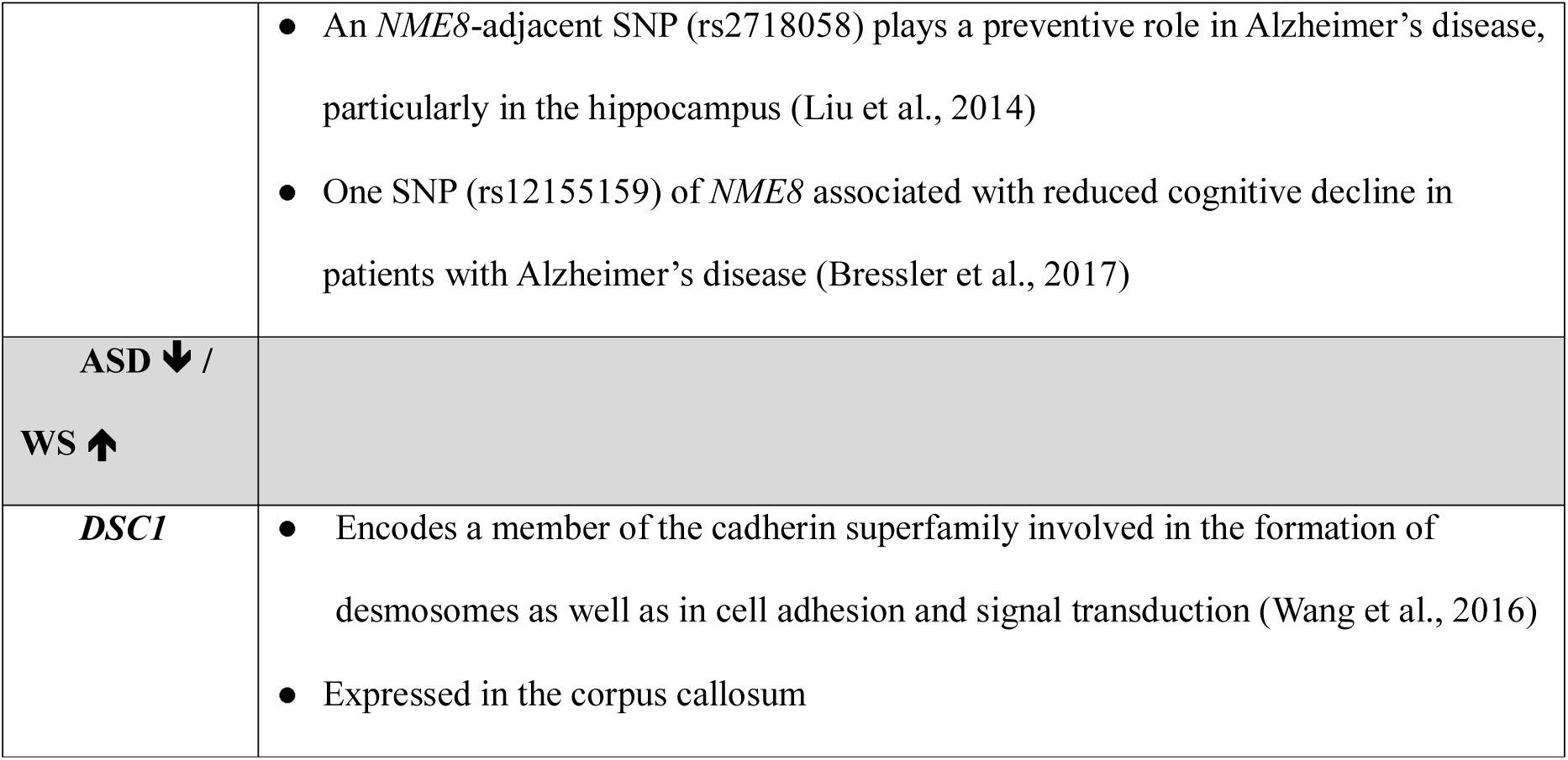
Summary list of the genes found dysregulated in subjects with ASD or WS that have a potential role in the etiopathogenesis of these conditions.

## Gene Ontology analyses

We also conducted GO analyses of the sets of DEGs to know more about possible biological functions that might be found similarly or differentially altered in ASD and WS, particularly, in connection with their distinctive socio-cognitive profiles. Because the number of genes showing opposite expression patterns was too small, GO analyses were performed only for genes found either upregulated or downregulated in the blood of subjects with ASD or WS compared to controls. We found that these two set of genes are significantly related to processes, functions, cellular components, and pathological phenotypes of interest for the ASD and the WS etiopathogenesis, with some interesting differences between both sets also existing (see Figure 3 for a graphical summary).

**Figure 3.**
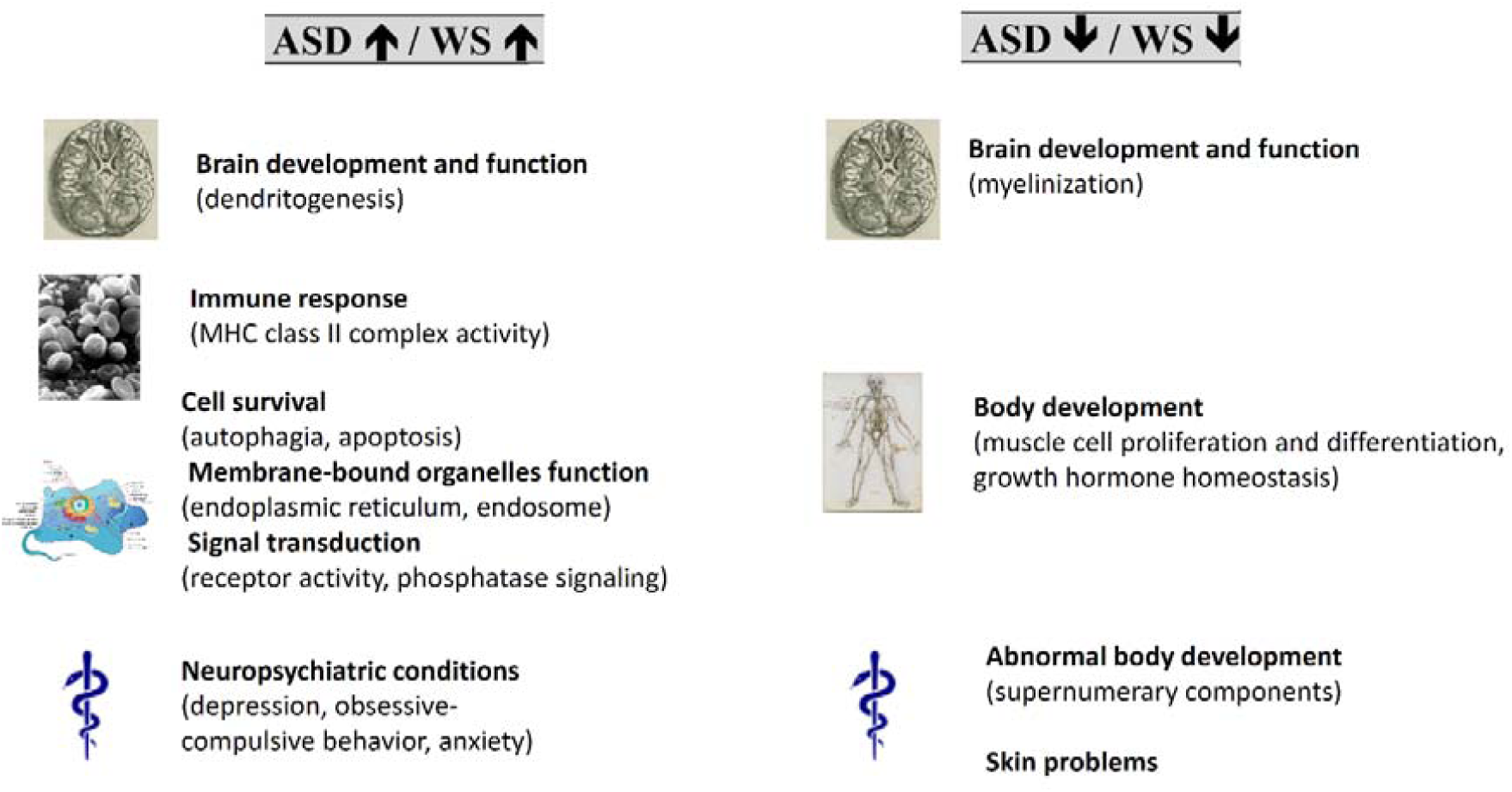
Graphical summary of GO analyses of genes found upregulated in both ASD and WS (left) and downregulated in both conditions (right). Illustrations are from Wikipedia (www.wikipedia.org), and all of them are in the public domain.

Accordingly, genes that are found upregulated (Figure 4) in both conditions compared to controls are significantly enriched in molecular, cellular, and biological processes important for neuron function, particularly dendrite extension (GO:1903861; GO:1903859). Multiple studies have pointed to alterations in dendrite growth, number, and morphology as a key aspect of the pathophysiology of ASD, with most alterations involving a generalized reduction in dendrite size and number, as well as an increase in spine densities with immature morphology (see Gilbert and Man, 2017, Martínez-Cerdeño, 2017, Joensuu et al., 2018 for selected reviews). Likewise, the neurons of animal models of selected genes within the WS region also exhibit an anomalous dendrite morphology. Accordingly, abnormal spine morphology, as well as abnormal synaptic function resulting in enhanced long-term potentiation (LTP) and altered fear responses and spatial learning, are observed in *Limk1*-knockout mice, supporting a role of this gene in the regulation of cofilin and actin cytoskeleton (Meng et al., 2002). Similarly, *FZD9* regulates dendritic spine formation in hippocampal neurons via its effect on Wnt5a signaling (Ramírez et al., 2016). Finally, neurite length is greater in mice lacking one copy of *Gtf2i*, another of the genes located within the chromosomal fragment deleted in WS (Deurloo et al., 2019). Additionally, genes upregulated in both conditions ASD and WS are preferentially associated to aspects of the immune response, particularly, to MHC class II complex function (GO:0032395; GO:0023026; GO:0042613). MHC genes have been found to be dysregulated in skin fibroblasts from people with WS (Henrichsen et al., 2011). Regarding ASD, because of the important role of the MHC in brain development and plasticity, changes in MHC expression resulting from mutations and/or immune dysregulation have been hypothesized to contribute to the altered brain connectivity and function typically found in subjects with this condition (see Needleman and McAllister, 2012 for review). Population-based epidemiological studies have found an association between ASD and MHC complex haplotypes (reviewed by Gesundheit et al., 2013). Additionally, genes upregulated in both ASD and WS are significantly associated with processes related to cell survival via autophagia, as in autophagosome assembly (GO:2000785), and to cell death via apoptosis, mostly in connection to cysteine-type endopeptidase activity (GO:0008635, GO:0097200, GO:0097153). In neurons, autophagy is involved in axon guidance, dendritic spine development and pruning, and synaptic plasticity (Hwang et al. 2019); altered autophagy has been associated as well to neurodegeneration (see Lee, 2012 for review) and to ASD (Hwang et al., 2019). Apoptosis is also crucially involved in brain development and wiring, and pathological activation of apoptotic death pathways resulting in neural cell death has been equally associated to ASD (Wei et al. 2014), particularly, endoplasmic reticulum stress resulting in apoptosis (Dong et al., 2018). Interestingly, genes upregulated in both ASD and WS are preferentially associated to the endoplasmic reticulum membrane (GO:0071556), as well as the endocytic vesicle membrane (GO:0030669; GO:0045334), the endosome membrane (GO:0031902) and the endoplasmic reticulum to Golgi vesicle membrane (GO:0012507). In a similar vein, increased apoptosis has been observed in animal models of WS, particularly, after knocking out several of the genes within the WS fragment, specifically, *WSTF* (Barnett et al., 2012) and *FZD9* (Zhao et al., 2005). Genes upregulated in both ASD and WS are also enriched in proteins involved in signal transduction activities, particularly receptor activity, mostly linked to G-protein (GO:0001608; GO:0045028) and protein tyrosine phosphatase signaling (GO:0007185; GO:0005001; GO:0019198).

**Figure 4.**
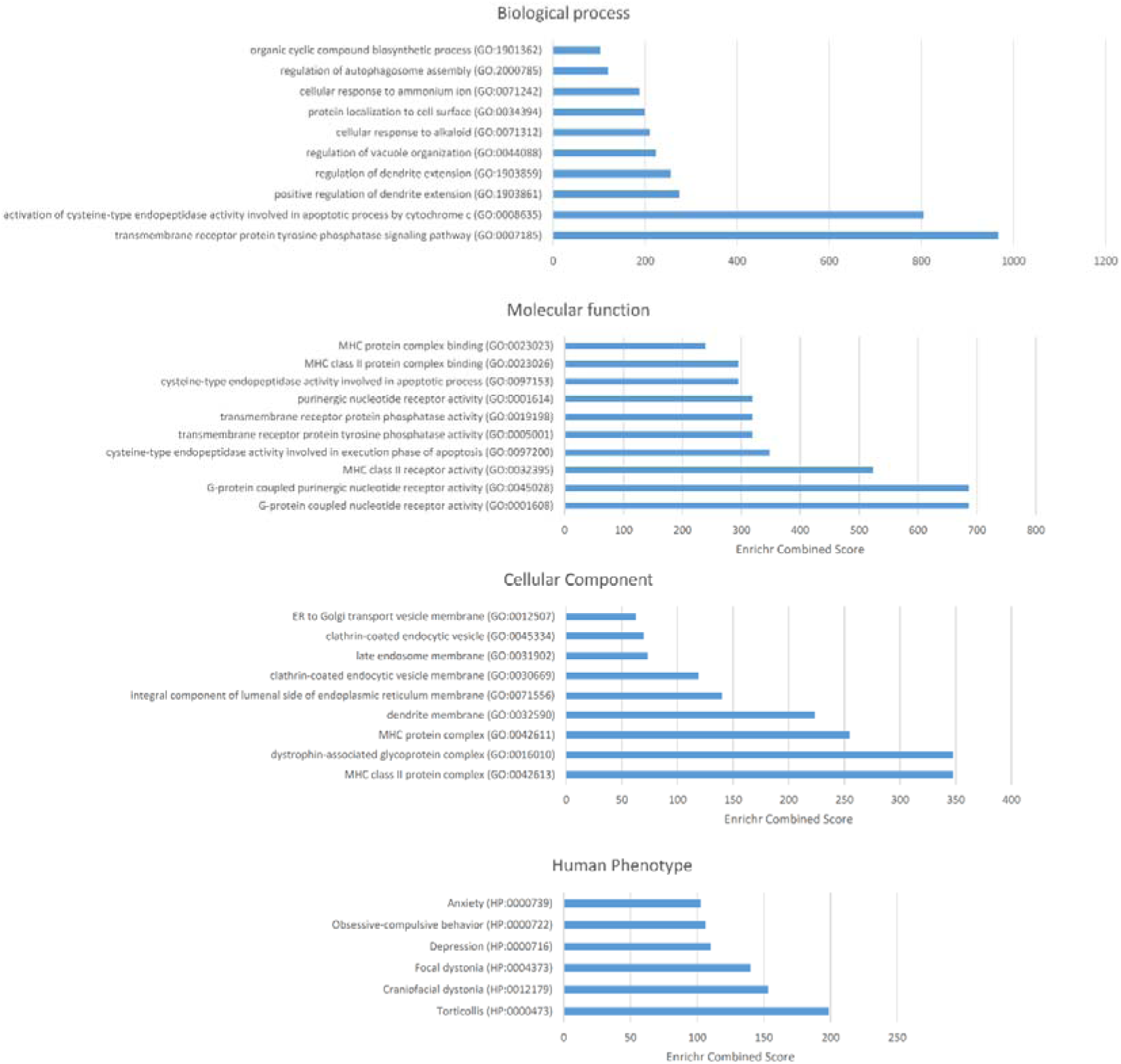
Functional enrichment analysis according to Enrichr of the set of genes that are significantly upregulated in the blood of subjects with WS or ASD compared to neurotypical controls. The figure shows the enrichment in biological processes, molecular function, cellular components, and human pathological phenotypes (from top to bottom). Only the top-10 functions have been included and only if their p<0.05. The p-value was computed using Fisher’s exact test. Enriched categories are ordered according to their Enrichr Combined Scores. This is a combination of the p-value and the z-score calculated by multiplying the two scores (Combined Score = ln(p-value) * z-score). The z-score is computed using a modification to Fisher’s exact test and assess the deviation from the expected rank. The Combined Score provides a compromise between both methods and it is claimed to report the best rankings when compared with the other scoring schemes. See http://amp.pharm.mssm.edu/Enrichr/help#background&q=5 for details.

Finally, it is worth highlighting that these genes are significantly related to neuropsychiatric conditions like depression (HP:0000716), obsessive-compulsive behavior (HP:0000722), and anxiety (HP:0000739), all of which are comorbid or share core symptoms with ASD (Matson and Nebel-Schwalm, 2007; Vannucchi et al., 2014; Rossen et al., 2018). As discussed in the previous section, anxiety is also prevalent within people with WS, although depression is also occasionally diagnosed in affected people (Stinton et al., 2010; 2012; Royston et al., 2017).

As shown in Figure 2, most genes upregulated in ASD and WS compared to controls exhibit similar fold changes in both conditions. However, some of them are more strongly upregulated in WS than in ASD, particularly, *GAPDHP33*, *ASGR1* and *RPS12P23*. Interestingly, a GWA analysis has associated *ASGR1* to animal personality and coping styles, particularly to latency, duration and frequency of struggling attempts by piglets during backtests, with one SNPs showing differential expression in the hypothalamus (Ponsuksili et al., 2015). Dosage perturbation of this gene has also probed to impact negatively on neurodevelopment in zebrafish, ultimately resulting in microcephaly, thus seemingly contributing to the cognitive symptoms of the 17p13.1 microdeletion syndrome, which include intellectual disability and poor to absent speech, as well as occasional autistic features (Carvalho et al., 2014).

Regarding the genes that are downregulated in both ASD and WS compared to controls (Figure 5), we found that they are significantly related to muscle cell proliferation and differentiation (GO:0014842; GO:0051151), but also to growth hormone homeostasis, particularly, growth hormone secretion (GO:0060124; GO:0030252) and insulin-like growth factor binding (GO:0031994; GO:0005520). These genes are also related to aspects of brain development, particularly, myelinization (GO:0031643). Children with ASD are known to exhibit an early generalized overgrowth (Van Daalen et al. 2007, Fukumoto et al. 2008, Chawarska et al. 2011), with postnatal overgrowth correlating with greater severity of social deficits and poorer verbal skills (Chawarska et al. 2011, Campbell et al. 2014). Significantly higher levels of several growth-related hormones have been found in children with ASD (Mills et al., 2007). Dysregulation of the overall systemic growth seems to account for the brain overgrowth also frequently observed in people with ASD, which correlates with lower functioning abilities (Sacco et al. 2015). Higher head circumference and increased brain size values are usually observed only during early childhood (Fukumoto et al. 2008, Courchesne et al. 2011, although see Raznahan et al. 2013). By contrast, children with WS typically show growth retardation, at least during their first years of life (Pankau et al., 1992; Morris et al., 1998). Still, growth hormone deficiency is rarely diagnosed (e.g. Levy-Shraga et al., 2018). Likewise, subjects with WS have smaller brain volumes compared to controls (Jernigan and Bellugi, 1990; Schmitt et al., 2001a; Reiss et al., 2004, Thompson et al., 2005 Meyer-Lindenberg et al., 2005; Jackowski et al., 2009), mostly due to a reduction of white matter (Thompson et al., 2005). Interestingly too, the insulin-like growth factor I, which is primarily involved in the regulation of the effects of growth hormone, also contributes to neural development, myelinisation, and protection (see Puche and Castilla-Cortázar, 2012 for review). Finally, the genes found downregulated in both ASD and WS are significantly associated to clinical phenotypes mostly involving an abnormal body development, like rhabdomyosarcoma (HP:0002859), supernumerary ribs (HP:0005815) or supernumerary bones of the axial skeleton (HP:0009144); or different skin problems, like white forelock (HP:0002211), aplasia cutis congenita (HP:0001057), patchy hypopigmentation of hair (HP:0011365), which are of less interest for the socio-cognitive profile of people with ASD and WS.

**Figure 5.**
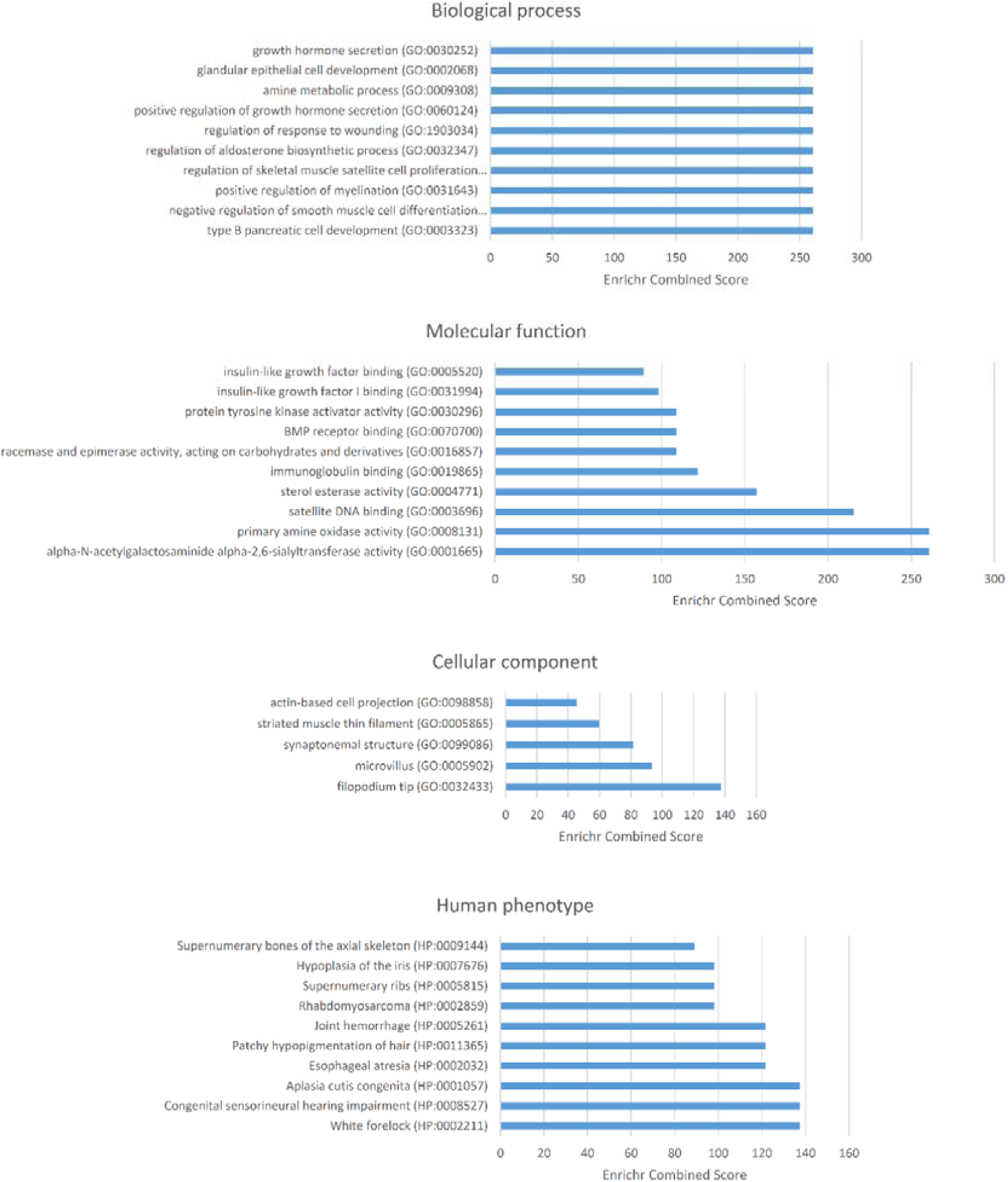
Functional enrichment analysis according to Enrichr of the set of genes that are significantly downregulated in the blood of subjects with WS or ASD compared to neurotypical controls. The figure shows the enrichment in biological processes, molecular function, cellular components, and human pathological phenotypes (from top to bottom). Only the top-10 functions have been included and only if their p<0.05. The p-value was computed using Fisher’s exact test. Enriched categories are ordered according to their Enrichr Combined Scores. This is a combination of the p-value and the z-score calculated by multiplying the two scores (Combined Score = ln(p-value) * z-score). The z-score is computed using a modification to Fisher’s exact test and assess the deviation from the expected rank. The Combined Score provides a compromise between both methods and it is claimed to report the best rankings when compared with the other scoring schemes. See http://amp.pharm.mssm.edu/Enrichr/help#background&q=5 for details.

As shown in Figure 2, most genes downregulated in both ASD and WS compared to controls exhibit similar fold changes in these two conditions. However, some of them are more strongly downregulated in WS than in ASD, particularly, *MLT4*, *ANKUB1*, *DNAJB5*, *HRASLS* and *TNNC2*. By contrast, *MYOM2* is more strongly downregulated in ASD than in WS. Interestingly, *DNAJB5* has been associated to response to social eavesdropping in zebrafish, particularly, to changes in the alertness status (Lopes et al., 2015).

## Expression pattern analyses

Finally, we report on the expression profiles of the genes that have been found to be dysregulated in the blood of subjects with WS or ASD, with a focus on the brain, considering our interest on behavioral and cognitive features. A heat map of the expression levels of these genes in the samples of the Human tissue compendium (Novartis) (Su et al., 2004), as generated by Gene Set Enrichment Analysis (GSEA) software (http://software.broadinstitute.org/gsea/index.jsp), is shown in Figure 6. According to the Human Gene Atlas (Su et al., 2004), genes found differentially upregulated in ASD and WS are predicted to be preferentially expressed in the thalamus (p = 0.06879; Enrichr combined score = 37.65). By contrast, genes found differentially downregulated in both conditions are predicted to be significantly expressed in the cerebellum (p = 0.03330; Enrichr combined score = 23.78). We also checked the expression profile of these two groups of genes during development via the Human Brain Transcriptome Database (http://hbatlas.org). We found that all the genes that are differentially upregulated in subjects with ASD or WS are expressed in the brain thorough development, but whereas some of them are expressed at similar levels across regions, others exhibit different expression levels in different brain areas, tending to be more expressed outside the neocortex, particularly, in the thalamus, but also in the striatum and the amygdala (Supplemental file 3). Also all the genes that are differentially downregulated in subjects with ASD or WS are expressed in the brain during development. However, among these genes, those exhibiting different levels of expression in different brain areas tend to be more expressed in the cerebellum, but also in the striatum (supplemental file 3). Overall, these findings suggest that abnormal gene upregulation can be expected to impact mostly on the thalamus, whereas downregulated genes tend to be more involved in cerebellar development and function. Both groups are expected to affect striatal regions.

**Figure 6.**
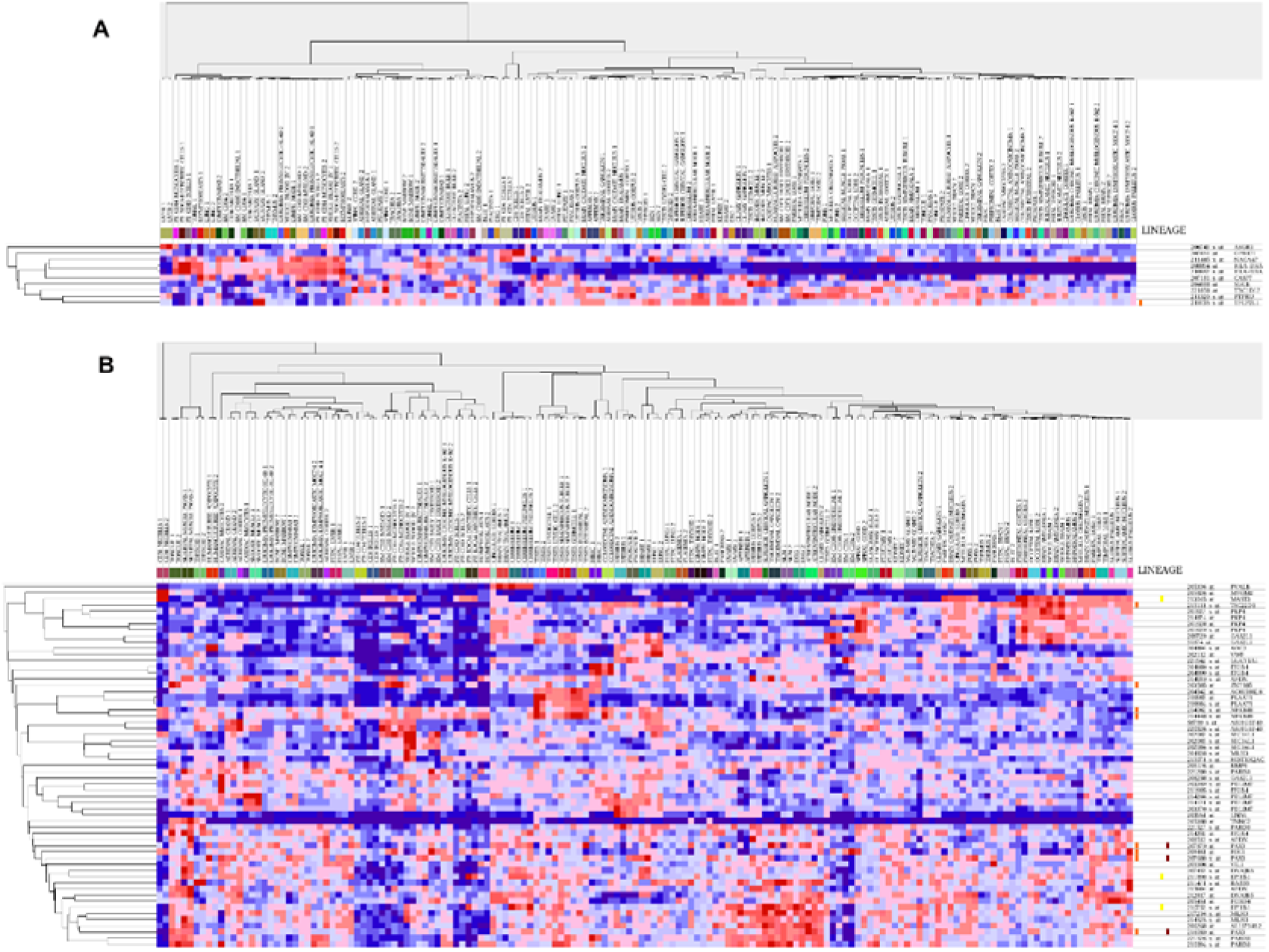
Heat maps of the expression levels of the set of genes that are significantly dysregulated in the blood of subjects with ASD or WS compared to controls. A. Heat map of the genes found upregulated in both conditions. B. Heat map of the genes found downregulated in both conditions. The heat maps were generated by the Gene Set Enrichment Analysis (GSEA) software using the samples of the Human tissue compendium (Novartis) (Su et al., 2004). GSEA is a computational method that determines whether an a priori defined set of genes shows statistically significant, concordant differences between two biological states. The heat maps include dendrograms clustering gene expression by gene and samples. Genes are identified by probe identifier, gene symbol, description, and gene family. See http://software.broadinstitute.org/gsea/index.jsp for details.

In subjects with WS, the thalamus shows structural and functional differences compared to neurotypical controls, including smaller volumes, reduced gray matter, and enhanced activity (Reiss et al., 2000; Schmitt et al., 2001; Tomaiuolo et al., 2002; Meyer-Lindenberg et al., 2004; Mobbs et al., 2007; Bódizs et al., 2012). Although thalamic gray matter reduction has been associated to their visuospatial impairment (Chiang et al., 2007; Campbell et al., 2009), abnormal thalamic development and function can be expected to contribute as well to other cognitive deficits, particularly language problems, because of the role of the thalamus as a sort of relay center, connecting many of the brain areas involved in language processing (Wahl et al., 2008; Murdoch, 2010; David et al., 2011). In people with ASD, in spite of previous inconsistent findings, recent research points to structural abnormalities in the thalamus too (Schuetze eal., 2016), as well as a to a disruption of local thalamic connectivity and a dysregulation of thalamo-cortical networks (Woodward et al., 2017; Tomasi and Volkow, 2019). Likewise, regarding the striatal regions, people with WS have reduced basal ganglia volumes (Jernigan et al., 1993, Bellugi et al., 1999, Reiss et al, 2000; Campbell et al., 2009). Moreover, children with Asperger syndrome show reduced volumes of the caudate, whereas high-functioning children with ASD exhibit smaller gray matter volumes in the frontostriatal regions (McAlonan et al., 2008). By contrast, adults with ASD exhibit a relative increase of both caudal putamen and pallidum, with restricted, repetitive behaviors positively correlating with surface area in bilateral globus pallidus (Schuetze et al., 2016). The basal ganglia control different cognitive and emotional functions, including language (Booth et al., 2007; Kotz et al., 2009; Viñas-Guasch and Wu, 2017). Overall, these findings point to changes in intrathalamic and transthalamic routes as an important cause of the perceptual, motoric, interoceptive, emotional, and cognitive impairments found in ASD. Specifically, as we discussed in depth in the previous section, socio-cognitive similarities between ASD and WS may stem from the disruption of striatal and thalamic connections, particularly in the domain of face processing and theory of mind (involving the thalamus), reward behavior (involving the striatum), and attention switching (involving the striatum and the thalamus). Finally, in people with WS, the cerebellum exhibits volume alterations that can be associated with their distinctive cognitive, affective and motor features (Osório et al., 2014). This is particularly true of language, given the role of the cerebellum in language processing and its impairment in language-related pathologies (Vias and Dick, 2017; Mariën and Borgatti 2018). In a similar vein, genetic, molecular, behavioral, and neuroimaging findings support the view that in people with ASD the cerebellum develops differently at multiple levels of neural structure and function, this circumstance contributing to facets of their behavioral, cognitive and affective distinctive profile (Becker and Stoodley, 2013; Hampson and Blatt, 2015).

## 5. Discussion

In this paper we have adopted a comparative-molecular approach in order to gain some insights about the causes of the deficits exhibited by people with ASD or WS in the domain of social cognition and social behavior. As discussed in sections 2 and 3, the ASD phenotype (loosely) mirrors the WS phenotype, although some overlap exists between both conditions. The same can be said of some of the genes contributing to these conditions, since genes within the WS region are also candidates for ASD (Sanders et al., 2011), and with subjects with WS having some risk of suffering from autistic behaviors (Tordjman et al., 2012, Klein-Tasman et al., 2018). However, to date most studies have focused on individual genes. For instance, one key gene within the WS region, namely *GTF2I*, which is regularly associated to the social phenotype of WS, has been found to be a risk factor for ASD (Malenfant et al., 2012). Interestingly, whereas hypersocial behavior has been observed in mice carrying a deletion of *Gtf2i*, no evidence of hyposocial behavior has been observed in mice with the *Gtf2i* duplication (Martin et al., 2017). This contradicts simplistic models of these diseases according to which socio-cognitive differences between ASD and WS might result from dosage changes in selected genes. At the very least, whole-genome analyses should be conducted in order to provide a more comprehensive view of the genetic mechanistics of the similarities and particularly, the differences between these two conditions.

In our study we have found that ASD and WS share a common pattern of gene dysregulation in the blood of affected people. The fact that most of the abnormally-expressed genes are found either upregulated or dysregulated in both conditions might account for their similarities in the socio-cognitive domain, particularly if one considers that most of these genes are involved in aspects of brain (abnormal) development and/or (dys)function. Accordingly, we have found that a significant number of the DEGs contribute to brain development and function (particularly, dendritogenesis) and are expressed in brain areas (particularly, the cerebellum, the thalamus and the striatum) of relevance for the ASD and the WS etiopathogenesis. Nonetheless, some remarkable phenotypical and neurobiological differences also exist between ASD and WS. We hypothesize that they might result in part from the opposite expression pattern exhibited by a small groups of genes, in part from the circumstance that some of the genes showing similar expression trends in ASD and WS still exhibit quantitative differences between conditions, with most of them being more dysregulated in WS than in ASD. Overall, the genes we highlight in the paper emerge as promising candidates for explaining the similarities and differences between ASD and WS in their social cognition and social behavior.

That said, caution is in order. First, our study has limitations. Besides the small size of our samples of patients, the abnormal patterns of gene expression uncovered by the microarray analyses need to be validated via other techniques e.g. RT-PCR. Second, a direct translation of gene dosage changes to differences in the severity (or even the nature) of abnormal phenotypical traits or disease symptoms should be avoided unless empirical evidence does exist. Actually, in many cases this translation is not observed, even for particular genes, as noted above. When several genes are involved, like in conditions resulting from copy-number variations (CNVs), mirror phenotypes are not usually found either. For instance, Smith-Magenis syndrome and Potocki-Lupski syndrome are reciprocal contiguous gene syndromes resulting from the microdeletion and the microduplication, respectively, of the same chromosomal region; although patients exibit some mirror traits, others are shared between conditions, with only a minor number of genes within the critical genomic region being dosage-sensitive (Neira-Fresneda and Potocki, 2015). More generally, the mechanisms by which gene-dosage changes result in disease are frequently opaque (see Rice and McLysaght, 2017 for discussion). Third, from translational medicine point of view, the gene expression changes we highlight in the paper should be regarded more a biomarker of ASD and WS than a mechanistic account of specific deficits of ASD and/or WS via the dysfunction of specific genes (incidentally, this is why in our analysis we have focused on GO analyses, instead of the functions performed by each of the DEGs). Alterations in the expression of individual genes can be potentially caused by unknown coincident events, can be of no developmental/biological relevance, and/or can be an adaptive response to the changes affecting to other genes actually contributing to the disease. But even in this case, caution is in order too. One reason is that we have attested differences in gene expression levels in adult subjects compared to neurotypical controls. Nonetheless, ASD and WS are developmental conditions, this entailing that changes in gene dosage could differ from one developmental stage to another, and/or impact differentially at different stages of development. Accordingly, even if these abnormal gene expression profiles could be used as reliable biomarkers of these two conditions, they should be validated in children, for whom an early diagnosis (particularly, of ASD) is intended.

In sum, more research is needed before promoting the genes we highlight in the paper to key casual factors in the emergence of the deficits observed in ASD and WS. To achieve this, besides the validation of our findings, as noted above, other approaches should be adopted, particularly, conducting additional in vitro and in vivo research (including the development of animal models of selected candidates) aimed to gain direct insights of the mechanistic of these genes, particularly, their contribution to brain development and function in areas involved in social cognition.

## Acknowledgements

The authors thank Ryo Kimura, from the Department of Anatomy and Developmental Biology, at the Graduate School of Medicine of Kyoto University, for his help with data analyses. The author declares no conflict of interests.

## Supplemental data

Supplemental file 1. List of genes considered in this analysis

Supplemental file 2. Functional characterization of genes found dysregulated in the blood of subjects with ASD or WS and with a potential impact on the etiopathogenesis of these conditions.

Supplemental file 3. Developmental expression profiles of the genes dysregulated in the blood of people with ASD or WS compared to controls according to the Human Brain Transcriptome Database.

## References

1. Adolphs R., Spezio M. (2006). Role of the amygdala in processing visual social stimuli. Prog. Brain Res. 156: 363–378.

2. Albig W, Doenecke D (1998). The human histone gene cluster at the D6S105 locus. Hum. Genet. 101 (3): 284–94.

3. Al-Hakbany M, Awadallah S, Al-Ayadhi L. (2014). The Relationship of HLA Class I and II Alleles and Haplotypes with Autism: A Case Control Study. Autism Res Treat. 2014: 242048

4. Allison T., Puce A., McCarthy G. (2000). Social perception from visual cues: role of the STS region. Trends Cogn. Sci. 4: 267–278.

5. Amodio D. M., Frith C. D. (2006). Meeting of minds: the medial frontal cortex and social cognition. Nat. Rev. Neurosci. 7: 268–277.

6. Amaral D. G., Schumann C. M., Nordahl C. W. (2008). Neuroanatomy of autism. Trends Neurosci. 31: 137–145.

7. Andari E., Duhamel J. R., Zalla T., Herbrecht E., Leboyer M., Sirigu A. (2010). Promoting social behavior with oxytocin in high-functioning autism spectrum disorders. Proc. Natl. Acad. Sci. U.S.A. 107: 4389–439410.

8. Asada K, Itakura S (2012). Social phenotypes of autism spectrum disorders and Williams syndrome: similarities and differences. Front Psychol. 3:247.

9. Assaf M, Jagannathan K, Calhoun VD, Miller L, Stevens MC, Sahl R, O’Boyle JG, Schultz RT, Pearlson GD (2010). Abnormal functional connectivity of default mode sub-networks in autism spectrum disorder patients. Neuroimage. 53(1):247–56.

10. Avery SN, Thornton-Wells TA, Anderson AW, Blackford JU. (2012). White matter integrity deficits in prefrontal-amygdala pathways in Williams syndrome. Neuroimage. 59: 887–94.

11. Azman J, Starcevic Klasan G, Ivanac D, Picard A, Jurisic-Erzen D, Nikolic M, Malnar D, Arbanas J, Jerkovic R (2011). Reg IV protein and mRNA expression in different rat organs. Acta Histochem.113(8):793-7.

12. Barak B, Feng G (2016). Neurobiology of social behavior abnormalities in autism and Williams syndrome. Nat Neurosci. 19(6):647–655.

13. Barnett C, Yazgan O, Kuo HC, Malakar S, Thomas T, Fitzgerald A, Harbour W, Henry JJ, Krebs JE. (2012). Williams Syndrome Transcription Factor is critical for neural crest cell function in Xenopus laevis. Mech Dev. 1299–12:324-38.

14. Baron-Cohen, S., Leslie, A. M., Frith, U. (1985). Does the autistic child have a ‘‘theory of mind’’? Cognition, 21(1), 37–46.

15. Baron-Cohen S, Wheelwright S (1999). ’Obsessions’ in children with autism or Asperger syndrome. Content analysis in terms of core domains of cognition. Br J Psychiatry. 175:484–90.

16. Becker EB, Stoodley CJ. (2013) Autism spectrum disorder and the cerebellum. Int Rev Neurobiol. 113:1–34.

17. Bellugi, U., Lichtenberger, L., Jones, W., Lai, Z., St.George, M. (2000). The neurocognitive profile Of Williams syndrome:a complex pattern of strengths and weaknesses. J. Cogn.Neurosci. 12(Suppl.),7– 29.

18. Bellugi U, Adolphs R, Cassady C, Chiles M (1999). Towards the neural basis for hypersociability in a genetic syndrome. Neuroreport. 10:1653–7.

19. Bellugi, U., Wang, P., Jernigan, T. (1994). Williams syndrome: an unusual neuropsychological profile. In: Broman, S., Grafman, J., editors. Atypical Cognitive Deficits in Developmental Disorders: Implications for Brain Function. Hillsdale, NJ: Erlbaum Press, pp. 23–56.

20. Benítez-Burraco A, Lattanzi W, Murphy E (2016). Language Impairments in ASD Resulting from a Failed Domestication of the Human Brain. Front Neurosci. 10:373.

21. Berger NI, Ingersoll B. A further investigation of goal-directed intention understanding in young children with autism spectrum disorders (2015). J Autism Dev Disord. 44(12):3204–14.

22. Bhatara A, Quintin EM, Levy B, Bellugi U, Fombonne E, Levitin DJ (2010). Perception of emotion in musical performance in adolescents with autism spectrum disorders. Autism Res. 3(5):214–25.

23. Bitsika V., Sharpley C. F., Andronicos N. M., Agnew L. L. (2015). Hypothalamus-pituitary-adrenal axis daily fluctuation, anxiety and age interact to predict cortisol concentrations in boys with an autism spectrum disorder. Physiol. Behav. 138, 200–207.

24. Bjørklund G, Meguid NA, El-Ansary A, El-Bana MA, Dadar M, Aaseth J, Hemimi M, Osredkar J, Chirumbolo S (2018). Diagnostic and Severity-Tracking Biomarkers for Autism Spectrum Disorder. J Mol Neurosci. 66(4):492–511.

25. Blomberg, S., Rosander, M., Andersson, G. (2006). Fears, hyperacusis and musicality in Williams’s syndrome. Research in Developmental Disabilities, 27, 668–80.

26. Bobeck EN, Gomes I, Pena D, Cummings KA, Clem RL, Mezei M, Devi LA (2017). The BigLEN-GPR171 Peptide Receptor System Within the Basolateral Amygdala Regulates Anxiety-Like Behavior and Contextual Fear Conditioning. Neuropsychopharmacology. 42(13):2527–2536.

27. Booth CA, Brown JT, Randall AD (2014). Neurophysiological modification of CA1 pyramidal neurons in a transgenic mouse expressing a truncated form of disrupted-in-schizophrenia 1. Eur J Neurosci. 39(7):1074–90.

28. Boyle EA, Li YI, Pritchard JK (2017). An expanded view of complex traits: from polygenic to omnigenic. Cell 169(7):1177–1186.

29. Bódizs R, Gombos F, Kovács I (2012). Sleep EEG fingerprints reveal accelerated thalamocortical oscillatory dynamics in Williams syndrome. Res Dev Disabil. 33(1):153–64.

30. Bonnel A, Mottron L, Peretz I, Trudel M, Gallun E, Bonnel AM (2003). Enhanced pitch sensitivity in individuals with autism: A signal detection analysis. Journal of Cognitive Neuroscience. 15(2):226– 235.

31. Bradshaw NJ, Porteous DJ. (2012). DISC1-binding proteins in neural development, signalling and schizophrenia. Neuropharmacology. 62(3):1230–41.

32. Brandon NJ, Sawa A. (2011). Linking neurodevelopmental and synaptic theories of mental illness through DISC1. Nat Rev Neurosci.18;12(12):707-22.

33. Bressler J, Mosley TH, Penman A, Gottesman RF, Windham BG, Knopman DS, Wruck LM, Boerwinkle E. (2017). Genetic variants associated with risk of Alzheimer’s disease contribute to cognitive change in midlife: The Atherosclerosis Risk in Communities Study. Am J Med Genet B Neuropsychiatr Genet. 174(3):269–282.

34. Brock, J. (2007).Language abilities in Williams syndrome: acritical review. Dev. Psychopathol. 19, 97–127.

35. Brock, J., Jarrold, C., Farran, E.K., Laws, G., Riby, D.M. (2007). Do children with Williams syndrome really have good vocabulary knowledge? Methods for comparing cognitive and linguistic abilities in developmental disorders. Clinical Linguistics & Phonetics, 21(9), 673–88.

36. Byars, S. G., Stearns, S. C., Boomsma, J. J. (2014). Opposite risk patterns for autism and schizophrenia are associated with normal variation in birth size: phenotypic support for hypothesized diametric gene-dosage effects. Proc. Biol. Sci. 281, 20140604.

37. Campbell LE, Daly E, Toal F, Stevens A, Azuma R, Karmiloff-Smith A, Murphy DG, Murphy KC (2009) Brain structural differences associated with the behavioural phenotype in children with Williams syndrome. Brain Res. 1258:96–107.

38. Campbell, D. J., Chang, J., Chawarska, K. (2014). Early generalized overgrowth in autism spectrum disorder: prevalence rates, gender effects, and clinical outcomes. J. Am. Acad. Child Adolesc. Psychiatry 53, 1063–73.

39. Carvalho CM, Vasanth S, Shinawi M (2014). Dosage changes of a segment at 17p13.1 lead to intellectual disability and microcephaly as a result of complex genetic interaction of multiple genes. Am J Hum Genet. 95(5):565–578

40. Charman, T., Swettenham, J., Baron-Cohen, S., Cox, A., Baird, G., Drew, A. (1997). Infants with autism: An investigation of empathy, pretend play, joint attention, and imitation. Developmental Psychology, 33, 781–789.

41. Charron, D. J., McDevitt, H. O. (1979). Analysis of HLA-D region-associated molecules with monoclonal antibody. Proc. Nat. Acad. Sci. 76: 6567–6571.

42. Chawarska, K., Campbell, D., Chen, L., Shic, F., Klin, A., Chang, J. (2011). Early generalized overgrowth in boys with autism. Arch. Gen. Psychiatry 68, 1021–31.

43. Chen L, Liu M, Luan Y, Liu Y, Zhang Z, Ma B, Liu X, Liu Y. (2018). BMP-6 protects retinal pigment epithelial cells from oxidative stress-induced injury by inhibiting the MAPK signaling pathways. Int J Mol Med. 42(2):1096–1105.

44. Chen EY, Tan CM, Kou Y, Duan Q, Wang Z, Meirelles GV, Clark NR, Ma’ayan A (2013) Enrichr: interactive and collaborative HTML5 gene list enrichment analysis tool. BMC Bioinformatics 14: 128

45. Cheetham ME, Caplan AJ (1998). Structure, function and evolution of DnaJ: conservation and adaptation of chaperone function. Cell Stress Chaperones. 3(1):28–36.

46. Cheng W, Rolls ET, Gu H, Zhang J, Feng J (2015). Autism: reduced connectivity between cortical areas involved in face expression, theory of mind, and the sense of self. Brain. 138(Pt 5):1382–93.

47. hiang MC, Reiss AL, Lee AD, Bellugi U, Galaburda AM, Korenberg JR, Mills DL, Toga AW, Thompson PM. (2007). 3D Pattern of brain abnormalities in Williams Syndrome visualized using tensor-based morphometry. NeuroImage 36.4: 1096–1109.

48. Chinen K, Takahashi E, Nakamura Y. (1996). Isolation and mapping of a human gene (SEC14L), partially homologous to yeast SEC14, that contains a variable number of tandem repeats (VNTR) site in its 3’ untranslated region. Cytogenet Cell Genet. 73(3):218–23.

49. Christ, S. E., Holt, D. D., White, D. A., Green, L. (2007). Inhibitory control in children with autism spectrum disorder. Journal of Autism and Developmental Disorders, 37(6), 1155–1165.

50. Colobran R, Pedrosa E, Carretero-Iglesia L, Juan M. (2010). Copy number variation in chemokine superfamily: the complex scene of CCL3L-CCL4L genes in health and disease. Clin Exp Immunol. 162(1):41–52.

51. Corbett BA, Schupp CW, Levine S, Mendoza S (2009). Comparing cortisol, stress, and sensory sensitivity in children with autism. Autism Res. 2(1):39–49.

52. Cornish K., Scerif G. Karmiloff-Smith A. (2007) Tracing syndrome-specific trajectories of attention across the lifespan. Cortex 43, 672–85.

53. Courchesne, E., Campbell, K., and Solso, S. (2011). Brain growth across the life span in autism: age-specific changes in anatomical pathology. Brain Res. 1380, 138–45.

54. Crabtree GW, Gogos JA (2014). Synaptic plasticity, neural circuits, and the emerging role of altered short-term information processing in schizophrenia. Front Synaptic Neurosci. 25;6:28.

55. Crespi B, Badcock C (2008). Psychosis and autism as diametrical disorders of the social brain. Behav Brain Sci 31:241–320.

56. Creutz CE, Tomsig JL, Snyder SL, Gautier MC, Skouri F, Beisson J, Cohen J. The copines, a novel class of C2 domain-containing, calcium-dependent, phospholipid-binding proteins conserved from Paramecium to humans. J Biol Chem. 1998 Jan 16;273(3):1393–402.

57. Critchley HD, Daly EM, Bullmore ET, Williams SC, Van Amelsvoort T, Robertson DM, Rowe A, Phillips M, McAlonan G, Howlin P, Murphy DG (2000). The functional neuroanatomy of social behaviour: changes in cerebral blood flow when people with autistic disorder process facial expressions. Brain. 123 (Pt 11):2203–12.

58. Dai L., Carter C. S., Ying J., Bellugi U., Pournajafi-Nazarloo H., Korenberg J. R. (2012). Oxytocin and vasopressin are dysregulated in Williams syndrome, a genetic disorder affecting social behavior. PLoS One 7:e38513.

59. Dalton KM, Nacewicz BM, Johnstone T, Schaefer HS, Gernsbacher MA, Goldsmith HH, Alexander AL, Davidson RJ (2005) Gaze fixation and the neural circuitry of face processing in autism. Nat Neurosci 8:519 –526.

60. Davenport EC, Szulc BR, Drew J, Taylor J, Morgan T, Higgs NF, López-Doménech G, Kittler JT (2019). Autism and Schizophrenia-Associated CYFIP1 Regulates the Balance of Synaptic Excitation and Inhibition. Cell Rep. 26(8):2037–2051.e6.

61. David, O., Maess, B., Eckstein, K., Friederici, A. D. (2011). Dynamic causal modeling of subcortical connectivity of language. The Journal of neuroscience: the official journal of the Society for Neuroscience, 31(7), 2712–2717. https://doi.org/10.1523/JNEUROSCI.3433-10.2011

62. Decety J., Jackson P. L. (2004). The functional architecture of human empathy. Behav. Cogn. Neurosci. Rev. 3, 71–100.

63. Dedovic K., Duchesne A., Andrews J., Engert V., Pruessner J. C. (2009). The brain and the stress axis: the neural correlates of cortisol regulation in response to stress. Neuroimage 47 864–871.

64. De Rubeis S, He X, Goldberg AP, Poultney CS, Samocha K, Cicek AE, Kou Y, LiuL Fromer M, Walker S, Singh T, Klei L, Kosmicki J, Shih-Chen F, Aleksic B, Biscaldi M, Bolton PF, Brownfeld JM, Cai J, Campbell NG, Carracedo A, Chahrour MH, Chiocchetti AG, Coon H, Crawford EL, Curran SR, Dawson G, Duketis E, Fernandez BA, Gallagher L, Geller E, Guter SJ, Hill RS, Ionita-Laza J, JimenzGonzalez P, Kilpinen H, Klauck SM, Kolevzon A, Lee I, Lei I, Lei J, Lehtimäki T, Lin CF, Ma’ayan A, Marshall CR, McInnes AL, Neale B, Owen MJ, Ozaki N, Parellada M, Parr JR, Purcell S, Puura K, Rajagopalan D, Rehnström K, Reichenberg A, SaboA Sachse M, Sanders SJ, Schafer C, Schulte-Rüther M, Skuse D, Stevens C, Szatmari P, Tammimies K, Valladares O, Voran A, Li-San W, Weiss LA, Willsey AJ, Yu TW, Yuen RK; DDD Study; Homozygosity Mapping Collaborative for Autism; UK10KConsortium, Cook EH, Freitag CM, Gill M, Hultman CM, Lehner T, Palotie A, Schellenberg GD, Sklar P, State MW, Sutcliffe JS, Walsh CA, Scherer SW, Zwick ME, Barett JC, Cutler DJ, Roeder K, Devlin B, Daly MJ, Buxbaum JD (2014). Synaptic,transcriptional and chromatin genes disrupted in autism. Nature. 515(7526):209-15.

65. Deurloo MHS, Turlova E, Chen WL, Lin YW, Tam E, Tassew NG, Wu M, Huang YC, Crawley JN, Monnier PP, Groffen AJA, Sun HS, Osborne LR, Feng ZP (2019). Transcription Factor 2I Regulates Neuronal Development via TRPC3 in 7q11.23 Disorder Models. Mol Neurobiol. 56(5):3313–3325.

66. Dewey, D., Cantell, M., Crawford, S.G. (2007). Motor and gestural performance in children with autism spectrum disorders, developmental coordination disorder, and/or attention deficit hyperactivity disorder. Journal of the International Neuropsychological Society, 13, 246–256.

67. Diez-Itza E, Martínez V, Antón A (2016). Narrative competence in Spanish-speaking adults with Williams syndrome. Psicothema. 28(3):291–7.

68. Dichter GS, Damiano CA, Allen JA (2012). Reward circuitry dysfunction in psychiatric and neurodevelopmental disorders and genetic syndromes: animal models and clinical findings. J Neurodev Disord. 64(1):19.

69. Dichter GS, Felder JN, Green SR, Rittenberg AM, Sasson NJ, Bodfish JW (2012a). Reward circuitry function in autism spectrum disorders. Soc Cogn Affect Neurosci. 7:160–72.

70. Dimova, D., Dyson, N (2005). The E2F transcriptional network: old acquaintances with new faces. Oncogene 24, 2810–2826.

71. Doherty-Sneddon G, Riby DM, Whittle L (2012). Gaze aversion as a cognitive load management strategy in autism spectrum disorder and Williams syndrome. J Child Psychol Psychiatry. 53(4):420–30.

72. Doll A., Grzeschik K. H. (2001). Characterization of two novel genes, WBSCR20 and WBSCR22, deleted in Williams–Beuren syndrome.Cytogenet. Cell Genet. 95, 20–27.

73. Domes G., Heinrichs M., Kumbier E., Grossmann A., Hauenstein K., Herpertz S. C. (2013). Effects of intranasal oxytocin on the neural basis of face processing in autism spectrum disorder. Biol. Psychiatry 74 164–171.

74. Dong D, Zielke HR, Yeh D, Yang P (2018). Cellular stress and apoptosis contribute to the pathogenesis of autism spectrum disorder. Autism Res. 11(7):1076–1090.

75. Doyle, T.F., Bellugi, U., Korenberg, J.R., Graham, J. (2004). ‘‘Everybody in the world is my friend’’: hypersociability in young children with Williams Syndrome. American Journal of Medical Genetics, 124A, 263–73.

76. Dykens, E. M. (2003). Anxiety, fears and phobias in persons with Williams syndrome. Developmental Neuropsychology, 23, 291–316.

77. Elliott D., Welsh T. N., Lyons J., Hansen S. Wu M. (2006) The visual regulation of goal-directed reaching movements in adults with Williams syndrome, Down syndrome, and other developmental delays. Motor Control 10, 34–54.

78. Fachim HA, Srisawat U, Dalton CF, Reynolds GP (2018). Parvalbumin promoter hypermethylation in postmortem brain in schizophrenia. Epigenomics. 10(5):519–524.

79. Fein, D., Dunn, M.A., Allen, D. M., Aram, R., Hall, N., Morris, R. (1996). Neuropsychological and language findings In: Rapin I, editor. Preschool Children with Inadequate Communication: Developmental Language Disorder, Autism, Low IQ. 123–154. London: MacKeith Press.

80. Firth, U., Snowling, M. (1983). Reading for meaning and reading for sound in autistic and dyslexic children. Journal of Developmental Psychology, 1, 329–42.

81. Fishman I, Yam A, Bellugi U, Lincoln A, Mills D (2011). Contrasting patterns of language-associated brain activity in autism and Williams syndrome. Soc Cogn Affect Neurosci. 6(5):630–8.

82. Fountain MD, Oleson DS, Rech ME, Segebrecht L, Hunter JV, McCarthy JM, Lupo PJ, Holtgrewe M, Moran R, Rosenfeld JA, Isidor B, Le Caignec C, Saenz MS, Pedersen RC, Morgan TM, Pfotenhauer JP, Xia F, Bi W, Kang SL, Patel A, Krantz ID, Raible SE, Smith W, Cristian I, Torti E, Juusola J, Millan F, Wentzensen IM, Person RE, Küry S, Bézieau S, Uguen K, Férec C, Munnich A, van Haelst M, Lichtenbelt KD, van Gassen K, Hagelstrom T, Chawla A, Perry DL, Taft RJ, Jones M, Masser-Frye D, Dyment D, Venkateswaran S, Li C, Escobar LF, Horn D, Spillmann RC, Peña L, Wierzba J, Strom TM, Parenti I, Kaiser FJ, Ehmke N, Schaaf CP (2019). Pathogenic variants in USP7 cause a neurodevelopmental disorder with speech delays, altered behavior, and neurologic anomalies. Genet Med. 21(8):1797–1807.

83. Freeman, L.; Dake, S (1996). Teach me language: A language manual for children with autism, Asperger’s syndrome and related developmental disorders. Langley, British Columbia, Canada.

84. Freese, J. L., Pino, D., Pleasure, S. J. (2010). Wnt signaling in development and disease. Neurobiology of disease, 38(2), 148–153.

85. Frigerio E, Burt DM, Gagliardi C, Cioffi G, Martelli S, Perrett DI (2006). Is everybody always my friend? Perception of approachability in Williams syndrome. Neuropsychologia. 44:254–259.

86. Fujiwara S, Matsui TS, Ohashi K, Mizuno K, Deguchi S (2019). Keratin-binding ability of the N-terminal Solo domain of Solo is critical for its function in cellular mechanotransduction. Genes Cells. 24(5):390–402.

87. Fukumoto, A., Hashimoto, T., Ito, H., Nishimura, M., Tsuda, Y., Miyazaki, M. (2008). Growth of head circumference in autistic infants during the first year of life. J. Autism Dev. Disord. 38, 411–8.

88. Gagliardi C., Martelli S., Burt M. D. Borgatti R. (2007) Evolution of neurologic features in Williams syndrome. Pediatric Neurology 36, 301–6.

89. Gallese V., Goldman A. (1998). Mirror neurons and the simulation theory of mind-reading. Trends Cogn. Sci. 2, 493–501.

90. Gallese V (2001). The’ shared manifold’ hypothesis. From mirror neurons to empathy. Journal of Consciousness Studies. 8:5–7.

91. Gao Y, Wang J, Shangguan S, Bao Y, Lu X, Zou J, Dai Y, Liu J, Zhang T. (2018). Quantitative Measurement of PARD3 Copy Number Variations in Human Neural Tube Defects. Cell Mol Neurobiol. 38(3):605–614.

92. Garland P, Quraishe S, French P, O’Connor V. (2008). Expression of the MAST family of serine/threonine kinases. Brain Res. 8 Feb 21;1195:12-9.

93. Gastgeb HZ, Strauss MS, Minshew NJ. (2006). Do individuals with autism process categories differently? The effect of typicality and development. Child Dev. 77:1717–1729.

94. Geschwind, G.H., State, M.W. (2015). Gene hunting in autism spectrum disorder: on the path to precision medicine. Lancet Neurol. 14, 1109–1120.

95. Gesundheit B, Rosenzweig JP, Naor D, Lerer B, Zachor DA, Procházka V, Melamed M, Kristt DA, Steinberg A, Shulman C, Hwang P, Koren G, Walfisch A, Passweg JR, Snowden JA, Tamouza R, Leboyer M, Farge-Bancel D, Ashwood P (2013). Immunological and autoimmune considerations of Autism Spectrum Disorders. J Autoimmun. 44:1–7.

96. Ghaffari M, Tahmasebi Birgani M, Kariminejad R, Saberi A (2018) Genotype-phenotype correlation and the size of microdeletion or microduplication of 7q11.23 region in patients with Williams-Beuren syndrome. Ann Hum Genet. 82(6):469–476.

97. Gibbard CR, Ren J, Skuse DH, Clayden JD, Clark CA (2018). Structural connectivity of the amygdala in young adults with autism spectrum disorder. Hum Brain Mapp. 39(3):1270–1282.

98. Gillberg C, Rasmussen P (1994). Brief report: four case histories and a literature review of Williams syndrome and autistic behavior. J Autism Dev Disord. 24(3):381–393.

99. Gilbert, J., Man, H. Y. (2017). Fundamental Elements in Autism: From Neurogenesis and Neurite Growth to Synaptic Plasticity. Frontiers in cellular neuroscience, 11, 359.

100. Gluderer S, Brunner E, Germann M, Jovaisaite V, Li C, Rentsch CA, Hafen E, Stocker H (2010). Madm (Mlf1 adapter molecule) cooperates with Bunched A to promote growth in Drosophila. J Biol. 9(1):9.

101. Goddard, M.N., van Rijn, S., Rombouts, S.A.R.B (2016). White matter microstructure in a genetically defined group at increased risk of autism symptoms, and a comparison with idiopathic autism: an exploratory study. Brain Imaging and Behavior 10, 1280–1288.

102. Golarai G., Hong S., Haas B. W., et al. (2010). The fusiform face area is enlarged in Williams syndrome. J. Neurosci. 19 6700–6712.

103. Goldstein, G., Minshew, N.J., Allen, D.N., Seaton, B.E. (2002). High-functioning autism and schizophrenia: a comparison of an early and late onset neurodevelopmental disorder. Archives of Clinical Neuropsychology, 17(5), 461–75.

104. Gonzalez-Brito MR, Bixby JL (2009). Protein tyrosine phosphatase receptor type O regulates development and function of the sensory nervous system. Mol Cell Neurosci. 42(4):458–65.

105. Goriounov, D., Leung, C. L., Liem, R. K. H. (2003). Protein products of human Gas2-related genes on chromosomes 17 and 22 (hGAR17 and hGAR22) associate with both microfilaments and microtubules. J. Cell Sci. 116: 1045–1058.

106. Gothelf, D., Searcy, Y.M., Reilly, J., et al. (2008). Association between cerebral shape and social use of language in Williams syndrome. American Journal of Medical Genetics Part A, 146A, 2753–61.

107. Gotts SJ, Simmons WK, Milbury LA, Wallace GL, Cox RW, Martin A (2012). Fractionation of social brain circuits in autism spectrum disorders. Brain. 135:2711–25.

108. Grabowski M, Zimprich A, Lorenz-Depiereux B, Grabowski M, Zimprich A, Lorenz-Depiereux B, Kalscheuer V, Asmus F, Gasser T, Meitinger T, Strom T. (2003). The epsilon-sarcoglycan gene (SGCE), mutated in myoclonus-dystonia syndrome, is maternally imprinted. Eur J Hum Genet. 11(2):138–144.

109. Graham, J. M., Rosner, B., Dykens, E., Visootsak, J. (2005). Behavioral features of CHARGE syndrome (Hall-Hittner syndrome)comparison with Down syndrome, Prader-Willi syndrome, and Williams syndrome. American Journal of Medical Genetics, 133, 240–247.

110. Guo H, Hirschhorn J, Dauber A (2018). Insights and Implications of Genome-Wide Association Studies of Height. The Journal of Clinical Endocrinology & Metabolism. 103; 9, 3155–3168.

111. Gyawali S, Patra BN (2019). Autism spectrum disorder: Trends in research exploring etiopathogenesis. Psychiatry Clin Neurosci. 73(8):466–475.

112. Haas B.W., Hoeft F., Searcy Y. M., Mills D., Bellugi U. Reiss A. (2010) Individual differences in social behavior predict amygdala response to fearful facial expressions in Williams syndrome. Neuropsychologia 48, 1283–8.

113. Haas BW, Sheau K, Kelley RG, Thompson PM, Reiss AL (2012). Regionally specific increased volume of the amygdala in Williams’s syndrome: Evidence from surface-based modeling. Hum Brain Mapp.

114. Haas BW, Mills D, Yam A, Hoeft F, Bellugi U, Reiss A (2009). Genetic influences on sociability: Heightened amygdala reactivity and event-related responses to positive social stimuli in Williams syndrome. J. Neurosci. 29:1132–1139.

115. Haas, Brian W, and Allan L Reiss (2012). Social brain development in Williams syndrome: the current status and directions for future research. Frontiers in Psychology 3 186.

116. Haas B, Barnea-Goraly M, Sheau K, Yamagata B, Ullas S, Reiss A (2013). Altered Microstructure Within Social-Cognitive Brain Networks During Childhood in Williams Syndrome. Cerebral Cortex.24; 10,2796–2806.

117. Haas B. W., Sheau K., Kelley R. G., Thompson P. M., Reiss A. L. (2014). Regionally specific increased volume of the amygdala in Williams syndrome: evidence from surface-based modeling. Hum. Brain Mapp. 35, 866–874.

118. Haas BW, Smith AK. (2015). Oxytocin, vasopressin, and Williams syndrome: epigenetic effects on abnormal social behavior. Front Genet. 6:28.

119. Hadjikhani N, Joseph RM, Snyder J, Tager-Flusberg H (2006). Anatomical differences in the mirror neuron system and social cognition network in autism. Cerebral Cortex 16:1276–1282.

120. Hale, C. M. Tager-Flusberg, H. (2005). Social communication in children with autism:the relationship between theory of mind and discourse development. Autism 9, 157–178.

121. Hampson DR, Blatt GJ (2015). Autism spectrum disorders and neuropathology of the cerebellum. Front Neurosci. 9:420.

122. Hapak SM, Rothlin CV, Ghosh S (2018). PAR3-PAR6-atypical PKC polarity complex proteins in neuronal polarization. Cell Mol Life Sci. 75(15):2735–2761.

123. Happé F.G.E. (1997). Central coherence and theory of mind in autism: reading homographs in context. British Journal of Developmental Psychology, 15, 1–12.

124. Happé F.G.E. (1995).The role of age and verbal ability in the theory of mind task performance of subjects with autism. Child Dev. 66, 843–855.

125. Happé F.G.E. (1993). Communicative competence and theory of mind in autism: a test ofrelevance theory. Cognition. 48(2):101–19.

126. Happé F, Frith U (1996). The neuropsychology of autism. Brain: A Journal of Neurology. 119:1377– 1400.

127. Hashemi E, Ariza J, Rogers H, Noctor SC, Martínez-Cerdeño V (2017). The Number of Parvalbumin-Expressing Interneurons Is Decreased in the Prefrontal Cortex in Autism. Cereb Cortex. 1;27(3):1931-1943.

128. Harris, G.J., Chabris, C.F., Clark, J., et al. (2006). Brain activation during semantic processing in autism spectrum disorders via function al magnetic imaging. Brain Cognition, 61, 54–68.

129. Haxby J. V., Hoffman E. A., Gobbini M. I. (2000). The distributed human neural system for face perception. Trends Cogn. Sci. 4, 223–233.

130. Heaton P, Hermelin B, Pring L (1998). Autism and pitch processing: A precursor for savant musical ability. Music Perception. 15(3):291–305.

131. Henrichsen CN, Csárdi G, Zabot MT, Fusco C, Bergmann S, Merla G, Reymond A (2011). Using transcription modules to identify expression clusters perturbed in Williams-Beuren syndrome. PLoS Comput Biol. 7(1):e1001054.

132. Heaton P (2003). Pitch memory, labelling and disembedding in autism. Journal of Child Psychology and Psychiatry. 44(4):543–551.

133. Hohman TJ, Koran ME, Thornton-Wells TA (2014). Alzheimer’s Neuroimaging Initiative. Genetic variation modifies risk for neurodegeneration based on biomarker status. Front Aging Neurosci. 4;6:183.

134. Hollander E., Bartz J., Chaplin W., Phillips A., Sumner J., Soorya L., et al. (2007). Oxytocin increases retention of social cognition in autism. Biol. Psychiatry 61 498–50310.1016/j.biopsych.2006.05.030

135. Homman-Ludiye J, Kwan WC, de Souza MJ, Rodger J, Bourne JA. (2017). Ephrin-A2 regulates excitatory neuron differentiation and interneuron migration in the developing neocortex. Sci Rep. 18;7(1):11813.

136. Humphray SJ, Oliver K, Hunt AR, Plumb RW, Loveland JE, Howe KL, Andrews TD, Searle S, Hunt SE, Scott CE, Jones MC, Ainscough R, Almeida JP, Ambrose KD, Ashwell RI, Babbage AK, Babbage S, Bagguley CL, Bailey J, Banerjee R, Barker DJ, Barlow KF, Bates K, Beasley H, Beasley O, Bird CP, Bray-Allen S, Brown AJ, Brown JY, Burford D, Burrill W, Burton J, Carder C, Carter NP, Chapman JC, Chen Y, Clarke G, Clark SY, Clee CM, Clegg S, Collier RE, Corby N, Crosier M, Cummings AT, Davies J, Dhami P, Dunn M, Dutta I, Dyer LW, Earthrowl ME, Faulkner L, Fleming CJ, Frankish A, Frankland JA, French L, Fricker DG, Garner P, Garnett J, Ghori J, Gilbert JG, Glison C, Grafham DV, Gribble S, Griffiths C, Griffiths-Jones S, Grocock R, Guy J, Hall RE, Hammond S, Harley JL, Harrison ES, Hart EA, Heath PD, Henderson CD, Hopkins BL, Howard PJ, Howden PJ, Huckle E, Johnson C, Johnson D, Joy AA, Kay M, Keenan S, Kershaw JK, Kimberley AM, King A, Knights A, Laird GK, Langford C, Lawlor S, Leongamornlert DA, Leversha M, LloydC Lloyd DM, Lovell J, Martin S, Mashreghi-Mohammadi M, Matthews L, McLaren S, McLay KE, McMurray A, Milne S, Nickerson T, Nisbett J, Nordsiek G, Pearce AV, Peck AI, Porter KM, Pandian R, Pelan S, Phillimore B, Povey S, Ramsey Y, Rand V, Scharfe M, Sehra HK, Shownkeen R, Sims SK, Skuce CD, Smith M, Steward CA, Swarbreck D, Sycamore N, Tester J, Thorpe A, Tracey A, Tromans A, Thomas DW, WallM Wallis JM, West AP, Whitehead SL, Willey DL, Williams SA, Wilming L, Wray PW, Young L, Ashurst JL, Coulson A, Blöcker H, Durbin R, Sulston JE, Hubbard T, Jackson MJ, Bentley DR, Beck S, Rogers J, Dunham I (2004). DNA sequence and analysis of human chromosome 9. Nature. 429(6990):369–74.

137. Hwang JY, Yan J, Zukin RS (2019). Autophagy and synaptic plasticity: epigenetic regulation. Curr Opin Neurobiol. 59:207–212.

138. Iacoboni M (2009). Imitation, empathy, and mirror neurons. Annual Review of Psychology. 60:653– 670.

139. Ikeda, W., Nakanishi, H., Miyoshi, J., Mandai, K., Ishizaki, H., Tanaka, M., Togawa, A., Takahashi, K., Nishioka, H., Yoshida, H., Mizoguchi, A., Nishikawa, S., Takai, Y. (1999). Afadin: A key molecule essential for structural organization of cell-cell junctions of polarized epithelia during embryogenesis. The Journal of cell biology, 146(5), 1117–1132.

140. Ikeda M, Yoshino M (2018). Nitric oxide augments single persistent Na(+) channel currents via the cGMP/PKG signaling pathway in Kenyon cells isolated from cricket mushroom bodies. J Neurophysiol. 120(2):720–728.

141. Ingersoll B, Walton K, Carlsen D, Hamlin T. (2013). Social intervention for adolescents with autism and significant intellectual disability: initial efficacy of reciprocal imitation training. Am J Intellect Dev Disabil. 118(4):247–61.

142. Isozaki Y, Sakai K, Kohiro K, Kagoshima K, Iwamura Y, Sato H, Rindner D, Fujiwara S, Yamashita K, Mizuno K, Ohashi K (2020). The Rho-guanine nucleotide exchange factor Solo decelerates collective cell migration by modulating the Rho-ROCK pathway and keratin networks. Mol Biol Cell. mbcE19070357.

143. Jackowski AP, Rando K, Maria de Araújo C, Del Cole CG, Silva I, Tavares de Lacerda AL. (2009). Brain abnormalities in Williams syndrome: a review of structural and functional magnetic resonance imaging findings. Eur J Paediatr Neurol. 13(4):305–16.

144. Jakobson CM, Jarosz DF (2019) Molecular origins of complex heritability in natural genotype-to-phenotype relationships. Cell Syst. 8(5):363–379.

145. Jansiewicz EM, Goldberg MC, Newschaffer CJ, Denckla MB, Landa R, Mostofsky SH (2006). Motor signs distinguish children with high functioning autism and Asperger’s syndrome from controls. J Autism Dev Disord. 36(5):613–21.

146. Järvinen A, Ng R, Bellugi U (2015). Autonomic response to approachability characteristics, approach behavior, and social functioning in Williams syndrome. Neuropsychologia. 78:159–70.

147. Järvinen-Pasley A, Wallace GL, Ramus F, Happé F, Heaton P (2008). Enhanced perceptual processing of speech in autism. Dev Sci. 11(1):109–21.

148. Järvinen A, Korenberg JR, Bellugi U (2013). The social phenotype of Williams syndrome. Curr Opin Neurobiol. 23(3):414–22.

149. Jawaid A, Riby DM, Owens J, White SW, Tarar T, Schulz PE (2012). ’Too withdrawn’ or ‘too friendly’: considering social vulnerability in two neuro-developmental disorders. J Intellect Disabil Res. 56(4):335–50.

150. Jernigan TL, Bellugi U. (1990). Anomalous brain morphology on magnetic resonance images in Williams syndrome and Down syndrome. Arch Neurol. 47(5):529–33.

151. Jernigan TL, Bellugi U, Sowell E, Doherty S, Hesselink JR. (1993) Cerebral morphologic distinctions between Williams and Down syndromes. Arch Neurol. 50:186–91.

152. John A. E., Rowe M. L., Mervis C. B. (2009). Referential communication skills of children with Williams syndrome: understanding when messages are not adequate, Am. J. Intellect. Dev. Disabil. 114, 85–99 10.1352/2009.114.85-99.

153. Joensuu M, Lanoue V, Hotulainen P (2018). Dendritic spine actin cytoskeleton in autism spectrum disorder. Prog Neuropsychopharmacol Biol Psychiatry. 8;84(Pt B):362-81.

154. Jung, E. M., Moffat, J. J., Liu, J., Dravid, S. M., Gurumurthy, C. B., Kim, W. Y. (2017). Arid1b haploinsufficiency disrupts cortical interneuron development and mouse behavior. Nature neuroscience, 20(12), 1694–1707.

155. Kana R, Keller T, Cherkassky V, Minshew N, Just M. (2006). Sentence comprehension in autism: thinking in pictures with decreased functional connectivity. Brain. 129; 9,2484–2493.

156. Karmiloff-Smith A, Broadbent H, Farran EK, Longhi E, D’Souza D, Metcalfe K, Tassabehji M, Wu R, Senju A, Happé F, Turnpenny P, Sansbury F (2012) Social cognition in williams syndrome: genotype/phenotype insights from partial deletion patients. Front Psychol. 3:168.

157. Kelley E, Paul J, Fein D, Naigles L (2006). Residual Language Deficits in Optimal Outcome Children with a History of Autism. J Autism Dev Disord. 36:807–828.

158. Keil R, Schulz J, Hatzfeld M (2012). p0071/PKP4, a multifunctional protein coordinating cell adhesion with cytoskeletal organization. Biol Chem. 394(8):1005–17.

159. Kennedy, D. P., Adolphs, R. (2012). The social brain in psychiatric and neurological disorders. Trends in cognitive sciences, 16(11), 559–572.

160. Klin A, Sparrow SS, de Bildt A, Cicchetti DV, Cohen DJ, Volkmar FR (1999). A normed study of face recognition in autism and related disorders. J Autism Dev Disord. 29(6):499–508.

161. Kimura R, Swarup V, Tomiwa K, Gandal MJ, Parikshak NN, Funabiki Y, Nakata M, Awaya T, Kato T, Iida K, Okazaki S, Matsushima K, Kato T, Murai T, Heike T, Geschwind DH, Hagiwara M. (2019) Integrative network analysis reveals biological pathways associated with Williams syndrome. J Child Psychol Psychiatry. 60(5):585–598.

162. Kjelgaard MM, Tager-Flusberg H. (2001). An Investigation of Language Impairment inAutism: Implications for Genetic Subgroups. Lang Cogn Process. 16(2-3):287–308.

163. Klein AJ, Armstrong BL, Greer MK, Brown FR 3rd (1990). Hyperacusis and otitis media in individuals with Williams syndrome. J Speech Hear Disord. 55(2):339–44.

164. Klein-Tasman BP, Phillips KD, Lord C, Mervis CB, Gallo FJ (2009). Overlap with the autism spectrum in young children with Williams syndrome. J Dev Behav Pediatr. 30(4):289–99.

165. Klein-Tasman BP, van der Fluit F, Mervis CB. (2018). Autism Spectrum Symptomatology in Children with Williams Syndrome Who Have Phrase Speech or Fluent Language. JAutism Dev Disord. 48(9):3037-3050.

166. Klein-Tasman BP, Mervis CB, Lord C, Phillips KD (2007). Socio-communicative deficits in young children with Williams syndrome: performance on the Autism Diagnostic Observation Schedule. Child Neuropsychol. 13(5):444–67.

167. Kliemann D, Dziobek I, Hatri A, Baudewig J, Heekeren HR (2012). The role of the amygdala in atypical gaze on emotional faces in autism spectrum disorders. J Neurosci. 32(28):9469–76.

168. Korpilahti P, Jansson-Verkasalo E, Mattila ML, Kuusikko S, Suominen K, Rytky S, Pauls DL, Moilanen I (2007). Processing of affective speech prosody is impaired in Asperger syndrome. J Autism Dev Disord 37(8):1539–49.

169. Kotz SA, Schwartze M, Schmidt-Kassow M (2009) Non-motor basal ganglia functions: a review and proposal for a model of sensory predictability in auditory language perception. Cortex. 45(8):982–90

170. Kuang S (2016). Two Polarities of Attention in Social Contexts: From Attending-to-Others to Attending-to-Self. Front Psychol. 7:63.

171. Kuleshov MV, Jones MR, Rouillard AD, Fernandez NF, Duan Q, Wang Z, Koplev S, Jenkins SL, Jagodnik KM, Lachmann A, McDermott MG, Monteiro CD, Gundersen GW, Ma’ayan A (2016) Enrichr: a comprehensive gene set enrichment analysis web server 2016 update. Nucleic Acids Research. 44(W1):W90–7

172. Korenberg JR, Chen XN, Hirota H, Lai Z, Bellugi U, Burian D, Roe B, Matsuoka R (2000). Genome structure and cognitive map of Williams syndrome. J Cogn Neurosci.12 Suppl 1:89-107.

173. Lacroix, A., Famelart, N., Guidetti, M. (2016). Language and emotional abilities in children with Williams syndrome and children with autism spectrum disorder: similarities and differences. Pediatric health, medicine and therapeutics, 7, 89–97.

174. Lacroix A, Guidetti M, Rogé B, Reilly J. (2009). Recognition of emotional and nonemotional facial expressions: a comparison between Williams syndrome and autism. Res Dev Disabil. 30(5):976–85.

175. Lalli MA, Jang J, Park JH, Wang Y, Guzman E, Zhou H, Audouard M, Bridges D, Tovar KR, Papuc SM, Tutulan-Cunita AC, Huang Y, Budisteanu M, Arghir A, Kosik KS. (2016). Haploinsufficiency of BAZ1B contributes to Williams syndrome through transcriptional dysregulation of neurodevelopmental pathways. Hum Mol Genet 25: 1294–306.

176. Lauber, E., Filice, F., Schwaller, B. (2018). Dysregulation of Parvalbumin Expression in the Cntnap2-/-Mouse Model of Autism Spectrum Disorder. Frontiers in molecular neuroscience, 11, 262.

177. Laws G, Bishop D. (2004) Pragmatic language impairment and social deficits in Williams syndrome: a comparison with Down’s syndrome and specific language impairment. Int J Lang Commun Disord. 39(1):45–64.

178. Lee JA (2012). Neuronal autophagy: a housekeeper or a fighter in neuronal cell survival?. Exp Neurobiol. 21(1):1–8.

179. Lense, M. D., Dykens, E. M. (2013). Cortisol reactivity and performance abilities in social situations in adults with Williams syndrome. American journal on intellectual and developmental disabilities, 118(5), 381–393.

180. Levy-Shraga Y, Gothelf D, Pinchevski-Kadir S, Katz U, Modan-Moses D (2018). Endocrine manifestations in children with Williams-Beuren syndrome. Acta Paediatr 107(4):678–684.

181. Lieberman HR (2007). Cognitive methods for assessing mental energy. Nutr Neurosci. 10(5-6):229–42.

182. Lincoln A, Lai Z, Jones W (2002). Shifting attention and joint attention dissociation in Williams síndrome: Implications for the cerebellum and social déficits in autism. Neurocase. 8(3), 226–232.

183. Little K, Riby DM, Janes E, Clark F, Fleck R, Rodgers J (2013). Heterogeneity of social approach behaviour in Williams syndrome: the role of response inhibition. Res Dev Disabil. 34(3):959–67.

184. Liu X, Li Z, Fan C, Zhang D, Chen J. (2017). Genetics implicate common mechanisms in autism and schizophrenia: synaptic activity and immunity. J Med Genet. 54(8):511–520.

185. Liu Y, Harding M, Pittman A, Dore J, Striessnig J, Rajadhyaksha A, Chen X (2014). Cav1.2 and Cav1.3 L-type calcium channels regulate dopaminergic firing activity in the mouse ventral tegmental area. J Neurophysiol 112:1119–30.

186. Lombardo M, Chakrabarti B, Bullmore E, Sadek S, Pasco G, Wheelwright S, Suckling J, MRC AIMS Consortium, Baron-Cohen S (2010). Atypical neural self-representation in autism. Brain. 133;2, 611–624.

187. Lopes JS, Abril-de-Abreu R, Oliveira RF (2015). Brain Transcriptomic Response to Social Eavesdropping in Zebrafish (Danio rerio). PLoS One. 10(12):e0145801

188. Lough E, Hanley M, Rodgers J, South M, Kirk H, Kennedy DP, Riby DM (2015). Violations of Personal Space in Young People with Autism Spectrum Disorders and Williams Syndrome: Insights from the Social Responsiveness Scale. J Autism Dev Disord. 45(12):4101–8.

189. Lukács A, Csaba P, Racsmány M (2004). Language in Hungarian Children with Williams syndrome. Language Acquisition and Language Disorders. 36; 187-220.

190. Lynch CJ, Uddin LQ, Supekar K, Khouzam A, Phillips J, Menon V (2013). Default mode network in childhood autism: posteromedial cortex heterogeneity and relationship with social deficits. Biol Psychiatry. 74(3):212–9.

191. Malenfant, P., Liu, X., Hudson, M. L., Qiao, Y., Hrynchak, M., Riendeau, N., et al. (2012). Association of GTF2i in the Williams-Beuren syndrome critical region with autism spectrum disorders. J. Autism Dev. Disord. 42, 1459–69

192. Mariën P, Borgatti R. (2018) Language and the cerebellum. Handb Clin Neurol. 154:181–202.

193. Martens MA, Wilson SJ, Dudgeon P, Reutens DC (2009). Approachability and the amygdala: insights from Williams syndrome. Neuropsychologia. 47(12):2446–53.

194. Martens M. A., Wilson S. J., Reutens D. C. (2008). Research review: Williams syndrome: a critical review of the cognitive, behavioral, and neuroanatomical phenotype. J. Child. Psychol. Psychiatry 49, 576–608.

195. Martin LA, Iceberg E, Allaf G (2017). Consistent hypersocial behavior in mice carrying a deletion of Gtf2i but no evidence of hyposocial behavior with Gtf2i duplication: Implications for Williams-Beuren syndrome and autism spectrum disorder. Brain Behav. 8(1):e00895.

196. Martínez-Cerdeño V. (2017). Dendrite and spine modifications in autism and related neurodevelopmental disorders in patients and animal models. Developmental Neurobiology, 77(4), 393–404.

197. Matson JL, Nebel-Schwalm MS (2007). Comorbid psychopathology with autism spectrum disorder in children: an overview. Res Dev Disabil. 28(4):341–52.

198. Maud C, Ryan J, McIntosh JE, Olsson CA (2018). The role of oxytocin receptor gene (OXTR) DNA methylation (DNAm) in human social and emotional functioning: a systematic narrative review. BMC Psychiatry. 18(1):154.

199. McAlonan GM, Suckling J, Wong N, Cheung V, Lienenkaemper N, Cheung C, Chua SE (2008). Distinct patterns of grey matter abnormality in high-functioning autism and Asperger’s syndrome. J Child Psychol Psychiatry. 49(12):1287–95.

200. McQuillin A, Sharp SI. (2018). Rare variant analysis in multiply affected families, association studies and functional analysis suggest a role for the ITG 4 gene in schizophrenia and bipolar disorder. Schizophr Res. 199:181–188.

201. Meng Y, Zhang Y, Tregoubov V, Janus C, Cruz L, Jackson M, Lu WY, MacDonald JF, Wang JY, Falls DL, Jia Z (2002). Abnormal spine morphology and enhanced LTP in LIMK-1 knockout mice. Neuron. 35(1):121–33.

202. Merla G., Ucla C., Guipponi M., Reymond A. (2002). Identification of additional transcripts in the Williams–Beuren syndrome critical region.Hum. Genet.110, 429–438.

203. Mervis CB, Klein-Tasman BP, Mastin ME (2001). Adaptive behavior of 4-through 8-year-old children with Williams syndrome. Am J Ment Retard. 106(1):82–93.

204. Mervis CB, Becerra AM (2007). Language and communicative development in Williams syndrome. Ment Retard Dev Disabil Res Rev. 13(1):3–15.

205. Mervis CB, Kistler DJ, John AE, Morris CA. (2012) Longitudinal assessment of intellectual abilities of children with Williams syndrome: multilevel modeling of performance on the Kaufman Brief Intelligence Test-Second Edition. Am J Intellect Dev Disabil. 117(2):134–55.

206. Meyer-Lindenberg A, Hariri AR, Munoz KE, Mervis CB, Mattay VS, Morris CA, Berman KF (2005). Neural correlates of genetically abnormal social cognition in Williams syndrome. Nat Neurosci. 8(8):991–3.

207. Mills JL, Hediger ML, Molloy CA, Chrousos GP, Manning-Courtney P, Yu KF, Brasington M, England LJ (2007).. Elevated levels of growth-related hormones in autism and autism spectrum disorder. Clin Endocrinol (Oxf). 67(2):230–7.

208. Mimura M, Hoeft F, Kato M, Kobayashi N, Sheau K, Piggot J, Mills D, Galaburda A, Korenberg JR, Bellugi U, Reiss AL (2010). A preliminary study of orbitofrontal activation and hypersociability in Williams Syndrome. J Neurodev Disord. 2(2):93–98.

209. Mobbs D, Eckert MA, Mills D, Korenberg J, Bellugi U, Galaburda AM, Reiss AL (2007). Frontostriatal dysfunction during response inhibition in Williams syndrome. Biol Psychiatry. 62(3):256–61.

210. Modahl C, Green L, Fein D, Morris M, Waterhouse L, Feinstein C, Levin H (1998). Plasma oxytocin levels in autistic children. Biol Psychiatry. 43(4):270–7.

211. Morris CA, Demsey SA, Leonard CO, Dilts C, Blackburn BL. (1998) Natural history of Williams syndrome: physical characteristics. J Pediatr 113:318–26.

212. Mosconi MW, Cody-Hazlett H, Poe MD, Gerig G, Gimpel-Smith R, Piven J (2009). Longitudinal study of amygdala volume and joint attention in 2- to 4-year-old children with autism. Arch Gen Psychiatry. 66(5):509–16.

213. Muñoz KE, Meyer-Lindenberg A, Hariri AR, Mervis CB, Mattay VS, Morris CA, Berman KF (2010). Abnormalities in neural processing of emotional stimuli in Williams syndrome vary according to social vs. non-social content. Neuroimage 50(1):340–6.

214. Murdoch BE (2010). The cerebellum and language: historical perspective and review. Cortex. 2010 Jul-Aug;46(7):858-68.

215. Murphy E, Benítez-Burraco A (2017). Language deficits in schizophrenia and autism as related oscillatory connectomopathies: An evolutionary account. Neurosci Biobehav Rev. 83:742–764.

216. Murphy ER, Foss-Feig J, Kenworthy L, Gaillard WD, Vaidya CJ (2012). Atypical Functional Connectivity of the Amygdala in Childhood Autism Spectrum Disorders during Spontaneous Attention to Eye-Gaze. Autism Res Treat. 2012:652408.

217. Needleman LA, McAllister AK (2012). The major histocompatibility complex and autism spectrum disorder. Dev Neurobiol. 72(10):1288–301.

218. Neira-Fresneda J, Potocki L (2015) Neurodevelopmental disorders associated with abnormal gene dosage: Smith-Magenis and Potocki-Lupski syndromes. J Pediatr Genet. 4(3):159–67. doi: 10.1055/s-0035-1564443.

219. Neumann ID (2002). Involvement of the brain oxytocin system in stress coping: interactions with the hypothalamo-pituitary-adrenal axis. Prog Brain Res. 39:147–62.

220. Newschaffer CJ, Croen LA, Daniels J, Giarelli E, Grether JK, Levy SE, Mandell DS, Miller LA, Pinto-Martin J, Reaven J, Reynolds AM, Rice CE, Schendel D, Windham GC (2007). The epidemiology of autism spectrum disorders. Annu Rev Public Health. 28:235–58.

221. Ng, R., Brown, T. T., Erhart, M., Järvinen, A. M., Korenberg, J. R., Bellugi, U., Halgren, E. (2016). Morphological differences in the mirror neuron system in Williams syndrome. Social neuroscience, 11(3), 277–288.

222. Nickl-Jockschat T, Rottschy C, Thommes J, Schneider F, Laird AR, Fox PT, Eickhoff SB (2015). Neural networks related to dysfunctional face processing in autism spectrum disorder. Brain Struct Funct. 220(4):2355–71.

223. Niego A, Benítez-Burraco A. (2019). Williams Syndrome, Human Self-Domestication, and Language Evolution. Front Psychol. 10:521. doi: 10.3389/fpsyg.2019.00521.

224. Niego A, Kimura R, Benítez-Burraco A (2020) Williams syndrome and autism: dissimilar socio-cognitive profiles with similar patterns of abnormal gene expression in the blood. bioRxiv 2020.03.15.992479; doi: https://doi.org/10.1101/2020.03.15.992479

225. Nummenmaa L, Calder AJ. Neural mechanisms of social attention (2009). Trends Cogn Sci. 13(3):135–43.

226. O’Brien NL, Fiorentino A, Curtis D, Rayner C, Petrosellini C, Al Eissa M, Bass NJ, McQuillin A, Sharp SI (2018) Rare variant analysis in multiply affected families, association studies and functional analysis suggest a role for the ITGΒ4 gene in schizophrenia and bipolar disorder. Schizophr Res. 199:181–188.

227. Oguchi ME, Noguchi K, Fukuda M (2017). TBC1D12 is a novel Rab11-binding protein that modulates neurite outgrowth of PC12 cells. PLoS One. 612(4):e0174883.

228. O’Hearn K., Roth J. K., Courtney S. M., Luna B., Street W., Terwillinger R. (2011). Object recognition in Williams syndrome: uneven ventral stream activation. Dev. Sci. 14 549–565.

229. Osbun N, Li J, O’Driscoll MC, Strominger Z, Wakahiro M, Rider E, Bukshpun P, Boland E, Spurrell CH, Schackwitz W, Pennacchio LA, Dobyns WB, Black GC, Sherr EH (2011). Genetic and functional analyses identify DISC1 as a novel callosal agenesis candidate gene. Am J Med Genet A. 155A(8):1865-76.

230. Osório A, Soares JM, Prieto MF, Vasconcelos C, Fernandes C, Sousa S, Carracedo A, Gonçalves OF, Sampaio A (2014) Cerebral and cerebellar MRI volumes in Williams syndrome. Res Dev Disabil. 35(4):922–8.

231. Palo OM, Antila M, Silander K, Hennah W, Kilpinen H, Soronen P, Tuulio-Henriksson A, Kieseppä T, Partonen T, Lönnqvist J, Peltonen L, Paunio T (2007).Association of distinct allelic haplotypes of DISC1 with psychotic and bipolar spectrum disorders and with underlying cognitive impairments. Hum Mol Genet. 16(20):2517–28.

232. Pannecoeck R, Serruys D, Benmeridja L, Delanghe JR, van Geel N, Speeckaert R Speeckaert MM. (2015). Vascular adhesion protein-1: Role in human pathology and application as a biomarker. Crit Rev Clin Lab Sci. 52(6):284-300.

233. Pankau R, Partsch CJ, Gosch A, Oppermann HC, Wessel A. (1992) Statural growth in Williams-Beuren syndrome. Eur J Pediatr. 151:751–755.

234. Paul LK, Corsello C, Tranel D, Adolphs R (2010). Does bilateral damage to the human amygdala produce autistic symptoms? J Neurodev Disord. 2(3):165–173.

235. Peall KJ, Dijk JM, Saunders-Pullman R, Dreissen YE, van Loon I, Cath D, Kurian MA, Owen MJ, Foncke EM, Morris HR, Gasser T, Bressman S, Asmus F, Tijssen MA (2015). Psychiatric disorders, myoclonus dystonia and SGCE: an international study. Ann Clin Transl Neurol. 3(1):4–11.

236. Peedicayil J, Grayson DR (2018a) An epigenetic basis for an omnigenic model of psychiatric disorders. J Theor Biol. 2018 443:52–55.

237. Peedicayil J, Grayson DR (2018b) Some implications of an epigenetic-based omnigenic model of psychiatric disorders. J Theor Biol. 452:81–84.

238. Pelphrey L, Adolphs R, Morris J (2004). Neuroanatomical substrates of social cognition dysfunction in autism. Mental Retardation and Developmental Disabilities Research Reviews. 10:259–71.

239. Perovic A, Modyanova N, Wexler K (2013). Comparison of Grammar in Neurodevelopmental Disorders: The Case of Binding in Williams Syndrome and Autism With and Without Language Impairment. Lang Acquis. 20(2):133–154.

240. Perovic, A., Wexler, K. (2007). Complex grammar in Williams syndrome. Clinical Linguistics and Phonetics 21: 729–745.

241. Perovic, A., Janke, V. (2013). Issues in the acquisition of binding, control and raising in high-functioning children with autism. UCL Working Papers in Linguistics 25: 131-143.

242. Philofsky A, Fidler DJ, Hepburn S (2007). Pragmatic language profiles of school-age children with autism spectrum disorders and Williams syndrome. Am J Speech Lang Pathol. 16(4):368–80.

243. Phillips, C. E., Jarrold, C., Baddeley, A. D., Grant, J., Karmiloff-Smith, A. (2004). Comprehension of spatial language terms in Williams syndrome: Evidence for an interaction between domains of strength and weakness. Cortex, 40, 85–101.

244. Pinto D, Pagnamenta AT, Klei L, Anney R, Merico D, Regan R, Conroy J, Magalhaes TR, Correia C, Abrahams BS, Almeida J, Bacchelli E, Bader GD, BaileyAJ Baird G, Battaglia A, Berney T, Bolshakova N, Bölte S, Bolton PF, BourgeronT Brennan S, Brian J, Bryson SE, Carson AR, Casallo G, Casey J, Chung BH, Cochrane L, Corsello C, Crawford EL, Crossett A, Cytrynbaum C, Dawson G, de Jonge M, Delorme R, Drmic I, Duketis E, Duque F, Estes A, Farrar P, Fernandez BA, Folstein SE, Fombonne E, Freitag CM, Gilbert J, Gillberg C, Glessner JT, Goldberg J, Green A, Green J, Guter SJ, Hakonarson H, Heron EA, Hill M, Holt R, Howe JL, Hughes G, Hus V, Igliozzi R, Kim C, Klauck SM, Kolevzon A, Korvatska O, Kustanovich V, Lajonchere CM, Lamb JA, Laskawiec M, Leboyer M, Le Couteur A, Leventhal BL, Lionel AC, Liu XQ, Lord C, Lotspeich L, Lund SC, Maestrini E, Mahoney W, Mantoulan C, Marshall CR, McConachie H, McDougle CJ, McGrath J, McMahon WM, Merikangas A, Migita O, Minshew NJ, Mirza GK, Munson J, Nelson SF, Noakes C, Noor A, Nygren G, Oliveira G, Papanikolaou K, Parr JR, Parrini B, PatonT Pickles A, Pilorge M, Piven J, Ponting CP, Posey DJ, Poustka A, Poustka F, Prasad A, Ragoussis J, Renshaw K, Rickaby J, Roberts W, Roeder K, Roge B, Rutter ML, Bierut LJ, Rice JP, Salt J, Sansom K, Sato D, Segurado R, Sequeira AF, SenmanL Shah N, Sheffield VC, Soorya L, Sousa I, Stein O, Sykes N, Stoppioni V, Strawbridge C, Tancredi R, Tansey K, Thiruvahindrapduram B, Thompson AP, Thomson S, Tryfon A, Tsiantis J, Van Engeland H, Vincent JB, Volkmar F, Wallace S, WangK Wang Z, Wassink TH, Webber C, Weksberg R, Wing K, Wittemeyer K, Wood S, Wu J, Yaspan BL, Zurawiecki D, Zwaigenbaum L, Buxbaum JD, Cantor RM, Cook EH, Coon H, Cuccaro ML, Devlin B, Ennis S, Gallagher L, Geschwind DH, Gill M, Haines JL, Hallmayer J, Miller J, Monaco AP, Nurnberger JI Jr Paterson AD, Pericak-Vance MA, Schellenberg GD, Szatmari P, Vicente AM, Vieland VJ, Wijsman EM, Scherer SW, Sutcliffe JS, Betancur C (2010). Functional impact of global rare copy number variation in autism spectrum disorders. Nature. 466(7304):368–72.

245. Pires CF, Rosa FF, Kurochkin I, Pereira CF (2019). Understanding and Modulating Immunity With Cell Reprogramming. Front Immunol. 10:2809.

246. Ponsuksili S, Zebunke M, Murani E (2015). Integrated Genome-wide association and hypothalamus eQTL studies indicate a link between the circadian rhythm-related gene PER1 and coping behavior. Sci Rep. 5:16264.

247. Porter, M.A., Coltheart, M., Langdon, R. (2007). The neuropsychological basis of hypersociability in Williams and Down syndrome. Neuropsychologia, 45, 2839–2849.

248. Preissler, M. A., Carey, S. (2005). The role of inferences about referential intent in word learning: evidence from autism. Cognition 97, B13–B23.

249. Puangpetch A, Suwannarat P, Chamnanphol M, Koomdee N, Ngamsamut N, Limsila P, Sukasem C (2015). Significant Association of HLA-B Alleles and Genotypes in Thai Children with Autism Spectrum Disorders: A Case-Control Study. Dis Markers. 724935.

250. Puche JE, Castilla-Cortázar I (2012) Human conditions of insulin-like growth factor-I (IGF-I) deficiency. J Transl Med. 10:224.

251. Puglia MH, Connelly JJ, Morris JP (2018). Epigenetic regulation of the oxytocin receptor is associated with neural response during selective social attention. Transl Psychiatry. 8(1):116.

252. Puls KL, Ni J, Liu D, Morahan G, Wright MD (1999). The molecular characterisation of a novel tetraspanin protein, TM4-B(1). Biochim Biophys Acta. 1447(1):93-9.

253. Ramírez VT, Ramos-Fernández E, Henríquez JP, Lorenzo A, Inestrosa NC (2016). Wnt-5a/Frizzled9 Receptor Signaling through the Gαo-Gβγ Complex Regulates Dendritic Spine Formation. J Biol Chem. 291(36):19092–107.

254. Raznahan, A., Wallace, G. L., Antezana, L., Greenstein, D., Lenroot, R., Thurm, A. (2013). Compared to what? Early brain overgrowth in autism and the perils of population norms. Biol. Psychiatry 74, 563–75.

255. Reilly, J., Losh, M., Bellugi, U., Wulfeck, B.(2004).“Frog, where are you?” Narratives in children with specific language impairment, early focal brain injury, and Williams syndrome. Brain Lang. 88, 229– 247.

256. Reif A, Schmitt A, Fritzen S, Lesch KP (2007). Neurogenesis and schizophrenia: dividing neurons in a divided mind? Eur Arch Psychiatry Clin Neurosci. 257(5):290–9.

257. Reiss AL, Eckert MA, Rose FE, Karchemskiy A, Kesler S, Chang M, Reynolds MF, Kwon H, Galaburda A (2004). An experiment of nature: brain anatomy parallels cognition and behavior in Williams syndrome. J Neurosci. 24(21):5009–15.

258. Reiss AL, Eliez S, Schmitt JE, Straus E, Lai Z, Jones W, Bellugi U (2000) Neuroanatomy of Williams syndrome: a high-resolution MRI study. J Cogn Neurosci. 12(Suppl 1):65–73.

259. Rhodes, S., Riby, D.M., Park, J., Fraser, E., Campbell, L.E. (2010). Neuropsychological functioning and executive control in WS. Neuropsychologia, 48, 1216–1226.

260. Riby DM, Hancock PJ. (2008) Viewing it differently: social scene perception in Williams syndrome and autism. Neuropsychologia. 46(11):2855–60.

261. Riby DM, Hancock PJ (2009). Do faces capture the attention of individuals with Williams syndrome or autism? Evidence from tracking eye movements. J Autism Dev Disord. 39(3):421–31.

262. Riby DM, Jones N, Brown PH, Robinson LJ, Langton SRH, Bruce V, Riby LM (2011). Attention to faces in Williams syndrome. J Autism Dev Disord 41:1228–1239.

263. Riches, Nick, Tom Loucas, Tony Charman, Emily Simonoff & Gillian Bair (2009). Sentence repetition in adolescents with specific language impairments and autism: An investigation of complex syntax. International Journal of Language and Communication Disorders 45. 47–60.

264. Rice AM, McLysaght A (2017) Dosage-sensitive genes in evolution and disease. BMC Biol. 15(1):78. doi: 10.1186/s12915-017-0418-y.

265. Rizzolatti G, Craighero L (2004). The mirror-neuron system. Annual Review of Neuroscience. 27:169–192.

266. Roberts, Joseph, Mabel Rice Helen Tager-Flusberg. (2004). Tense marking in children with autism. Applied Psycholinguistics 25. 429–448.

267. Rodgers J, Riby DM, Janes E, Connolly B, McConachie H (2012). Anxiety and repetitive behaviours in autism spectrum disorders and williams syndrome: a cross-syndrome comparison. J Autism Dev Disord. 42(2):175–80.

268. Rodríguez-Fornells A, Cunillera T, Mestres-Missé A, de Diego-Balaguer R (2009). Neurophysiological mechanisms involved in language learning in adults. Philos Trans R Soc Lond B Biol Sci. 364(1536):3711–35.

269. Rollins, B., Martin, M. V., Morgan, L., Vawter, M. P. (2010). Analysis of whole genome biomarker expression in blood and brain. American journal of medical genetics. Part B, Neuropsychiatric genetics: the official publication of the International Society of Psychiatric Genetics, 153B(4), 919– 936.

270. Rolls ET (2015). Limbic systems for emotion and for memory, but no single limbicsystem. Cortex. 62:119–57.

271. Rose, F. E., Lincoln, A. J., Lai, Z., Ene, M., Searcy, Y. M., Bellugi, U. (2007). Orientation and affective expression effects on face recognition in Williams syndrome and autism. Journal of Autism and Developmental Disorders, 37, 513–522.

272. Rosen TE, Mazefsky CA, Vasa RA, Lerner MD (2018). Co-occurring psychiatric conditions in autism spectrum disorder. Int Rev Psychiatry. 30(1):40–61.

273. Rosenthal SL, Kamboh MI (2014). Late-Onset Alzheimer’s Disease Genes and the Potentially Implicated Pathways. Curr Genet Med Rep. 2:85–101.

274. Royston R, Howlin P, Waite J, Oliver C (2017). Anxiety Disorders in Williams Syndrome Contrasted with Intellectual Disability and the General Population: A Systematic Review and Meta-Analysis. J Autism Dev Disord. 47(12):3765–3777.1083-1086.

275. Sacco, R., Gabriele, S., and Persico, A. M. (2015). Head circumference and brain size in autism spectrum disorder: A systematic review and meta-analysis. Psychiatry Res. 234, 239–51.

276. Sampaio A, Moreira PS, Osório A, Magalhães R, Vasconcelos C, Férnandez M, Carracedo A, Alegria J, Gonçalves ÓF, Soares JM (2016). Altered functional connectivity of the default mode network in Williams syndrome: a multimodal approach. Dev Sci. 19(4):686–95.

277. Sanders, S. J., Ercan-Sencicek, A. G., Hus, V., Luo, R., Murtha, M. T., Moreno-De-Luca, D., Chu, S. H., Moreau, M. P., Gupta, A. R., Thomson, S. A., Mason, C. E., Bilguvar, K., et al. (2011). Multiple recurrent de novo CNVs, including duplications of the 7q11.23 Williams syndrome region, are strongly associated with autism. Neuron 70: 863–885.

278. Sarret C, Ashkavand Z, Paules E, Dorboz I, Pediaditakis P, Sumner S, Eymard-Pierre E, Francannet C, Krupenko NI, Boespflug-Tanguy O, Krupenko SA (2019). Deleterious mutations in ALDH1L2 suggest a novel cause for neuro-ichthyotic syndrome. NPJ Genom Med. 4:17.

279. Schäffner I, Minakaki G, Khan MA, Balta EA, Schlötzer-Schrehardt U, Schwarz TJ, Beckervordersandforth R, Winner B, Webb AE, DePinho RA, Paik J, Wurst W, Klucken J, Lie DC (2018). FoxO Function Is Essential for Maintenance of Autophagic Flux and Neuronal Morphogenesis in Adult Neurogenesis. Neuron. 99(6):1188–1203.

280. Schuetze M, Park MT, Cho IY, MacMaster FP, Chakravarty MM, Bray SL. (2016) Morphological Alterations in the Thalamus, Striatum, and Pallidum in Autism Spectrum Disorder. Neuropsychopharmacology. 41(11):2627–37.

281. Schmitt JE, Eliez S, Warsofsky IS, Bellugi U, Reiss AL (2001) Enlarged cerebellar vermis in Williams syndrome. J Psychiatr Res. 35(4):225–9.

282. Schultz, R.T. (2005). Developmental deficits in social perception in autism: The role of the amygdala and fusiform face area. International Journal of Developmental Neuroscience, 23, 125–141.

283. Schumann CM, Hamstra J, Goodlin-Jones BL, Lotspeich LJ, Kwon H, Buonocore MH, Lammers CR, Reiss AL, Amaral DG (2004). The amygdala is enlarged in children but not adolescents with autism; the hippocampus is enlarged at all ages. J Neurosci. 24(28):6392–401.

284. Schwaller B, Meyer M, Schiffmann S (2002). ’New’ functions for ‘old’ proteins: the role of the calcium-binding proteins calbindin D-28k, calretinin and parvalbumin, in cerebellar physiology. Studies with knockout mice. Cerebellum. 1 (4): 241–58.

285. Seshadri S, Faust T, Ishizuka K, Delevich K, Chung Y, Kim SH, Cowles M, Niwa M, Jaaro-Peled H, Tomoda T, Lai C, Anton ES, Li B, Sawa A (2015). Interneuronal DISC1 regulates NRG1-ErbB4 signalling and excitatory-inhibitory synapse formation in the mature cortex. Nat Commun. 6:10118.

286. Shang C, Liu Z, Chen Z, Shi Y, Wang Q, Liu S, Li D, Cao P (2015). BRAIN CIRCUITS. A parvalbumin-positive excitatory visual pathway to trigger fear responses in mice. Science. 348(6242):1472–7.

287. Shen, L., Feng, C., Zhang, K., Chen, Y., Gao, Y., Ke, J., Chen, X., Lin, J., Li, C., Iqbal, J., Zhao, Y., Wang, W. (2019). Proteomics Study of Peripheral Blood Mononuclear Cells (PBMCs) in Autistic Children. Frontiers in cellular neuroscience, 13, 105.

288. Shi J, Potash JB, Knowles JA, Weissman MM, Coryell W, Scheftner WA, Lawson WB, DePaulo JR Jr, Gejman PV, Sanders AR, Johnson JK, Adams P, Chaudhury S, Jancic D, Evgrafov O, Zvinyatskovskiy A, Ertman N, Gladis M, Neimanas K, Goodell M, Hale N, Ney N, Verma R, Mirel D, Holmans P, Levinson DF (2011). Genome-wide association study of recurrent early-onset major depressive disorder. Mol Psychiatry. 16(2):193–201.

289. Siegel, D.J., Minshew, N.J., Goldstein, G. (1996). Wechsler IQ profiles in diagnosis of high-functioning autism. Journal of Autism and Developmental Disorders, 26, 389–406.

290. Sigman M., Spence S. J., Wang A. T. (2006). Autism from developmental and neuropsychological perspectives. Annu. Rev. Clin. Psychol. 2, 327–355.

291. Smirnov A, Cappello A, Lena AM, Anemona L, Mauriello A, Di Daniele N, Annicchiarico-Petruzzelli M, Melino G, Candi E (2018). ZNF185 is a p53 target genefollowing DNA damage. Aging (Albany NY). 10(11):3308–3326.

292. Sparaci L, Stefanini S, D’Elia L, Vicari S, Rizzolatti G (2014). What and why understanding in autism spectrum disorders and williams syndrome: similarities and differences. Autism Res. 7(4):421–32.

293. Spratt E. G., Nicholas J. S., Brady K. T., Carpenter L. A., Hatcher C. R., Meekins K. A. (2012). Enhanced cortisol response to stress in children in autism. J. Autism Dev. Disord. 42, 75–81.

294. Stefanacci L., Amaral D. G. (2000). Topographic organization ofcortical inputs to the lateral nucleus of the macaque monkey amygdala: a retrograde tracing study. J. Comp. Neurol. 421 52–7910.1002.

295. Stinton C, Tomlinson K, Estes Z (2012). Examining reports of mental health in adults with Williams syndrome. Res Dev Disabil. 33(1):144–52.

296. Stinton C, Elison S, Howlin P (2010). Mental health problems in adults with Williams syndrome. Am J Intellect Dev Disabil. 115(1):3–18.

297. Stojanovik V, James D (2006). Short-term longitudinal study of a child with Williams syndrome. Int J Lang Commun Disord. 41(2):213–23.

298. Stojanovik V. (2006). Social interaction deficits and conversational inadequacy in Williams syndrome. J Neurolinguistics. 19(2):157–173.

299. Strauss, M. S., Newell, L. C., Giovannelli, J. L., Best, C. A., Minshew, N. J. (2009). The development of facial gender discrimination in typically developing individuals and individuals with autism. Child Development.

300. Su X, Liu X, Ni L, Shi W, Zhu H, Shi J, Chen J, Gu Z, Gao Y, Lan Q, Huang Q (2016). GFAP expression is regulated by Pax3 in brain glioma stem cells. Oncol Rep. 36(3):1277–84.

301. Su AI, Wiltshire T, Batalov S, Lapp H, Ching KA, Block D, Zhang J, Soden R, Hayakawa M, Kreiman G, Cooke MP, Walker JR, Hogenesch JB (2004). A gene atlas of the mouse and human protein-encoding transcriptomes. Proc Natl Acad Sci U S A. 101(16):6062–7.

302. Sudiwala S, Palmer A, Massa V, Burns AJ, Dunlevy LPE, de Castro SCP, Savery D, Leung KY, Copp AJ, Greene NDE (2019). Cellular mechanisms underlying Pax3-related neural tube defects and their prevention by folic acid. Dis Model Mech. 12(11).

303. Sullivan K, Winner E, Tager-Flusberg H (2003). Can adolescents with Williams syndrome tell the difference between lies and jokes? Dev Neuropsychol. 23(1–2):85–103.

304. Sui J., Rotshtein P., Humphreys G. W. (2013). Coupling social attention to the self forms a network for personal significance. Proc. Natl. Acad. Sci. U.S.A. 110, 7607–7612.

305. Surian, L., Baron-Cohen, S., Van derLely, H. (1996). Are children with autism deaf to Gricean maxims? Cogn.Neuropsychiatry 1, 55–71.

306. Swartz JR, Wiggins JL, Carrasco M, Lord C, Monk CS (2013). Amygdala habituation and prefrontal functional connectivity in youth with autism spectrum disorders. J Am Acad Child Adolesc Psychiatry. 52(1):84–93.

307. Swensen LD, Kelley E, Fein D, Naigles LR (2007). Processes of language acquisition in children with autism: evidence from preferential looking. Child Dev. 78:542–557.

308. Tager-Flusberg, H. (2003). Language impairments in children with complex neurodevelopmental disorders: the case of autism. In: Levy, Y., Schaeffer, J.C., editors. Language Competence Across Populations: Toward a Definition of Specific Language Impairment. Mahwah, NJ: Lawrence Erlbaum Associates, pp. 297–321.

309. Tager-Flusberg H (1981). On the nature of linguistic functioning in early infantile autism. J Autism Dev Disord. 11(1):45–56.

310. Tager-Flusberg, H. (2000). Understanding the language and communicative impairments in autism. International Review of Research in Mental Retardation. 23; 185–205.

311. Tager-Flusberg H, Sullivan K (2000). A componential view of theory of mind: evidence from Williams syndrome. Cognition. 76(1):59–90.

312. Tager-Flusberg, H. (2004). Language and communicative deficits and their effects on learning and behavior. In: Prior, M., editor. Asperger Syndrome: Behavioral and Educational Aspects. New York: Guilford Press, pp. 85–103.

313. Tager-Flusberg H, Skwerer DP, Joseph RM (2006). Model syndromes for investigating social cognitive and affective neuroscience: a comparison of Autism and Williams syndrome. Soc Cogn Affect Neurosci 1(3):175–82.

314. Tassabehji M, Read AP, Newton VE, Harris R, Balling R, Gruss P, Strachan T (1992). Waardenburg’s syndrome patients have mutations in the human homologue of the Pax-3 paired box gene. Nature. 355(6361):635–6.

315. Tassabehji M (2003) Williams-Beuren syndrome: a challenge for genotype-phenotype correlations. Hum Mol Genet. 12 Spec No 2:R229-37.

316. Teitelbaum, P., Teitelbaum, O., Nye, J., Fryman, J., Maurer, R.G. (1998). Movement analysis in infancy may be useful for early diagnosis of autism. Proceedings of the National Academy of Sciences of the United States of America, 95, 13982–13987.

317. Tek S, Jaffery G, Fein D, Naigles L (2008). Do children with autism spectrum disorders show a shape bias in word learning? Autism Res. 1:208–222.

318. Telang U, Morris ME (2010). Effect of orally administered phenethyl isothiocyanate on hepatic gene expression in rats. Mol Nutr Food Res. 54(12):1802–6.

319. Trauner D. A., Bellugi U. Chase C. (1989) Neurologic features of Williams and Down syndromes. Pediatric Neurology 5, 166–8.

320. Thakur, D., Martens, M. A., Smith, D. S., Roth, E. (2018). Williams Syndrome and Music: A Systematic Integrative Review. Frontiers in Psychology, 9, 2203.

321. Thomson, P. A., Parla, J. S., McRae, A. F., Kramer, M., Ramakrishnan, K., Yao, J., Soares, D. C., McCarthy, S., Morris, S. W., Cardone, L., Cass, S., Ghiban, E., Hennah, W., Evans, K. L., Rebolini, D., Millar, J. K., Harris, S. E., Starr, J. M., MacIntyre, D. J., Generation Scotland, Porteous, D. J. (2014). 708 Common and 2010 rare DISC1 locus variants identified in 1542 subjects: analysis for association with psychiatric disorder and cognitive traits. Molecular psychiatry, 19(6), 668–675.

322. Thompson PM, Lee AD, Dutton RA, Geaga JA, Hayashi KM, Eckert MA, Bellugi U, Galaburda AM, Korenberg R, Mills DL, Toga AW, Reiss AL. (2005). Abnormal cortical complexity and thickness profiles mapped in Williams syndrome. J Neurosci. 25(16):4146-58.

323. Thornton-Wells, T. A., Avery, S. N., Blackford, J. U. (2011). Using novel control groups to dissect the amygdala’s role in Williams syndrome. Developmental cognitive neuroscience, 1(3), 295–304.

324. Tordjman S, Anderson GM, Botbol M, Toutain A, Sarda P, Carlier M, Saugier-Veber P, Baumann C, Cohen D, Lagneaux C, Tabet AC, Verloes A. (2012). Autistic disorder in patients with Williams-Beuren syndrome: a reconsideration of the Williams-Beuren syndrome phenotype. PLoS One. 7(3):e30778.

325. Tomaiuolo F, Di Paola M, Caravale B, Vicari S, Petrides M, Caltagirone C. (2002) Morphology and morphometry of the corpus callosum in Williams syndrome: a T1-weighted MRI study. Neuroreport. 13:2281–2284

326. Tomasi, D., Volkow, N. D. (2019). Reduced Local and Increased Long-Range Functional Connectivity of the Thalamus in Autism Spectrum Disorder. Cerebral cortex (New York, N.Y.: 1991), 29(2), 573– 585.

327. Tong F, Zhang M, Guo X, Shi H, Li L, Guan W, Wang H, Yang S (2016). Expression patterns of SH3BGR family members in zebrafish development. Dev Genes Evol. 226(4):287–95.

328. Toyoshima, D., Mandai, K., Maruo, T., Supriyanto, I., Togashi, H., Inoue, T., Mori, M. and Takai, Y. (2014). Afadin regulates puncta adherentia junction formation and presynaptic differentiation in hippocampal neurons. PloS one, 9(2), e89763.

329. Tropea D, Hardingham N, Millar K, Fox K (2018). Mechanisms underlying the role of DISC1 in synaptic plasticity. J Physiol. 596(14):2747–2771.

330. Tyra M, Ropka-Molik K, Piórkowska K, Oczkowicz M, Szyndler-Nędza M, Małopolska M (2019). Association of Ghrelin Gene Polymorphisms with Fattening Traits and Feed Intake in Pig: A Preliminary Study. Animals (Basel). 9(7).

331. Uddin LQ, Supekar K, Lynch CJ, Khouzam A, Phillips J, Feinstein C (2013). Salience network–based classification and prediction of symptom severity in children with autism. JAMA Psychiatry. 70:869– 79.

332. Udwin, O., Yule, W. (1990). Expressive language of children with Williams syndrome. American Journal of Medical Genetics, 6(Suppl.), 108–14.

333. Van Daalen E, Swinkels SH, Dietz C, van Engeland H, Buitelaar JK (2007). Body length and head growth in the first year of life in autism. Pediatr Neurol. 37(5):324–30.

334. Valenta T, Lukas J, Doubravska L, Fafilek B, Korinek V (2006). HIC1 attenuates Wnt signaling by recruitment of TCF-4 and beta-catenin to the nuclear bodies. EMBO J. 25(11):2326–37.

335. Van Leeuwen JE, Rafalovich I, Sellers K, Jones KA, Griffith TN, Huda R, Miller RJ, Srivastava DP, Penzes P (2014). Coordinated nuclear and synaptic shuttling of afadin promotes spine plasticity and histone modifications. J Biol Chem. 289(15):10831–42.

336. Vannucchi G, Masi G, Toni C, Dell’Osso L, Marazziti D, Perugi G (2014). Clinical features, developmental course, and psychiatric comorbidity of adult autism spectrum disorders. CNS Spectr. 19(2):157–64.

337. Van Overwalle F, Baetens K (2009). Understanding others’ actions and goals by mirror and mentalizing systems: a meta-analysis. Neuroimage. 48:564–584.

338. Vargas DM, De Bastiani MA, Zimmer ER, Klamt F (2018). Alzheimer’s disease master regulators analysis: search for potential molecular targets and drug repositioning candidates. Alzheimers Res Ther. 10(1):59.

339. Verheyen GR, Villafuerte SM, Del-Favero J, Souery D, Mendlewicz J, Van Broeckhoven C, Raeymaekers P (1999). Genetic refinement and physical mapping of a chromosome 18q candidate region for bipolar disorder. Eur J Hum Genet. 7(4):427–34.

340. Vias C, Dick AS (2017). Cerebellar Contributions to Language in Typical and Atypical Development: A Review. Dev Neuropsychol. 42(6):404–421.

341. Viñas-Guasch N, Wu YJ (2017) The role of the putamen in language: a meta-analytic connectivity modeling study. Brain Struct Funct. 222(9):3991–4004.

342. Vivanti G, Hocking DR, Fanning P, Dissanayake C (2016). Social affiliation motives modulate spontaneous learning in Williams syndrome but not in autism. Mol Autism. 7(1):40.

343. Vithayathil J, Pucilowska J, Landreth GE (2018). ERK/MAPK signaling and autism spectrum disorders. Prog Brain Res. 241:63–112.

344. Walenski, M., Tager-Flusberg, H., Ullman, M.T. (2006). Language in autism. In: Moldin, S., Rubenstein, J., editors. Understanding Autism: From Basic Neuroscience to Treatment. New York: Taylor & Francis, pp. 175–203.

345. Wahl, M., Marzinzik, F., Friederici, A. D., Hahne, A., Kupsch, A., Schneider, G. H., Saddy, D., Curio, G., and Klostermann, F. (2008). The human thalamus processes syntactic and semantic language violations. Neuron 59, 695–707.

346. Wallace S, Guo DC, Regalado E, Mellor-Crummey L, Bamshad M, Nickerson DA, Dauser R, Hanchard N, Marom R, Martin E, Berka V, Sharina I, Ganesan V, Saunders D, Morris SA, Milewicz DM (2016). Disrupted nitric oxide signaling due to GUCY1A3 mutations increases risk for Moyamoya disease, achalasia and hypertension. Clin Genet. 90(4):351–60.

347. Wallace GL, Shaw P, Lee NR, Clasen LS, Raznahan A, Lenroot RK, Giedd JN (2012). Distinct cortical correlates of autistic versus antisocial personality traits in a longitudinal sample of typically developing youth. Journal of Neuroscience. 32:4856–4860.

348. Walker RM, Hill AE, Newman AC, Hamilton G, Torrance HS, Anderson SM, Ogawa F, Derizioti P, Nicod J, Vernes SC, Fisher SE, Thomson PA, Porteous DJ, Evans KL (2012). The DISC1 promoter: characterization and regulation by FOXP2. Hum Mol Genet. 21(13):2862–72.

349. Wang M, Zhou J, He F, Cai C, Wang H, Wang Y, Lin Y, Rong H, Cheng G, Xu R, Zhou W (2019). Alteration of gut microbiota-associated epitopes in children with autism spectrum disorders. Brain Behav. Immun. 75:192–199.

350. Wang X, Wang X, Zhang S, Sun H, Li S, Ding H, You Y, Zhang X, Ye SD (2019). The transcription factor TFCP2L1 induces expression of distinct target genes and promotes self-renewal of mouse and human embryonic stem cells. J Biol Chem. 294(15):6007–6016.

351. Wang S, Liang Q, Qiao H, Li H, Shen T, Ji F, Jiao J (2016). Development. 143: 2732–2740.

352. Wei H, Alberts I, Li X. (2014). The apoptotic perspective of autism. Int J Dev Neurosci. 36:13–8.

353. White, S. W., Albano, A. M., Johnson, C. R., Kasari, C., Ollendick, T., Klin, A. (2010). Development of a cognitive behavioural intervention program to treat anxiety and social deficits in teens with high functioning autism. Clinical Child Family Psychology Review, 13, 77–90.

354. Witt SH, Streit F, Jungkunz M, Frank J, Awasthi S, Reinbold CS, Treutlein J, Degenhardt F, Forstner AJ, Heilmann-Heimbach S, Dietl L, Schwarze CE, Schendel D, Strohmaier J, Abdellaoui A, Adolfsson R, Air TM, Akil H, Alda M, Alliey-RodriguezN Andreassen OA, Babadjanova G, Bass NJ, Bauer M, Baune BT, Bellivier F, Bergen S, Bethell A, Biernacka JM, Blackwood DHR, Boks MP, Boomsma DI, Børglum AD, Borrmann-Hassenbach M, Brennan P, Budde M, Buttenschøn HN, Byrne EM, Cervantes P, Clarke TK, Craddock N, Cruceanu C, Curtis D, Czerski PM, Dannlowski U, Davis T, de Geus EJC, Di Florio A, Djurovic S, Domenici E, Edenberg HJ, Etain B, FischerSB, Forty L, Fraser C, Frye MA, Fullerton JM, Gade K, Gershon ES, Giegling I, Gordon SD, Gordon-Smith K, Grabe HJ, Green EK, Greenwood TA, Grigoroiu-Serbanescu M, Guzman-Parra J, Hall LS, Hamshere M, Hauser J, Hautzinger M, Heilbronner U, Herms S, Hitturlingappa S, Hoffmann P, Holmans P, Hottenga JJ, Jamain S, Jones I, Jones LA, Juréus A, Kahn RS, Kammerer-Ciernioch J, Kirov G, Kittel-Schneider S, Kloiber S, Knott SV, Kogevinas M, Landén M, Leber M, Leboyer M, Li QS, Lissowska J, Lucae S, Martin NG, Mayoral-Cleries F, McElroy SL, McIntosh AM, McKay JD, McQuillin A, Medland SE, Middeldorp CM, Milaneschi Y, Mitchell PB, Montgomery GW, Morken G, Mors O, Mühleisen TW, Müller-Myhsok B, Myers RM, Nievergelt CM, Nurnberger JI, O’Donovan MC, Loohuis LMO, Ophoff R, Oruc L, Owen MJ, Paciga SA, Penninx BWJH, Perry A, Pfennig A, Potash JB, Preisig M, Reif A, Rivas F, Rouleau GA, Schofield PR, Schulze TG, Schwarz M, Scott L, Sinnamon GCB, Stahl EA, StraussJ Turecki G, Van der Auwera S, Vedder H, Vincent JB, Willemsen G, Witt CC, Wray NR, Xi HS (2017). Genome-wide associationstudy of borderline personality disorder reveals genetic overlap with bipolardisorder, major depression and schizophrenia. Transl Psychiatry. 7(6):e1155.

355. Witt SH, Sommer WH, Hansson AC, Sticht C, Rietschel M, Witt CC (2013). Comparison of gene expression profiles in the blood, hippocampus and prefrontal cortex of rats. In Silico Pharmacol. 1:15.

356. Wójciak P, Remlinger-Molenda A, Rybakowski J (2012). The role of oxytocin and vasopressin in central nervous system activity and mental disorders. Psychiatr Pol. 46(6):1043–52.

357. Wolters FJ, Boender J, de Vries PS, Sonneveld MA, Koudstaal PJ, de Maat MP, Franco OH, Ikram MK, Leebeek FW, Ikram MA. (2018). Von Willebrand factor and ADAMTS13 activity in relation to risk of dementia: a population-based study. Sci Rep. 8(1):5474.

358. Woodward ND, Giraldo-Chica M, Rogers B, Cascio CJ (2017). Thalamocortical dysconnectivity in autism spectrum disorder: An analysis of the Autism Brain Imaging Data Exchange. Biol Psychiatry Cogn Neurosci Neuroimaging. 2(1):76–84.

359. Xiao, J., Vemula, S. R., Xue, Y., Khan, M. M., Carlisle, F. A., Waite, A. J., Blake, D. J., Dragatsis, I., Zhao, Y., LeDoux, M. S. (2017). Role of major and brain-specific Sgce isoforms in the pathogenesis of myoclonus-dystonia syndrome. Neurobiology of disease, 98, 52–65.

360. Xie Z, Yang X, Deng X, Ma M, Shu K (2017). A Genome-Wide Association Study and Complex Network Identify Four Core Hub Genes in Bipolar Disorder. Int J Mol Sci 18(12).

361. Yang X, Ru W, Wang B, Gao X, Yang L, Li S, Xi S, Gong P (2016). Investigating the genetic basis of attention to facial expressions: the role of the norepinephrine transporter gene. Psychiatr Genet. 26(6):266–271.

362. Yao Z, Darowski K, St-Denis N, Wong V, Offensperger F, Villedieu A, Amin S, Malty R, Aoki H, Guo H, Xu Y, Iorio C, Kotlyar M, Emili A, Jurisica I, Neel BG, Babu M, Gingras AC, Stagljar I. (2017). A Global Analysis of the Receptor Tyrosine Kinase-Protein Phosphatase Interactome. Mol Cell. 65(2):347–360.

363. Zalla, Tiziana, and Marco Sperduti (2013). The amygdala and the relevance detection theory of autism: an evolutionary perspective. Frontiers in human neuroscience vol. 7 894.

364. Zarrei M, Burton CL, Engchuan W, Young EJ, Higginbotham EJ, MacDonald JR, Trost B, Chan AJS, Walker S, Lamoureux S, Heung T, Mojarad BA, Kellam B, Paton T, Faheem M, Miron K, Lu C, Wang T, Samler K, Wang X, Costain G, Hoang N, Pellecchia G, Wei J, Patel RV, Thiruvahindrapuram B, Roifman M, Merico D, Goodale T, Drmic I, Speevak M, Howe JL, Yuen RKC, Buchanan JA, Vorstman JAS, Marshall CR, Wintle RF, Rosenberg DR, Hanna GL, Woodbury-Smith M, Cytrynbaum C, Zwaigenbaum L, Elsabbagh M, Flanagan J, Fernandez BA, Carter MT, Szatmari P, Roberts W, Lerch J, Liu X, Nicolson R, Georgiades S, Weksberg R, Arnold PD, Bassett AS, Crosbie J, Schachar R, Stavropoulos DJ, Anagnostou E, Scherer SW (2019). A large data resource of genomic copy number variation across neurodevelopmental disorders. NPJ Genom Med. 4:26.

365. Zhang J, Guo W, Tian B, Sun M, Li H, Zhou L, Liu X. (2015). Puerarin attenuates cognitive dysfunction and oxidative stress in vascular dementia rats induced by chronic ischemia. Int J Clin Exp Pathol. 8(5):4695–704.

366. Zhang, H. F., Dai, Y. C., Wu, J., Jia, M. X., Zhang, J. S., Shou, X. J., Han, S. P., Zhang, R., Han, J. S. (2016). Plasma Oxytocin and Arginine-Vasopressin Levels in Children with Autism Spectrum Disorder in China: Associations with Symptoms. Neuroscience bulletin, 32(5), 423–432.

367. Zhang Y, Yuan X, Wang Z, Li R (2014). The canonical Wnt signaling pathway in autism. CNS Neurol Disord Drug Targets. 13(5):765–70.

368. Zhao C, Avilés C, Abel RA, Almli CR, McQuillen P, Pleasure SJ. (2005). Hippocampal and visuospatial learning defects in mice with a deletion of frizzled 9, a gene in the Williams syndrome deletion interval. Development 132(12):2917–27.

369. Zhou Y, Qiu L, Sterpka A, Wang H, Chu F, Chen X (2019). Comparative Phosphoproteomic Profiling of Type III Adenylyl Cyclase Knockout and Control, Male, and Female Mice. Front Cell Neurosci. 13:34

370. Ziegler C, Grundner-Culemann F, Schiele MA, Schlosser P, Kollert L, Mahr M, Gajewska A, Lesch KP, Deckert J, Köttgen A, Domschke K (2019).The DNA methylome in panic disorder: a case-control and longitudinal psychotherapy-epigenetic study. Transl Psychiatry. 9(1):314.

