## Supplemental file 2 for "Autism and Williams syndrome: dissimilar socio-cognitive profiles with similar patterns of abnormal gene expression in the blood"

**Functional characterization of genes found dysregulated in the blood of subjects with ASD or WS and with a potential impact on the etiopathogenesis of these conditions.**

**A. Genes upregulated in both ASD and WS**

*GPR171*

This gene encodes a protein related to the G protein coupled receptor, which acts as a receptor for the neuropeptide bigLEN as part of the system regulating anxiety and fear responses, and which is expressed in the basolateral amygdala (Bobeck et al. 2017).

*SGCE*.

This gene encodes a member of the sarcoglycan family and causes myoclonus dystonia, a hyperkinetic movement disorder frequently associated to social phobia (Peall et al., 2015), as well as anxiety and obsessive compulsive disorder (Saunders–Pullman et al., 2002; van Tricht et al., 2012).

**B. Genes downregulated genes in both ASD and WS**

*DSEL*

This gene encodes a protein involved in the metabolism of chondroitin sulfate/dermatan and glycosaminoglycan which is expressed in the brain and which has been associated to bipolar disorder (Verheyen et al., 1999) and depression (Shi et al., 2011).

*EPHB1*

This gene, encodes a receptor for ephrin-B family members. A polymorphism of this gene has been associated to attentive behavior to faces (Yang et al., 2016). *EphB1* knocked-out mice exhibit aberrant thalamic-cortical axon guidance (Robichaux et al., 2014), which, as discussed in the main text (Results, section 2), is a usual finding in both ASD and WS.

*FBXL13*

This gene encodes a protein ubiquitin ligase associated with the Class 1 MHC mediated antigen processing and presentation and has been highlighted as one core gene involved in bipolar disorder (Xie et al., 2017).

*FOXO4*

This gene encodes a fork-head transcription factor targeted by USP7, a candidate for ASD and for neurodevelopmental disorders involving speech delay and altered behavior (Fountain et al., 2019). *FOXO4* is involved dendrite development, with *FOXO4* deficiency resulting in altered dendritic morphology and abnormal spine density and positioning (Schaffner et al., 2018). As discussed below, dendrite abnormalities are regularly associated to both ASD and WS.

*ITGB4*

This gene encodes one component of a receptor for laminin, an extracellular matrix protein involved in cellular interaction, motility and signaling (Tarone et al., 2000). The gene has been associated to schizophrenia and bipolar disorder (O’Brien et al., 2018). Interestingly, ITGB4 binds NRG1 as part as Neuregulin-ERBB signaling, which plays a key role in neuronal migration and differentiation, synaptogenesis, neural myelination, neurotransmission, and synaptic plasticity (Mei and Nave, 2014; Kataria et al., 2019). Interestingly too, neuregulins interact with *LIMK1*, one of the genes deleted in WS, which encodes a protein kinase involved in visuospatial cognition (Wang et al., 1998). One important neuregulin, namely NRG1, is seemingly involved in social behavior, as Nrg1(+/-) mice show reduced social activity, mimicking aspects of the social dysfunctional behaviour observed in subjects with ASD (Ehrlichman et al., 2009).

*PKP4*

This gene encodes a protein involved in cellular adhesion, motility and division, as well as neurite outgrowth (Keil et al., 2013). The gene has been implicated in the sexual dimorphism observed in ASD (Zhou et al., 2019), but it has been associated as well to bipolar disorder, major depression, and schizophrenia (Witt et al., 2017).

*PVALB*

This gene encodes parvalbumin. The gene is expressed in GABAergic interneurons in many brain regions of interest for the ASD and WS pathogenesis, including the thalamus, the cortex, and the basal ganglia (Schwaller et al., 2002). *PVALB* has been directly related to the ASD phenotype. Accordingly, reduced transcription of *PVALB* caused by the happloinsufficiency of the ASD-candidate *ARID1B* has been hypothesized to contribute to the cognitive and social deficits observed in subjects with this condition (Jung et al., 2017). Likewise, *PVALB* expression is dysregulated in mice knocked-out for *Cntnap2*, a robust candidate for ASD (Lauber et al., 2018). Interestingly, activation of selected paravalbumin pathways is involved in triggering fear responses and induced conditioned aversion (Shang et al., 2015), which are abnormal in people with ASD or WS.

*VIL1*

This gene encodes an actin-binding protein and it has been highlighted as a risk factor for ASD (De Rubeis et al., 2014).

**C. Genes upregulated in ASD but downregulated in WS**

*CCL4L1*

This gene encodes a cytokine seemingly involved in inflammatory and immunoregulatory processes (Colobran et al., 2010). The gene has been identified as a risk factor for panic disorder, with patients exhibiting increased methylation of one intron of the gene compared to controls (Ziegler et al., 2019). Whereas panic disorder is not usually described in people with WS, it has been reported in subjects with ASD, who score high in the anxiety subscale of panic (Gillott et al., 2007). Panic behaviors by individuals with ASD have also been found to be associated with a specific allele of the *OTXR* gene, which encodes the oxytocin receptor (de Oliveira Pereira Ribeiro et al., 2018).

*CLEC9A*

This gene is involved in cell reprogramming into dendritic cells (Pires et al., 2019), but also in the response to dying cells, as it triggers endocytosis (Schulz et al., 2002; Sancho et al., 2009).

*NME8*

This gene encodes a protein involved in ciliary dynamics and it has been associated with Alzheimer’s disease and specifically, with the abnormal cognitive processes involved in the disease. Accordingly, an *NME8*-adjacent SNP (rs2718058) seems to play a preventive role in this condition, reducing brain neurodegeneration, as well as brain atrophy and brain hypometabolism (Liu et al., 2014). Likewise, one particular SNP (rs12155159) of *NME8* has been associated to less cognitive decline on a domain-specific cognitive test score (the delayed word recall test) in people with Alzheimer’s disease (Bressler et al., 2017).

**D. Genes downregulated in WS but upregulated ASD**

*DSC1*

This gene encodes a member of the cadherin superfamily which is involved in the formation of desmosomes, as well as in cell adhesion and signal transduction (Wang et al., 2016). The gene is expressed in the mouse corpus callosum (Miyata et al., 2015). People with WS exhibit a shorter corpus callosum with altered morphology compared to controls, and this circumstance has been hypothesized to account for some of their cognitive impairments (Tomaiuolo et al., 2002; Sampaio et al. 2013). Decreased volumes and signs of less myelination of the corpus callosum have been observed as well in people with ASD, although contradictory results can be found in the literature, suggesting that within-group differences in callosal morphology may be related to within-group differences in cognitive performance (see Travers et al., 2012 for review and discussion).

**References**

Bobeck EN, Gomes I, Pena D, Cummings KA, Clem RL, Mezei M, Devi LA (2017). The BigLEN-GPR171 Peptide Receptor System Within the Basolateral Amygdala Regulates Anxiety-Like Behavior and Contextual Fear Conditioning. Neuropsychopharmacology.

42(13):2527-2536.

Bressler J, Mosley TH, Penman A, Gottesman RF, Windham BG, Knopman DS, Wruck LM, Boerwinkle E. (2017). Genetic variants associated with risk of Alzheimer's disease contribute to cognitive change in midlife: The Atherosclerosis Risk in Communities Study. Am J Med Genet B Neuropsychiatr Genet. 174(3):269-282.

Colobran R, Pedrosa E, Carretero-Iglesia L, Juan M. (2010). Copy number variation in chemokine superfamily: the complex scene of CCL3L-CCL4L genes in health and disease. Clin Exp Immunol. 162(1):41-52.

de Oliveira Pereira Ribeiro L, Vargas-Pinilla P, Kappel DB, Longo D, Ranzan J, Becker MM, Dos Santos Riesgo R, Schuler-Faccini L, Roman T, Schuch JB (2018). Evidence for Association Between OXTR Gene and ASD Clinical Phenotypes. J Mol Neurosci. 65(2):213-221.

De Rubeis S, He X, Goldberg AP, Poultney CS, Samocha K, Cicek AE, Kou Y, LiuL, Fromer M, Walker S, Singh T, Klei L, Kosmicki J, Shih-Chen F, Aleksic B,Biscaldi M, Bolton PF, Brownfeld JM, Cai J, Campbell NG, Carracedo A, ChahrourMH, Chiocchetti AG, Coon H, Crawford EL, Curran SR, Dawson G, Duketis E,Fernandez BA, Gallagher L, Geller E, Guter SJ, Hill RS, Ionita-Laza J, JimenzGonzalez P, Kilpinen H, Klauck SM, Kolevzon A, Lee I, Lei I, Lei J, Lehtimäki T, Lin CF, Ma'ayan A, Marshall CR, McInnes AL, Neale B, Owen MJ, Ozaki N, Parellada M, Parr JR, Purcell S, Puura K, Rajagopalan D, Rehnström K, Reichenberg A, SaboA, Sachse M, Sanders SJ, Schafer C, Schulte-Rüther M, Skuse D, Stevens C,Szatmari P, Tammimies K, Valladares O, Voran A, Li-San W, Weiss LA, Willsey AJ,Yu TW, Yuen RK; DDD Study; Homozygosity Mapping Collaborative for Autism; UK10KConsortium, Cook EH, Freitag CM, Gill M, Hultman CM, Lehner T, Palotie A,Schellenberg GD, Sklar P, State MW, Sutcliffe JS, Walsh CA, Scherer SW, Zwick ME,Barett JC, Cutler DJ, Roeder K, Devlin B, Daly MJ, Buxbaum JD (2014). Synaptic,transcriptional and chromatin genes disrupted in autism. Nature. 515(7526):209-15.

Ehrlichman RS, Luminais SN, White SL, Rudnick ND, Ma N, Dow HC, Kreibich AS, Abel T, Brodkin ES, Hahn CG, Siegel SJ (2009). Neuregulin 1 transgenic mice display reduced mismatch negativity, contextual fear conditioning and social interactions. Brain Res. 1294:116-27.

Fountain MD, Oleson DS, Rech ME, Segebrecht L, Hunter JV, McCarthy JM, Lupo PJ, Holtgrewe M, Moran R, Rosenfeld JA, Isidor B, Le Caignec C, Saenz MS, Pedersen RC, Morgan TM, Pfotenhauer JP, Xia F, Bi W, Kang SL, Patel A, Krantz ID, Raible SE, Smith W, Cristian I, Torti E, Juusola J, Millan F, Wentzensen IM, Person RE, Küry S, Bézieau S, Uguen K, Férec C, Munnich A, van Haelst M, Lichtenbelt KD, van Gassen K, Hagelstrom T, Chawla A, Perry DL, Taft RJ, Jones M, Masser-Frye D, Dyment D, Venkateswaran S, Li C, Escobar LF, Horn D, Spillmann RC, Peña L, Wierzba J, Strom TM, Parenti I, Kaiser FJ, Ehmke N, Schaaf CP (2019). Pathogenic variants in USP7 cause a neurodevelopmental disorder with speech delays, altered behavior, and neurologic anomalies. Genet Med. 21(8):1797-1807.

Gillott A, Standen PJ (2007). Levels of anxiety and sources of stress in adults with autism. J Intellect Disabil. 11(4):359-70.

Jung EM, Moffat JJ, Liu J, Dravid SM, Gurumurthy CB, Kim WY (2017). Arid1b haploinsufficiency disrupts cortical interneuron development and mouse behavior. Nat Neurosci. 20(12):1694-1707.

Kataria N, Martinez CA, Kerr B, Zaiter SS, Morgan M, McAlpine SR, Cook KM (2019). C-Terminal HSP90 Inhibitors Block the HIF-1 Hypoxic Response by Degrading HIF-1α through the Oxygen-Dependent Degradation Pathway. Cell Physiol Biochem. 53(3):480-495.

Keil R, Schulz J, Hatzfeld M (2013). p0071/PKP4, a multifunctional protein coordinating cell adhesion with cytoskeletal organization. Biol Chem. 394(8):1005-17

Lauber E, Filice F, Schwaller B (2018). Dysregulation of Parvalbumin Expression in the Cntnap2-/- Mouse Model of Autism Spectrum Disorder. Front Mol Neurosci. 2;11:262.

Liu Y, Yu JT, Wang HF, Hao XK, Yang YF, Jiang T, Zhu XC, Cao L, Zhang DQ, Tan L (2014). Association between NME8 locus polymorphism and cognitive decline,cerebrospinal fluid and neuroimaging biomarkers in Alzheimer's disease. PLoS One.8;9(12):e114777.

Mei L, Nave KA (2014). Neuregulin-ERBB signaling in the nervous system and neuropsychiatric diseases. Neuron. 83(1):27-49.

Miyata S, Yoshikawa K, Taniguchi M, Ishikawa T, Tanaka T, Shimizu S, Tohyama M (2015). Sgk1 regulates desmoglein 1 expression levels in oligodendrocytes in the mouse corpus callosum after chronic stress exposure. Biochem Biophys Res Commun. 464(1):76-82.

O'Brien NL, Fiorentino A, Curtis D, Rayner C, Petrosellini C, Al Eissa M, Bass NJ, McQuillin A, Sharp SI. (2018). Rare variant analysis in multiply affected families, association studies and functional analysis suggest a role for the ITGΒ4 gene in schizophrenia and bipolar disorder. Schizophr Res. 199:181-188.

Peall KJ, Dijk JM, Saunders-Pullman R, Dreissen YE, van Loon I, Cath D, Kurian MA, Owen MJ, Foncke EM, Morris HR, Gasser T, Bressman S, Asmus F, Tijssen MA (2015). Psychiatric disorders, myoclonus dystonia and SGCE: an international study. Ann Clin Transl Neurol. 3(1):4-11.

Pires CF, Rosa FF, Kurochkin I, Pereira CF (2019). Understanding and Modulating Immunity With Cell Reprogramming. Front Immunol. 10:2809.

Robichaux MA, Chenaux G, Ho HY, Soskis MJ, Dravis C, Kwan KY, Šestan N, Greenberg ME, Henkemeyer M, Cowan CW (2014). EphB receptor forward signaling regulates area-specific reciprocal thalamic and cortical axon pathfinding. Proc Natl Acad Sci U S A. 111(6):2188-93.

Sampaio A, Bouix S, Sousa N, Vasconcelos C, Férnandez M, Shenton ME, Gonçalves ÓF (2013). Morphometry of corpus callosum in Williams syndrome: shape as an index of neural development. Brain Struct Funct. 218(3):711-20.

Sancho D, Joffre OP, Keller AM, Rogers NC, Martínez D, Hernanz-Falcón P, Rosewell I, Reis e Sousa C (2009). Identification of a dendritic cell receptor that couples sensing of necrosis to immunity. Nature. 458(7240):899-903.

Saunders-Pullman R, Ozelius L, Bressman SB (2002). Inherited myoclonus-dystonia. Adv Neurol. 89:185-91.

Schäffner I, Minakaki G, Khan MA, Balta EA, Schlötzer-Schrehardt U, Schwarz TJ, Beckervordersandforth R, Winner B, Webb AE, DePinho RA, Paik J, Wurst W,Klucken J, Lie DC (2018). FoxO Function Is Essential for Maintenance of Autophagic Flux and Neuronal Morphogenesis in Adult Neurogenesis. Neuron. 99(6):1188-1203.

Schulz O, Reis e Sousa C (2002). Cross-presentation of cell-associated antigens by CD8alpha+ dendritic cells is attributable to their ability to internalize dead cells. Immunology. 107(2):183-9.

Schwaller B, Meyer M, Schiffmann S (2002). "'New' functions for 'old' proteins: the role of the calcium-binding proteins calbindin D-28k, calretinin and parvalbumin, in cerebellar physiology. Studies with knockout mice". Cerebellum. 1 (4): 241–58.

Shang C, Liu Z, Chen Z, Shi Y, Wang Q, Liu S, Li D, Cao P (2015).BRAIN CIRCUITS. A parvalbumin-positive excitatory visual pathway to trigger fear responses in mice.Science. 348(6242):1472-7.

Shi J, Potash JB, Knowles JA, Weissman MM, Coryell W, Scheftner WA, Lawson WB,DePaulo JR Jr, Gejman PV, Sanders AR, Johnson JK, Adams P, Chaudhury S, Jancic D,Evgrafov O, Zvinyatskovskiy A, Ertman N, Gladis M, Neimanas K, Goodell M, Hale N,Ney N, Verma R, Mirel D, Holmans P, Levinson DF (2011). Genome-wide association study of recurrent early-onset major depressive disorder. Mol Psychiatry. 16(2):193-201.

Tarone G, Hirsch E, Brancaccio M, De Acetis M, Barberis L, Balzac F, Retta SF, Botta C, Altruda F, Silengo L (2000). Integrin function and regulation in development. Int J Dev Biol. 44(6):725-31.

Tomaiuolo F, Di Paola M, Caravale B, Vicari S, Petrides M, Caltagirone C (2002). Morphology and morphometry of the corpus callosum in Williams syndrome: a T1-weighted MRI study. Neuroreport. 13(17):2281-4.

Travers BG, Adluru N, Ennis C, Tromp do PM, Destiche D, Doran S, Bigler ED, Lange N, Lainhart JE, Alexander AL (2012). Diffusion tensor imaging in autism spectrum disorder: a review. Autism Res. 5(5):289-313.

van Tricht MJ, Dreissen YE, Cath D, Dijk JM, Contarino MF, van der Salm SM, Foncke EM, Groen JL, Schmand B, Tijssen MA (2012). Cognition and psychopathology in myoclonus-dystonia. J Neurol Neurosurg Psychiatry. 83(8):814-20.

Verheyen GR, Villafuerte SM, Del-Favero J, Souery D, Mendlewicz J, Van Broeckhoven C, Raeymaekers P (1999). Genetic refinement and physical mapping of a chromosome 18q candidate region for bipolar disorder. Eur J Hum Genet. 7(4):427-34.

Wang Y, Chen C, Wang X, Jin F, Liu Y, Liu H, Li T, Fu J (2016). Lower DSC1 expression is related to the poor differentiation and prognosis of head and neck squamous cell carcinoma (HNSCC). J Cancer Res Clin Oncol. 142(12):2461-2468.
